# Mitochondrial complex III-derived ROS amplify immunometabolic changes in astrocytes and promote dementia pathology

**DOI:** 10.1101/2024.08.19.608708

**Authors:** Daniel Barnett, Till S. Zimmer, Caroline Booraem, Fernando Palaguachi, Samantha M. Meadows, Haopeng Xiao, Edward T. Chouchani, Anna G. Orr, Adam L. Orr

## Abstract

Neurodegenerative disorders alter mitochondrial functions, including the production of reactive oxygen species (ROS). Mitochondrial complex III (CIII) generates ROS implicated in redox signaling, but its triggers, targets, and disease relevance are not clear. Using site-selective suppressors and genetic manipulations together with mitochondrial ROS imaging and multiomic profiling, we found that CIII is the dominant source of ROS production in astrocytes exposed to neuropathology-related stimuli. Astrocytic CIII-ROS production was dependent on nuclear factor-κB (NF-κB) and the mitochondrial sodium-calcium exchanger (NCLX) and caused oxidation of select cysteines within immune and metabolism-associated proteins linked to neurological disease. CIII-ROS amplified metabolomic and pathology-associated transcriptional changes in astrocytes, with STAT3 activity as a major mediator, and facilitated neuronal toxicity in a non-cell-autonomous manner. As proof-of-concept, suppression of CIII-ROS in mice decreased dementia-linked tauopathy and neuroimmune cascades and extended lifespan. Our findings establish CIII-ROS as an important immunometabolic signal transducer and tractable therapeutic target in neurodegenerative disease.

## Main

Mitochondria couple metabolism and respiration to energy production, but also produce ROS via partial reduction of oxygen. Mitochondria are key contributors to total ROS levels and their impairment or cell stress can dramatically increase ROS production.^1^ Mitochondria generate ROS (mtROS) from at least 11 distinct sites, with respiratory complex I (CI) and complex III (CIII) considered the major sources and each linked to distinct signaling and disease states.^2–8^ CIII has a high capacity for ROS production due in part to its ubiquitous and high expression.^9–11^ Moreover, CIII is topologically poised to influence intracellular signaling by generating ROS away from the inner mitochondrial matrix unlike CI and most other sites.^9^

The exact regulators and effects of site-specific mtROS production in health and disease remain undefined due to long-standing challenges, including off-target effects of most genetic and pharmacologic manipulations of mtROS and the direct dependence of mtROS on metabolism. To address these challenges, we previously identified potent, small-molecule <u>S</u>uppressors of <u>E</u>lectron <u>L</u>eak (SELs), including <u>S</u>uppressors of Complex <u>III_Qo_ E</u>lectron <u>L</u>eak (S3QELs or “sequels”) and <u>S</u>uppressors of Complex <u>I_Q_ E</u>lectron <u>L</u>eak (S1QELs or “cycles”) which selectively block ROS production from a single mtROS site without altering normal mitochondrial functions like ATP production.^12,13^ SELs are efficacious in diverse systems, including cultured cells, invertebrates, and mice,^7,12–19^ and uniquely enable the dissociation of site-specific mtROS production from other metabolic processes.

In the brain, mitochondrial dysfunction is an early and prevalent feature of neurodegenerative disorders,^20–22^ and mtROS are implicated as central, feed-forward drivers of cell dysfunction and neuropathology.^23–29^ Notably, a dominantly-inherited mutation in the CIII subunit UQCRC1 is associated with parkinsonism and polyneuropathy and linked to increased basal ROS production in human neural cells.^30^ Similarly, mice with brain-specific deficiency in RISP, the CIII subunit necessary for carrying electrons away from the ROS production site, have increased oxidative stress and early mortality, suggesting that increased CIII-ROS is sufficient to cause neuropathology.^31^ Indeed, the main contributing factors to many neurodegenerative disorders, including aging, amyloid-β (Aβ) accumulation, tauopathy, and neuroinflammation, are linked to mitochondrial dysfunction and increased mtROS, suggesting a central role for mitochondrial oxidative mechanisms in disease.^32–38^ In support, cognitive resilience to dementia-related pathology is associated with lower expression of proteins linked to oxidative stress.^39^ Yet, despite mounting evidence implicating mtROS in disease, the exact triggers, sources, and downstream oxidation targets of mtROS and their contributions to cell signaling and disease mechanisms remain unclear.

Here, we used SELs along with multiple other pharmacological and genetic manipulations to establish that astrocytic CIII-ROS induction by dementia-linked stimuli reinforces specific transcriptional programs and neuroimmune cascades. CIII-ROS emerge as key pathogenic signal transducers which can be safely and selectively suppressed to alleviate disease-associated changes in mitochondrial redox signaling, glial responses, and neuroimmune cascades.

## Results

### Select stimuli trigger astrocytic complex III-derived ROS through NF-κB and NCLX activity

Astrocytes are essential to proper CNS function and involved in dementia-linked pathogenesis and neuroimmune cascades.^40–44^ Astrocytes have a high capacity for mtROS production, including from CI and CIII,^13,45^ and manipulation of astrocytic redox balance and intracellular ROS influences brain metabolism, neuronal function, and cognition,^42,46–49^ and might promote neuropathology.^50–52^ To investigate the induction and regulation of astrocytic ROS in response to pathogenic stimuli, we first measured the rates of cellular H_2_O_2_ efflux in primary mouse astrocytes (Extended Data Fig. 1a,b) upon stimulation with neuroimmune factors induced in neurological disease.^53–58^ The cytokines interleukin-6 (IL-6) and interferon-γ (IFN-γ) had no effects on H_2_O_2_ efflux, whereas a cocktail of cytokines linked to astrocyte-dependent neurotoxicity, consisting of IL-1α, tumor necrosis factor-α (TNFα), and complement component 1q (C1q),^53^ increased the rates of astrocytic H_2_O_2_ production by 24 h (Fig. 1a). The cocktail-induced H_2_O_2_ production was attributable to IL-1α, but not to TNFα or C1q (Fig. 1a), suggesting that astrocytic ROS is engaged by specific types of ligands and signaling mechanisms. To test the contribution of CIII-ROS in astrocyte H_2_O_2_ efflux, we treated astrocytes with IL-1α and one of two structurally distinct S3QELs (Fig. 1b) to rule out common off-target effects. S3QEL1.2 and S3QEL2 inhibited IL-1α-induced H_2_O_2_ production to a similar extent (54.7% and 56.3%) (Fig. 1c), suggesting that CIII is the dominant source of IL-1α-induced increases in H_2_O_2_ levels. To rule out off-target metabolic effects of SELs, ATP levels were measured in astrocytes metabolically restricted to pyruvate- and galactose-dependent respiration. As reported in other cell types,^7,12,13,59^ S3QELs and S1QELs did not affect mitochondrial metabolism (Extended Data Fig. 1c,d).

**Figure 1.**
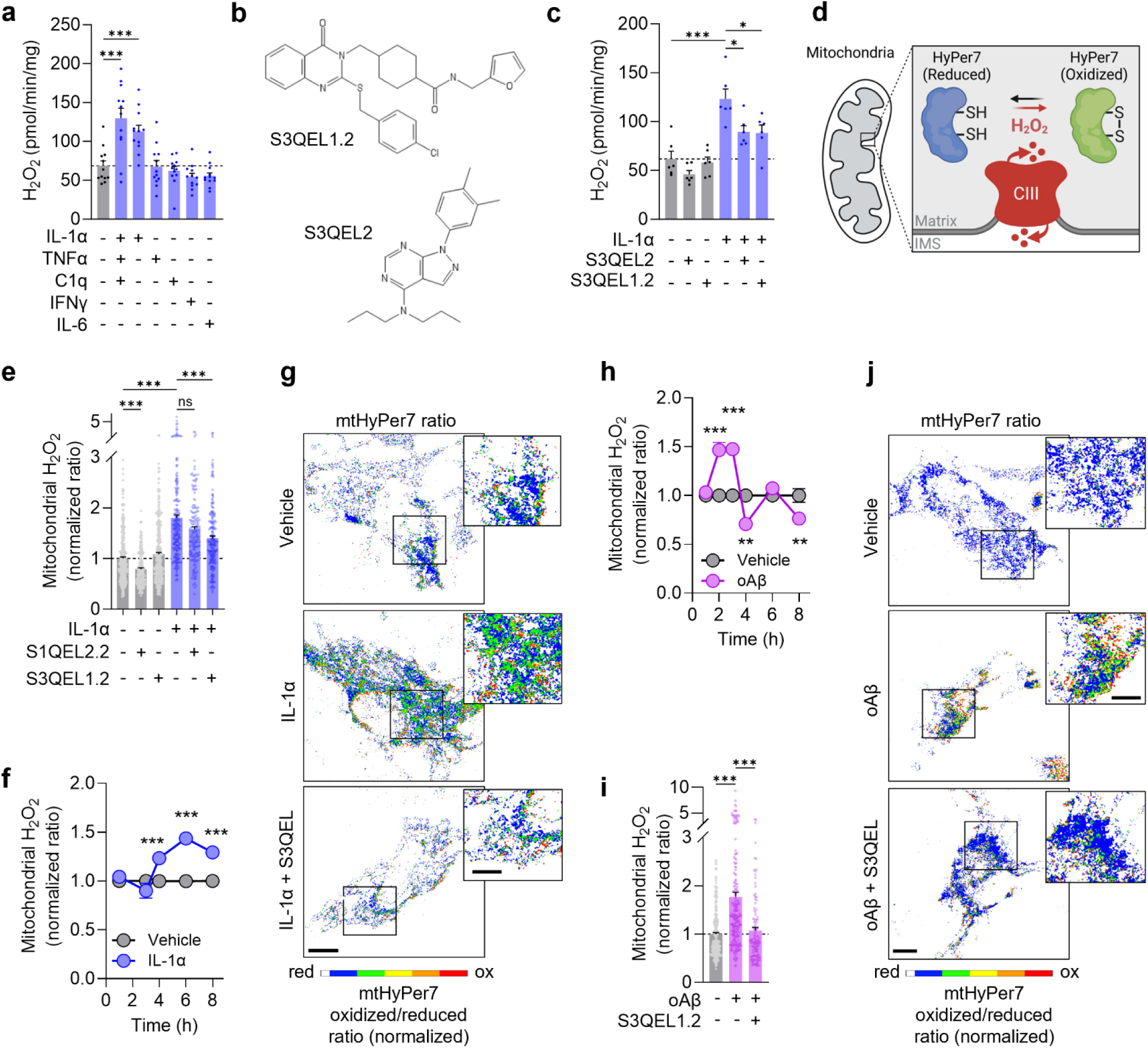
Astrocytic CIII-ROS are induced by specific stimuli. a–c, Cellular H_2_O_2_ efflux after 24-h treatment with vehicle, IL-1α (3 ng/mL), TNFα (30 ng/mL), C1q (400 ng/mL), IL-6 (33 ng/mL), IFN-γ (10 ng/mL) with or without S3QEL2 (20 μM) or S3QEL1.2 (1 µM). n=6–12 wells, ANOVA with Dunnett’s (**a**) or Tukey’s test (**c**). **d**, Schematic of mtHyPer7 sensing mitochondrial CIII-derived H_2_O_2_. **e**, Measurement of mtHyPer7 fluorescence 6 h after treatment with vehicle, IL-1α, S3QEL1.2 (3 μM), or S1QEL2.2 (10 μM). n=162–297 cells, Kruskal Wallace with Dunn’s test. **f**, Time-course of mtHyPer7 fluorescence after vehicle or IL-1α. n=56–296 cells, Mann-Whitney, unpaired two-tailed t-test. **g**, Representative images of mtHyPer7 oxidized/reduced ratios after 6-h treatment with vehicle, IL-1α, or IL-1α with S3QEL1.2. Scale bars: 10 µm; 5 µm (inset). **h**, Time-course of mtHyPer7 fluorescence after vehicle or oAβ (300 nM). n=54–310 cells, Mann-Whitney, unpaired two-tailed t-test. **i**, Quantification of mtHyPer7 after 3-h treatment with vehicle, oAβ, or S3QEL1.2 (3 μM). n=110–233 cells, Kruskal Wallace with Dunn’s test. **j**, Representative images of mtHyPer7 oxidized/reduced ratios after 3-h treatment with vehicle, oAβ, or oAβ with S3QEL1.2. Scale bars: 10 µm, 5 µm (inset). Data are shown as mean ± SEM. *p<0.05, **p<0.01, ***p<0.001.

To further investigate the mechanisms regulating CIII-ROS, we targeted the highly sensitive and selective ratiometric H_2_O_2_ sensor HyPer7^60^ to mitochondria (mtHyPer7; Fig. 1d and Extended Data Fig. 1e,f) and performed live-cell confocal imaging. We confirmed that mtHyPer7 is oxidized in response to treatment with antimycin A, a potent inducer of CIII-ROS production.^61,62^ Antimycin A-induced oxidation was dose-dependently suppressed by S3QEL1.2 (Extended Data Fig. 1g,h), indicating that mtHyPer7 enables precise and spatially resolved measurements of astrocytic CIII-ROS.

Live-cell imaging of mtHyPer7 revealed that S3QEL1.2 had no effects on baseline H_2_O_2_ levels, whereas the CI-ROS suppressor S1QEL2.2 reduced baseline mitochondrial H_2_O_2_ levels (Fig. 1e), suggesting that CI-ROS plays a greater role than CIII-ROS in setting the basal redox tone in astrocyte mitochondria, in line with previous reports.^13,45^ To better understand the temporal dynamics of mtROS induction, we performed confocal imaging of mtHyPer7 at different timepoints after stimulation with IL-1α. Mitochondrial H_2_O_2_ was increased within 4 h, peaked by 6 h, and remained elevated until at least 8 h (Fig. 1f). In contrast to its lack of effect on basal mtHyPer7 oxidation, S3QEL1.2 inhibited IL-1α-induced responses by approx. 49% (Fig. 1e,g). Notably, CI-ROS suppressor S1QEL2.2 and the NOX-ROS inhibitor APX-115 had no effects (Fig. 1e and Extended Data Fig. 1i). These results suggest that a major component of IL-1α-induced mtROS is derived from CIII, but not from CI or NOX enzymes.

In addition to IL-1α, we examined the effects of oAβ, which is implicated in Alzheimer’s disease and may cause pathology in part through its effects on astrocyte function and increasing intracellular ROS levels.^22,63–66^ Indeed, we found that oAβ caused dynamic changes in astrocytic mtROS levels, increasing mtROS within 2 h, which remained elevated until 3 h after stimulation (Fig. 1h). The onset and recovery of oAβ effects were faster and more dynamic as compared to IL-1α effects (Figs. 1f, h), possibly due to the multiple targets and intracellular pathways engaged by oAβ and its dynamic aggregation states. Importantly, treatment with S3QEL1.2 fully blocked this induction (Fig. 1i,j), revealing a pronounced role of CIII in oAβ- induced mtROS responses. Unlike IL-1α and oAβ, stimulation with IL-6 did not increase astrocytic mtROS across multiple timepoints (Extended Data Fig. 1j), confirming its lack of effect on cellular H_2_O_2_ efflux (Fig. 1a) and further highlighting the stimulus-specificity of mtROS induction.

To further investigate the involvement of CIII in IL-1α and oAβ-induced ROS production, we generated mice that express alternative oxidase (AOX) selectively in astrocytes. AOX is a mitochondrial enzyme not normally expressed in mammals, but when ectopically expressed, it localizes to the inner mitochondrial membrane and consumes oxygen through the oxidation of ubiquinol, the substrate required for CIII-ROS production (Fig. 2a).^67^ By shunting ubiquinol, AOX suppresses CIII-ROS^68^ but is also reported to mitigate ubiquinol-dependent ROS production from CI. However, since blockade of astrocytic CIII-ROS and not CI-ROS affects cytokine-induced mtROS in astrocytes (Fig. 1e), ectopic expression of AOX functionally complements the pharmacological suppression of CIII-ROS during cell stimulation.

**Figure 2.**
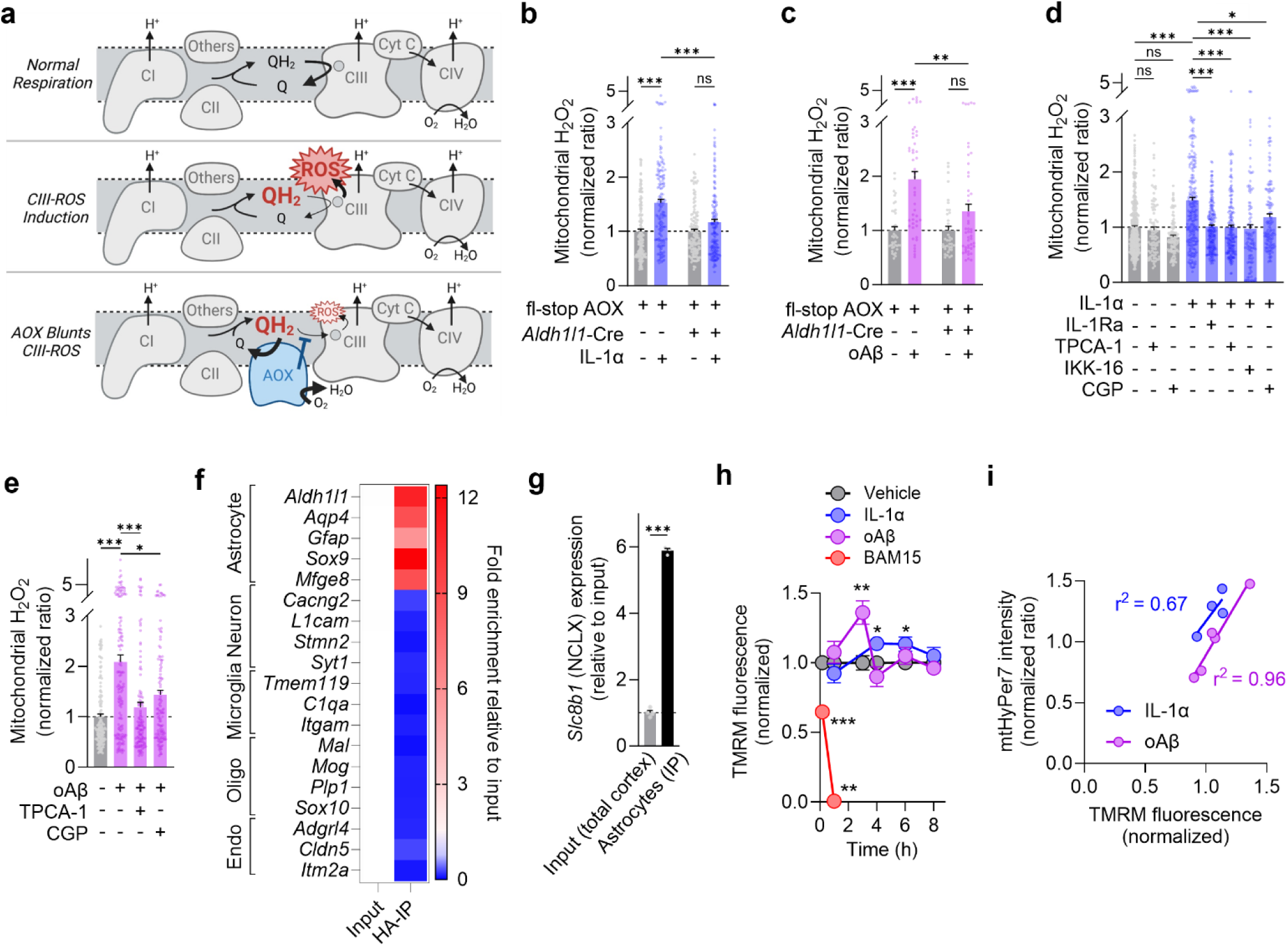
NF-κB and NCLX regulate astrocytic CIII-ROS induction. a, Schematic depicting electron flow to the full reduction of O_2_ to H_2_O at complex IV (top panel) or the partial reduction to superoxide (ROS) at CIII (middle panel). Ectopically expressed alternative oxidase (AOX) blunts CIII-ROS by consuming ubiquinol (QH_2_) and O_2_ without producing ROS (bottom panel). **b**,**c**, Quantification of mtHyPer7 in *Aldh1l1*-Cre/fl-stop AOX or control fl-stop AOX astrocytes treated with vehicle, IL-1α (6 h), or oAβ (3 h). n=37–161 cells, Kruskal-Wallis with Dunn’s test. **d,e**, Quantification of mtHyPer7 after IL-1α (6 h) or oAβ (3 h) co-treatment with vehicle, IL-1Ra (1 μg/mL), TPCA-1 (1 μM), IKK-16 (3 μM) or CGP 37157 (CGP; 10 μM). Inhibitors were added 1 h prior and again at the start of stimulation with IL-1α or oAβ. n=84– 257 cells, Kruskal-Wallis with Dunn’s test. **f,g**, Gene expression of cell type-specific markers and *Slc8b1* (NCLX) in total and ribosomal-captured mRNA in cortical samples from *Aldh1l1*-Cre/HA-RiboTag^fl/fl^ mice. n=3 mice, unpaired two-tailed t-test. **h**, Time-dependent changes in mitochondrial membrane potential after treatment with vehicle, IL-1α, oAβ, or BAM15 as a positive control. n=3–21 wells, unpaired two-tailed t-test. (**i**) Correlational analysis of mtHyPer7 versus TMRM changes after treatment with IL-1α or oAβ as compared to vehicle. Data are shown as mean ± SEM. *p<0.05, **p<0.01, **p<0.001.

To verify AOX function, we evaluated its ability to alleviate metabolic deficits induced by dysfunction or inhibition of respiratory complexes III and IV, but not complex I.^6,7,67–71^ As expected (Extended Data Fig. 2a), inhibition of electron transport at CIII with myxothiazol alone, or together with inhibition of CI with rotenone, effectively blocked complex IV-dependent mitochondrial respiration in fl-stop-AOX control astrocytes (Extended Data Fig. 2b). In contrast, double transgenic *Aldh1l1*-AOX astrocytes maintained substantial oxygen consumption in the presence of myxothiazol, which increases ubiquinol for AOX activation, but not in the presence of rotenone, which lowers ubiquinol (Extended Data Fig. 2b). These results indicate that AOX was functional and activated by ubiquinol oxidation. Notably, AOX expression did not affect basal mtROS or ATP levels, suggesting that AOX does not alter mitochondrial energy production or redox state in astrocytes at baseline (Extended Data Fig. 2c,d).

Consistent with our findings that CIII-ROS is triggered by IL-1α and oAβ, *Aldh1l1*-AOX astrocytes had reduced mtHyPer7 responses to IL-1α and oAβ as compared to control fl-stop-AOX astrocytes (reduced by 68.2% and 62.3%, respectively) (Fig. 2b,c). To test whether functional AOX is required for lowering mtROS responses, we assessed wild-type astrocytes acutely transfected with functional or a catalytically-inactive mutant AOX^68,72^ and confirmed that functional AOX but not mutant AOX effectively suppressed mtROS responses to antimycin A, IL-1α, and oAβ (Extended Data Fig. 2e–g). Together, these results suggest that CIII is a major driver of astrocytic mtROS responses to disease-related factors.

To identify the molecular mechanisms required for the induction of CIII-ROS, we first tested the involvement of nuclear factor-κB (NF-κB), a transcription factor (TF) activated by IL-1R signaling and associated with astrocytic responses to diverse stimuli.^73–75^ Indeed, IL-1α rapidly increased NF-κB phosphorylation and this increase preceded CIII-ROS induction and was not affected by S3QEL1.2 (Extended Data Fig. 2h). Importantly, inhibition of NF-κB activation with TPCA-1 or IKK-16 or preventing IL- 1R activation with IL-1R antagonist protein (IL-1Ra) potently suppressed mtROS induction by IL-1α and oAβ (Fig. 2d,e), indicating that NF-κB activation is necessary for the induction of CIII-ROS. Inhibition of NF-κB in the absence of stimuli had no effect on basal mtROS (Fig. 2d).

To further explore specific mechanisms within mitochondria which promote CIII-ROS production in astrocytes, we first examined whether IL-1α altered the expression of electron transport chain components. However, subunits of complexes I–V were unchanged following stimulation (Extended Data Fig. 2i). Next, we tested the expression and activity of the mitochondrial sodium-calcium exchanger (NCLX), an inner-membrane protein enriched in astrocytes (Fig. 2f,g) that regulates mitochondrial ion homeostasis and was suggested to promote CIII-ROS in peripheral cells under hypoxia.^76^ Although IL-1α did not increase RNA levels of NCLX (Extended Data Fig. 2j), blockade of NCLX ion flux with the potent inhibitor CGP 37157^77–79^ markedly decreased mtROS induction by IL-1α and oAβ (Fig. 2d,e), suggesting that NCLX activity mediates stimuli-induced production of CIII-ROS in astrocytes. Inhibition of NCLX had minimal effects on basal mtROS (Fig. 2d). Notably, mitochondrial membrane potential (ΔΨm) is a strong driver of NCLX activity.^80,81^ In agreement, we found that IL-1α and oAβ caused hyperpolarization of mitochondria in astrocytes across time scales closely matching those of mtROS induction (Fig. 2h). Treatment with BAM15, a mitochondria-selective uncoupler, was used as a positive control to depolarize mitochondria.^82^ Additionally, there were strong positive correlations between ΔΨm and mtROS alterations upon IL-1α and oAβ treatments (Fig. 2i). These results suggest that ΔΨm hyperpolarization and the associated increases in NCLX activity promote CIII-ROS induction in an NF-κB-dependent manner.

### CIII-ROS induce oxidation of diverse but specific target proteins linked to disease

CIII-ROS is proposed to be a major contributor to redox signaling but its exact oxidation targets are largely unknown.^11,83,84^ Previous studies in isolated mitochondria have suggested that different sources of ROS (e.g., CIII vs. CI) oxidize unique mitochondrial targets.^85^ However, it is not known if CIII or any other source of mtROS causes oxidation of select targets in intact cells and whether these putative oxidation patterns are affected by physiological stimuli.

To profile disease-relevant CIII-specific oxidation targets across the entire astrocytic proteome under physiologically relevant conditions, we performed unbiased stoichiometric cysteine-redox proteomics in astrocytes stimulated with IL-1α with or without S3QEL1.2 co-treatment and compared to vehicle as control (Fig. 3a).^86^ We identified and quantified the percent reversible oxidation of a total of 8,263 unique cysteine-containing peptides (cys-peptides), with nearly identical overall ion intensity, cysteine abundance, and coverage across the proteome between conditions (Fig. 3b,c).

**Figure 3.**
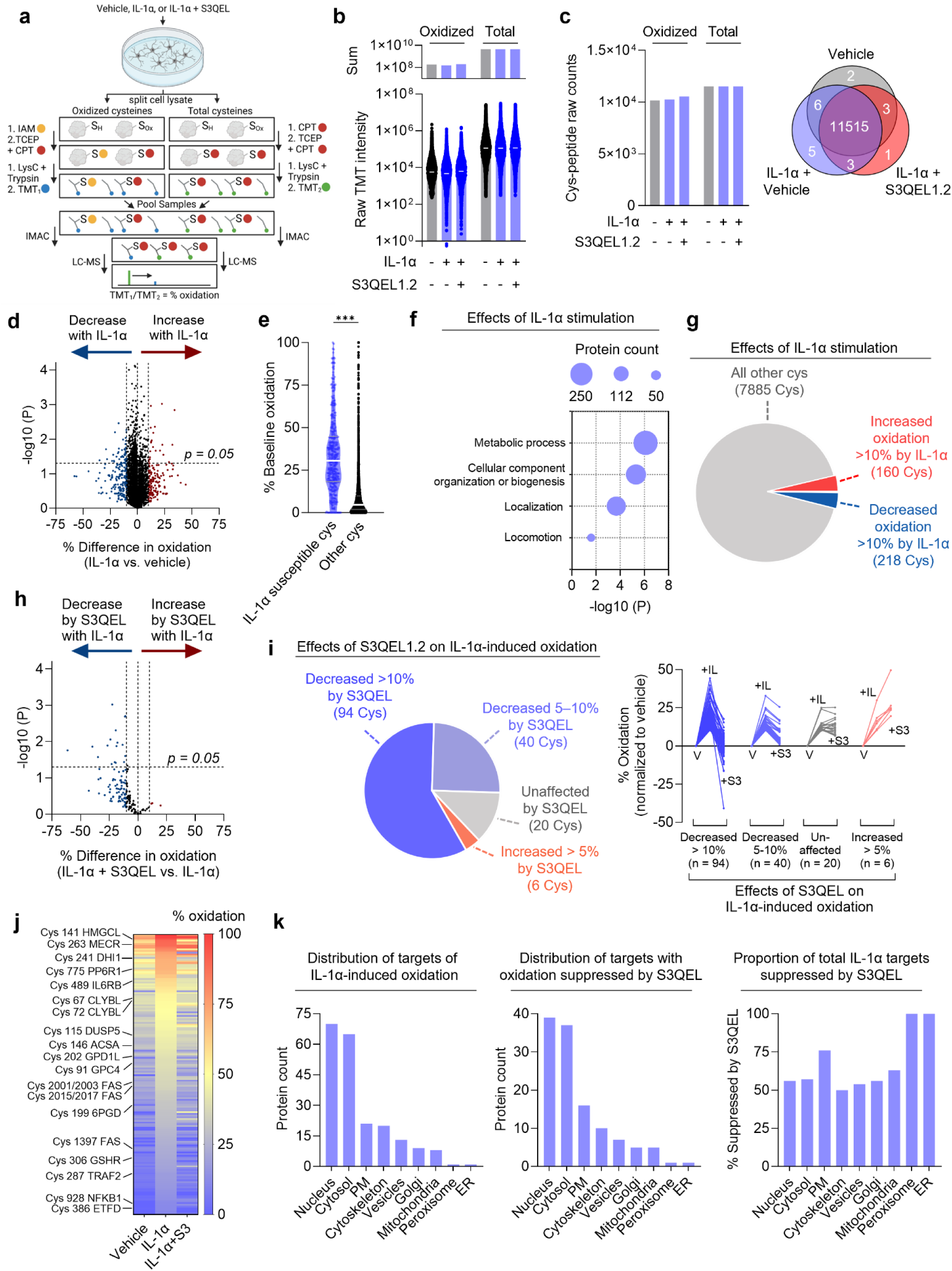
CIII-ROS oxidize disease-relevant cysteines across the proteome. a, Schematic outlining the method of stoichiometric redox proteomics applied to astrocytes after 6-h treatment with vehicle, IL-1α, or IL-1α with S3QEL1.2 (3 μM). Adapted from ref^86^. **b**, Raw and summed TMT ion intensities averaged across replicates for each condition. **c**, Raw cysteine (cys)-peptide counts averaged across replicates for each condition and Venn diagram of all cys-peptides detected in each condition. **d**, Volcano plot of 8,263 identified cys-peptides across all samples. Maroon and blue circles highlight individual cysteines whose oxidation increased or decreased by 10% or more after IL-1α treatment relative to vehicle. **e**, Comparison of baseline oxidation of cys-residues susceptible or not susceptible to IL-1α-dependent modification. Unpaired two-tailed t-test: ***p<0.001. **f**, Gene ontology enrichment analysis for cys-peptides oxidized ≥ 10% by IL-1α. Significance threshold was set as p<0.05 following Benjamini-Hochberg correction for multiple testing. **g**, Cys-peptides grouped according to IL-1α-induced effects. **h**, Volcano plot showing IL- 1α-induced oxidized cys-peptides modified by co-treatment with S3QEL1.2. **i**, Proportions and plots of IL- 1α-induced oxidized cys-peptides based on effect of S3QEL1.2. **j**, Heatmap of the effects of IL-1α with or without S3QEL1.2 on redox states of 160 cys-peptides oxidized ≥ 10% after IL-1α treatment. Specific redox-active cys-residues in proteins linked to immune, metabolic, and neuropathogenic processes are highlighted. **k**, Subcellular distribution of redox-active proteins influenced by IL-1α and S3QEL1.2.

Although IL-1α substantially increased astrocytic mtROS levels (as shown in Fig. 1), we did not detect global increases in oxidation states in the cysteine proteome. Instead, less than 4% of mitochondrial cys-peptides detected (25 out of 718) and 5% of total cys-peptides detected (378 out of 8,263) changed their baseline oxidation by 10% or more at the time of maximal mtROS induction by IL-1α (Fig. 3d; also see Fig. 1f), a cutoff consistent with previous reports on meaningful effects of cysteine modifications.^86,87^ Notably, our profiling shows that cysteines that were susceptible to IL-1α-induced oxidation had higher baseline oxidation levels (median of 34% oxidized) as compared to cysteines in the total proteome (median of 8% oxidized) (Fig. 3e), suggesting that this subpopulation of cysteines in the astrocyte proteome is on average more redox-active at baseline and responsive to cell stimulation, possibly due to their specific intracellular microenvironment and redox properties. Proteins susceptible to IL-1α-induced oxidation were highly enriched for metabolic processes, indicating that IL-1α-mediated redox signaling may be an intracellular mechanism to modulate metabolic functions (Fig. 3f).

Consistent with CIII-ROS promoting oxidative changes, S3QEL1.2 inhibited IL-1α-induced oxidation of the astrocytic proteome. Among the 160 cys-peptides (from 150 unique proteins) that were oxidized 10% or more after IL-1α (Fig. 3g), 59% were decreased at least 10% by S3QEL and 84% were decreased at least 5% by S3QEL (Fig. 3h,i). Top CIII-ROS-sensitive cysteines included proteins linked to immune pathways (NFKB1, PP6R1, IL6RB, TRAF2), lipid metabolism (FAS, ACSA, ETFD, HMGCL, MECR, CLYBL), dementia-related pathways (GPC4), and redox regulation (GSHR, AMPL) (Fig. 3j). Of note, our results validate metabolic proteins previously reported to be targets of antimycin A-induced ROS in isolated mitochondria (e.g., ETFD, CLYBL)^85^ and more specifically identify the individual cysteines in these proteins sensitive to CIII-ROS.

In support of an intracellular signaling role for CIII-ROS beyond the mitochondria, S3QEL-sensitive cysteines were found across multiple intracellular compartments, suggesting that CIII-ROS can diffuse to oxidize extra-mitochondrial targets or engage in long-distance relays to promote cysteine modifications at multiple distinct intracellular sites (Fig. 3k). Overall, these results indicate that CIII-ROS are major contributors to increased protein oxidation upon IL-1α stimulation and reveal CIII-ROS as a previously unrecognized source of redox-based modifications within disease-associated proteins. To our knowledge, these findings are also the first example of wide-ranging protein oxidation across multiple subcellular compartments caused by mtROS and that a single mitochondrial site induced by a disease-relevant factor can promote these effects.

Next, we performed polar metabolomic profiling to determine whether IL-1α promotes CIII-ROS by altering metabolic pathways or whether CIII-ROS are acting upstream of metabolic changes. We found that IL-1α stimulation of astrocytes increased the levels of several metabolites (Extended Data Fig. 3a) including the tricarboxylic acid cycle intermediates malate and fumarate as well as argininosuccinate, which can generate fumarate and participate in the urea cycle (Extended Data Fig. 3b-d). Inhibition of NF-κB with TPCA-1 prevented many of the IL-1α-induced metabolic changes (Extended Data Fig. 3e). Moreover, treatment with S3QEL1.2 also suppressed these changes (Extended Data Fig. 3f), indicating that NF-κB- induced CIII-ROS production is promoting this metabolic rewiring.

### CIII-ROS promote astrocytic STAT3 activity and gene expression linked to neural deficits

Gene expression changes are integral to astrocytic responses in various neurological disorders,^42,88^ but whether mtROS influence astrocytic transcriptional activities is not clear. To investigate if induction of CIII- ROS regulates specific transcriptional programs, we performed untargeted RNA sequencing in IL-1α- stimulated astrocytes in the presence or absence of S3QEL1.2. TF enrichment analyses confirmed that NF- κB is a central mediator of IL-1α-induced changes in astrocytic gene expression (Fig. 4a), consistent with the observed NF-κB-dependence of CIII-ROS induction (Fig. 2d,e).

**Figure 4.**
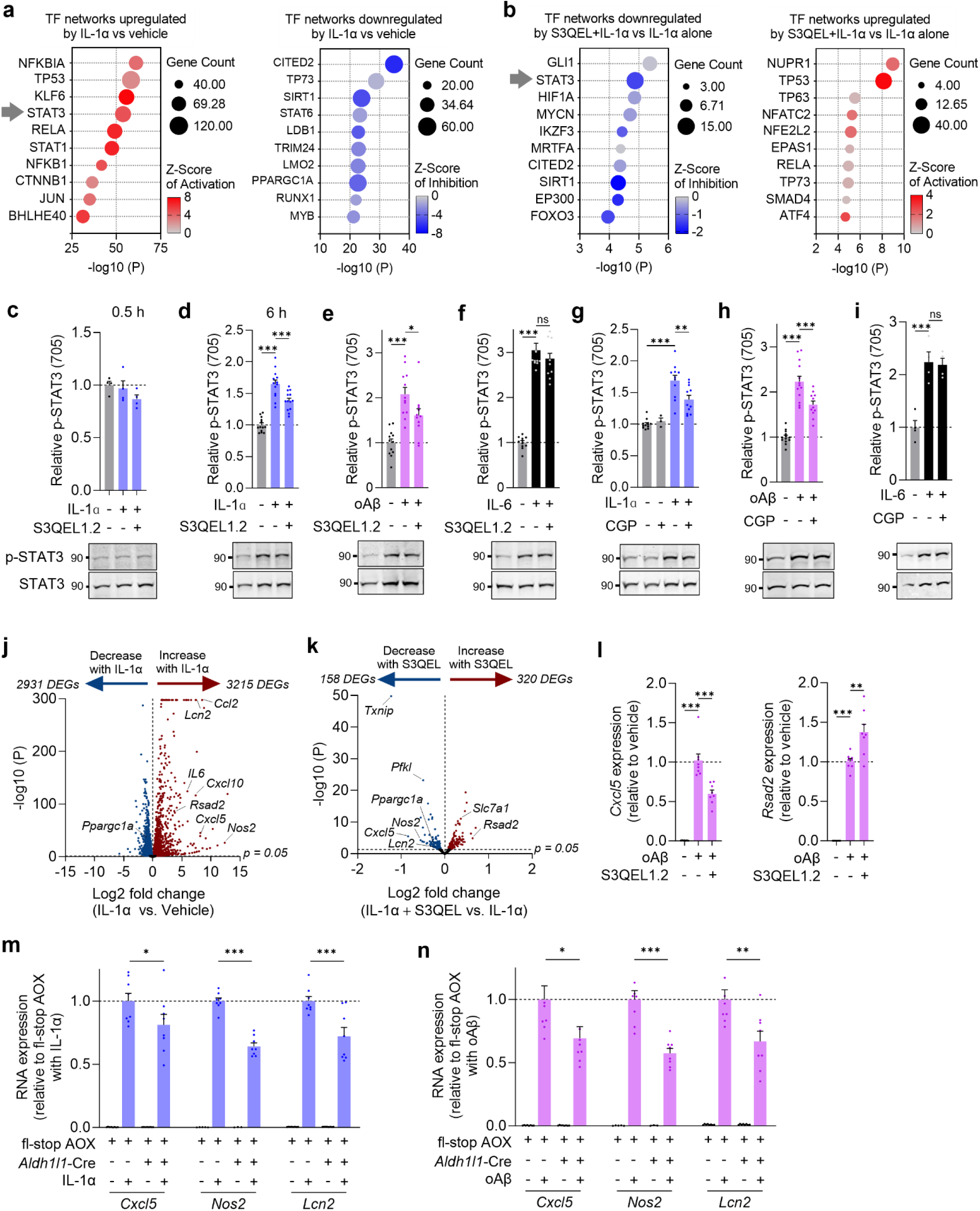
CIII-ROS promote context-specific STAT3 signaling and gene expression changes. a,b, Transcription factor network analysis of gene expression changes after 6-h treatment with vehicle, IL- 1α, or IL-1α with S3QEL1.2 (3 μM). n=3 replicate wells, listed factors exceed the p<0.05 threshold, Fischer’s exact test. **c**–**f**, Representative Western blots and quantification of p-STAT3(Y705) relative to total STAT3 levels after 0.5-h (c) or 6-h (d, e, f) treatments with vehicle, IL-1α, oAβ, IL-6 or S3QEL1.2. n=7–14 wells, one-way ANOVA with Tukey’s test. **g–i**, Representative Western blot and quantification of p-STAT3(Y705) levels after 1-h pre-treatment with CGP 37157 (CGP; 10 μM), then 6-h treatment with vehicle, IL-1α, oAβ, or CGP37157. n=3–13 wells, one-way ANOVA with Tukey’s test. **j,k**, Volcano plots of gene expression changes after 6-h treatments with vehicle, IL-1α, or IL-1α with S3QEL1.2. n=3 replicate wells. **l**, Relative RNA levels of *Cxcl5* and *Rsad2* after 6-h treatments with vehicle, oAβ, or oAβ with S3QEL1.2. n=7–8 wells, one-way ANOVA with Tukey’s test. **m,n**, Relative RNA levels of *Cxcl5*, *Nos2*, and *Lcn2* in control fl-stop AOX and *Aldh1l1*-Cre/fl-stop AOX astrocytes after 6-h treatment with vehicle, IL-1α, or oAβ. n=3–8 wells, two-way ANOVA with Bonferroni’s test. Data are shown as mean ± SEM. *p<0.05, **p<0.01, ***p<0.001.

To identify specific TFs downstream of CIII-ROS induction that may regulate transcriptional changes, we compared the TF networks differentially affected by IL-1α in the presence of S3QEL (Fig. 4b). STAT3 emerged as a top TF that was strongly involved in IL-1α-related gene expression changes and was downregulated by S3QEL (Fig. 4a,b). STAT3 is known to facilitate changes in astrocytic gene expression and cell function and to affect pathogenesis in CNS injury and disease.^89–96^ Of note, HIF-1α gene network activity was also reduced by S3QEL co-treatment, consistent with previous reports in other cell types that CIII-ROS regulates HIF-1α stabilization.^2,3,12^ Although S3QEL alone did not change basal mtHyPer7 oxidation (Fig. 1e), S3QEL did alter a small subset of basal transcripts (Extended Data Fig. 4a), suggesting that CIII-ROS might be produced at low levels under basal conditions, in agreement with previous reports.^59,97,98^ Basal genes and transcriptional networks affected by CIII-ROS are distinct from those affected by CIII-ROS in the context of IL-1α stimulation (Fig. 4b and Extended Data Fig. 4b).

To confirm the predictions from TF pathway analyses that CIII-ROS modulates STAT3 activity, we next measured the levels of phosphorylated STAT3 (p-STAT3, Y705).^99,100^ We found that IL-1α treatment increased astrocytic p-STAT3 levels in a similar delayed timeframe as mtROS induction (at 6 h but not 0.5 h; Fig. 4c,d), consistent with p-STAT3 being secondary to the early-phase activation of NF-κB. Suppression of CIII-ROS with S3QEL1.2 decreased IL-1α-induced p-STAT3 levels by 41% (Fig. 4d), similar to the 49– 56% decreases in IL-1α-induced mtROS (Fig. 1c,e). S3QEL1.2 also blocked IL-1α-induced nuclear translocation of p-STAT3 (Extended Data Fig. 4c). In contrast, inhibition of CI-ROS with S1QEL or inhibition of NOX-derived ROS with APX-115 did not affect p-STAT3 levels (Extended Data Fig. 4d,e), further highlighting the specific functional link between CIII-ROS and STAT3 activation. In line with its observed effects on CIII-ROS, oAβ also induced astrocytic p-STAT3, and this effect was suppressed by S3QEL1.2 (Fig. 4e). Although IL-1α increased the expression of other known STAT3 activators, including IL-6 and oncostatin M, the induction of these genes was not affected by S3QEL1.2 co-treatment or AOX expression, and astrocytic STAT3 responses to IL-6 and oncostatin M were not affected by S3QEL1.2 (Fig. 4f and Extended Data Fig. 4f–h). Together, these results suggest that the effects of CIII-ROS on STAT3 activity are likely independent of these autocrine mechanisms and that CIII-ROS modulates STAT3 activity in specific conditions. Indeed, we also found that inhibition of NCLX decreased p-STAT3 induction by IL-1α or oAβ (Fig. 4g,h) but did not affect p-STAT3 induction by IL-6 (Fig. 4i), further indicating that NCLX-dependent CIII-ROS is induced in specific pathogenic contexts to amplify STAT3 activity.

Transcriptional profiling also revealed that various genes known to be regulated by STAT3 (Fig. 4j,k; e.g., *Nos2*, *Lcn2*, *Rsad2*, and *Cxcl5*), including several secreted factors, are modulated by S3QEL. These genes are implicated in inflammation, astrocyte responses, and neurodegeneration,^58,89,101–111^ further linking CIII-ROS to disease pathogenesis. oAβ affected some of the same genes as affected by IL-1α, and S3QEL1.2 modulated these genes in a similar manner (Fig. 4i). AOX-mediated inhibition of CIII-ROS was also sufficient to decrease IL-1α- and oAβ-mediated changes in gene expression (Fig. 4m,n). Of note, the thioredoxin (Trx)-interacting protein TXNIP emerged as a transcript significantly suppressed by S3QEL at baseline and upon stimulation (Fig. 4k and Extended Data Fig. 4a). TXNIP binds and inhibits Trx, thereby preventing Trx-mediated reduction of intracellular thiols. Elevations in astrocytic TXNIP have been associated with inflammatory and dementia-related processes.^112,113^ Our data suggest that CIII-ROS is a regulator of astrocytic Trx function and intracellular redox environment through modulation of TXNIP expression. Thus, CIII-ROS induction promotes the expression of factors associated with pathological conditions and may contribute to a feedforward pathogenic process that leads to chronic disease.

Given our findings that CIII-ROS modulates astrocytic expression of factors that can affect neuronal health and function, we next tested whether CIII-ROS influences neuronal integrity and activities in a co-culture model of neurodegenerative pathology. In these experiments, to examine the effects of astrocytes on neuronal pathogenesis, we used a culture system with neuron-specific proteinopathy. Thus, we transduced primary mouse neurons with AAV vectors encoding a synapsin promoter-regulated mutant form of human tau (tauP301S, 1N/4R isoform) associated with early-onset Frontotemporal Dementia with Parkinsonism (FTD-P).^114,115^ Transduced tauP301S was localized throughout neuronal cell bodies and processes (Fig. 5a) and caused neuronal cell loss by six days after transduction (Fig. 5b–d). In these experiments and related *in vivo* testing described below, we used S3QEL2 rather than S3QEL1.2 due to its availability, better solubility in serum-free neuronal media, and superior oral bioavailability and brain penetrance in mice (Extended Data Fig. 5a–d).

**Figure 5.**
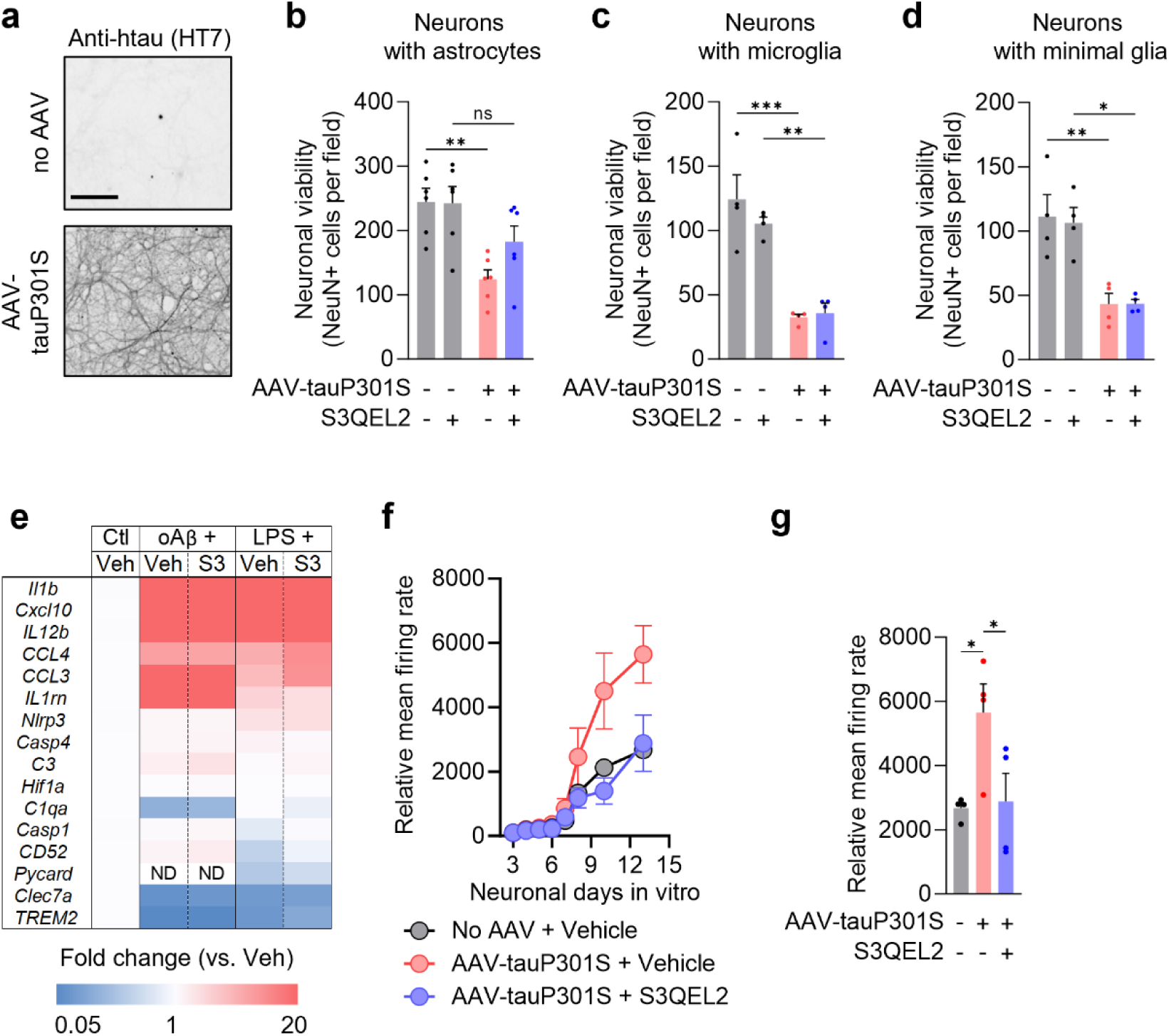
CIII-ROS promote tau-induced non-cell-autonomous neuronal degeneration. a, Human tau immunolabeling (clone HT7) in primary neurons untransduced or transduced with AAV-DJ- *hSyn*-tauP301S. Scale bar: 100 µm. **b**–**d**, Neuronal viability in astrocytic-neuronal co-cultures (**b**), microglial-neuronal co-cultures (**c**), or neuron-only cultures (**d**) untransduced or transduced with AAV-DJ-*hSyn*-tauP301S (1.2x10^9 particles per 15,000 neurons) and treated with vehicle or S3QEL2 (1 µM) for 4–6 days. n=4–6 wells per condition, 4–9 images per well, two-way ANOVA with Sidak’s multiple-comparison test. **e**, Inflammatory gene expression profiling in primary microglia stimulated with oAβ (3 µM) or lipopolysaccharide (LPS) (100 ng/mL) and treated with vehicle (Veh) or S3QEL2 (S3) (20 µM). **f,g**, Mean neuronal firing rates (% baseline) at indicated days *in vitro* (f) or on day 13 *in vitro* (g) in astrocytic-neuronal co-cultures untransduced or transduced with AAV-DJ-*hSyn1*-tauP301S (0.8x10^9 particles per 50,000 neurons) and treated with vehicle or S3QEL2 (1 µM). n=4 wells per condition, one-way ANOVA with Dunnett’s post-test. Data are shown as mean ± SEM. *p<0.05, **p<0.01, ***p<0.001.

We found that tauP301S-expressing neurons co-cultured with astrocytes had improved viability after S3QEL2 treatment as compared to vehicle-treated control co-cultures (Fig. 5b). To test whether S3QEL2 effects on neuronal viability can be mediated by microglia or cell-autonomous mechanisms in neurons, primary neurons were co-cultured with microglia or in the absence of added glia (approx. 95% neuronal purity), were transduced with AAV-*hSyn1*-tauP301S, and treated with S3QEL2 or vehicle for the duration of tauP301S expression. S3QEL2 did not affect neuronal viability in these cultures (Fig. 5c,d), suggesting that tauopathy-induced neurotoxicity was independent of microglial and neuronal CIII-ROS and that the effects of S3QEL2 were mediated non-cell-autonomously by astrocytes. In further support, S3QEL2 had minimal effects on inflammation-related gene expression changes in isolated primary microglia treated with lipopolysaccharide (LPS) or oAβ (Fig. 5e). Given that tauopathy can promote aberrant neuronal activity and hypersynchrony,^116^ we also tested neuronal firing patterns in primary neurons transduced with low amounts of AAV-*hSyn1*-tauP301S and co-cultured with astrocytes. TauP301S-expressing cultures had increased neuronal activity as compared to control cultures without tauP301S, and S3QEL2 normalized the activity of tauP301S-expressing neurons to control levels (Fig. 5f,g). Together, these results suggest that astrocytic CIII-ROS promote tauopathy-related neuronal dysfunction.

### Suppression of CIII-ROS alleviates dementia-linked pathology and extends lifespan in mice

We next explored whether CIII-ROS contributes to dementia-related neuropathology *in vivo* using mouse models. Pharmacokinetic measurements in mice indicated that S3QEL2 readily penetrates the gut and blood-brain barriers following oral administration (Extended Data Fig. 5c; brain-to-plasma ratio was approximately 2). Voluntary oral consumption of S3QEL2 formulated in almond butter enabled low-stress drug delivery and prolonged brain exposure (Extended Data Fig. 5d). S3QEL2 was well-tolerated when formulated in almond butter or rodent chow, which facilitated continuous and chronic drug delivery (Extended Data Fig. 5e,f). Chronic administration of high doses of S3QEL2 in chow for over 12 months had no detectable adverse health effects and did not alter body weight, metabolism, or general behavior in mice (Extended Data Fig. 5g–i).

To test the therapeutic potential of CIII-ROS suppression, we used the well-characterized transgenic mouse model of FTD-P (PS19 line),^117^ which expresses prion promoter-regulated mutant human tauP301S. TauP301S mice have robust tau hyperphosphorylation, weight loss (Extended Data Fig. 5j), progressive neuroinflammation (Extended Data Fig. 5k), and astrocytic and microglial alterations preceding hippocampal neuron loss and early mortality.^106,118–120^ To test if CIII-ROS suppression in adulthood alleviates ongoing tauopathy and related neuroimmune cascades, male tauP301S mice and nontransgenic (NTG) littermate controls received once daily oral administration of S3QEL2 formulated in almond butter starting after the onset of tau hyperphosphorylation, glial reactivity, and neuroinflammation (8.5–10 months of age). S3QEL2 administration continued for six weeks, which did not affect body weight (Extended Data Fig. 5e). Targeted transcriptional profiling in hippocampal tissue revealed that 10-month-old tauP301S mice had robust age-dependent changes in diverse gene transcripts linked to neuroinflammation and glial responses as compared to NTG littermates, and oral treatment with S3QEL2 for six weeks markedly reduced these changes (Fig. 6a). Upregulation of transcripts implicated in astrocytic responses in disease, including *Gfap* and complement component 3 (*C3*), were blunted by S3QEL2 (Fig. 6a). Astrocytic C3 promotes microglial reactivity and neuropathology in tauP301S mice.^106^ Indeed, S3QEL2 also suppressed genes involved in microglial reactivity and inflammasome-regulated cytokine production (Fig. 6a).

**Figure 6.**
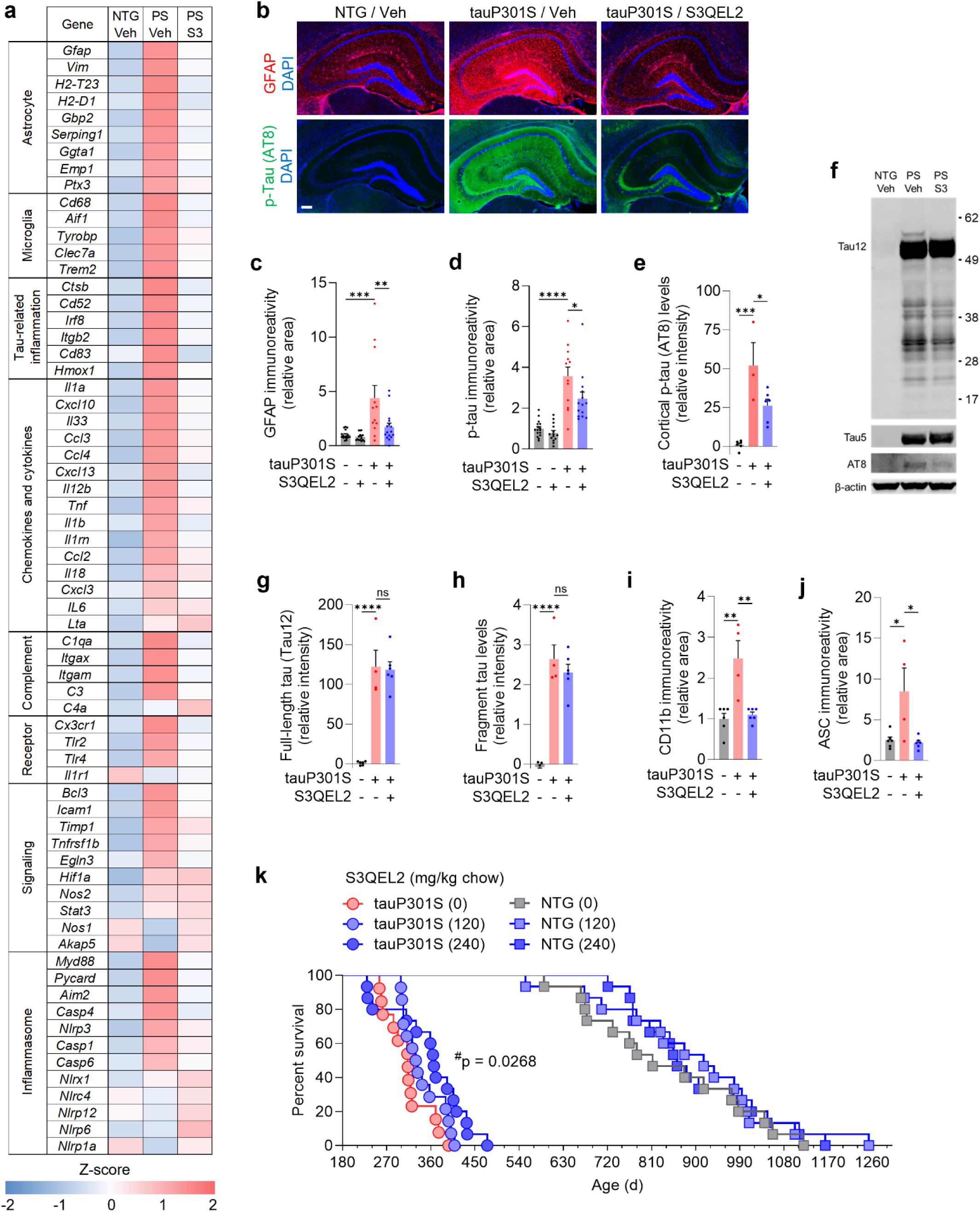
Chronic suppression of CIII-ROS reduces pathogenic cascades and early mortality in tauopathy. a, RT-qPCR quantification of neuroinflammatory and glial gene expression in hippocampal tissue of 10-month-old male nontransgenic (NTG) and hTauP301S (PS) mice after six weeks of oral administration with 5 mg/kg/day S3QEL2 (S3) or vehicle (Veh). Color indicates row Z-scores. n=3–5 mice per group. **b**–**d**, Representative images (b) and quantification (c, d) of GFAP and p-tau (AT8) immunolabeling in the hippocampus of NTG and tauP301S mice after six weeks of oral administration with 5 mg/kg/day S3QEL2 or vehicle. n=12–16 mice per group, ANOVA with Sidak’s test. Scale bar: 500 µm. **e**– **h**, Representative immunoblots (F) and quantification of p-tau (clone AT8) (e) or full-length and fragmented tau (clone Tau12) in NTG and tauP301S mice (g, h). n=4–6 mice per group, ANOVA with Sidak’s test. **i,j**, Quantification of CD11b and ASC (*Pycard*) immunolabeling in the hippocampus of NTG and tauP301S mice after six weeks of oral administration with 5 mg/kg/day S3QEL2 or vehicle. n=4–6 mice per group, ANOVA with Sidak’s test. **k**, Lifespan of NTG and tauP301S littermate mice treated with formulated chow containing S3QEL2 starting at four months of age. n=14–15 mice per genotype and dose. Pairwise Mantel-Cox test, ^#^p=0.0268 for tauP301S (240) vs tauP301S (0). Data are shown as mean ± SEM. *p<0.05, **p<0.01, ***p<0.001, ****p<0.0001.

We further examined the effects of S3QEL2 on neuropathological hallmarks at the protein level. Consistent with the transcriptional patterns, S3QEL2 treatment reduced hippocampal levels of glial fibrillary acidic protein (GFAP) and phosphorylated tau (AT8-reactive S202/T205) (Fig. 6b–d). TauP301S mice treated with S3QEL also had decreased cortical levels of phosphorylated tau (Fig. 6e,f). In contrast, S3QEL2 did not affect the total levels of human tau or tau fragments (Fig. 6f–h). S3QEL2 treatment also mitigated the increases in microglial CD11b protein and the inflammasome adaptor protein ASC (*Pycard*) (Fig. 6i,j). Together with our findings in isolated cells and co-cultures, these data suggest that targeting CIII- ROS may alleviate tauopathy-related pathogenesis.

We also tested two alternative dosing regimens by administering S3QEL2 to tauP301S mice either 1) continuously and chronically in chow instead of almond butter from 4 months of age until 10 months of age (“chronic dosing”, CD cohort), or 2) continuously and chronically in chow from 4 months of age through the natural lifespan (“chronic survival”, CS cohort). In contrast to the beneficial effects of S3QEL2 administered for six weeks via once daily feedings at 8–10 months of age (Fig. 6a–j), S3QEL2 administered continuously in chow did not affect hippocampal pathology in tauP301S mice in the CD cohort (Extended Data Fig. 6a–c), likely because S3QEL2 brain levels were 5-fold lower with *ad libitum* chow administration as compared to the levels reached with once daily dosing (Extended Data Fig. 5d vs. 5f). Nonetheless, tauP301S littermates that were randomly assigned to the CS cohort and received S3QEL2 in chow for their lifespan had an extended median age at death (366 vs. 313 d; 17% increase) and longer overall lifespan (475 vs. 396 d; 19.9% increase; p = 0.0268) as compared to tauP301S mice that received control chow with otherwise identical formulation (Fig. 6k). At advanced stages of pathology, tauP301S mice have progressive hindlimb paralysis and peripheral wasting.^117^ It is possible that S3QEL affected these or other disease processes that lead to early mortality. Thus, different drug delivery methods affect brain bioavailability and therapeutic effects of S3QEL *in vivo*. Collectively, our findings suggest that targeting CIII-ROS may represent an efficacious and well-tolerated therapeutic approach for dementia-linked pathogenesis and other conditions with aberrant glial responses and inflammation.

## Discussion

Mitochondria are increasingly recognized as sensors and integrators of diverse endogenous and external stimuli and as multifunctional regulators of cellular states and physiology, in part through mtROS induction and modulation of nuclear gene transcription.^121–125^ mtROS, particularly H_2_O_2_, are diffusible signals that communicate cell redox status primarily through reversible, post-translational oxidation of cysteines.^126,127^ Whether CIII-ROS-mediated cysteine modifications can serve as extra-mitochondrial signals that affect cell function has remained unclear.^122,128^ CIII-ROS have been implicated in the inactivation of cytosolic prolyl hydroxylases to promote hypoxic signaling^2^ and in the activation of macrophage NF-κB essential modulator (NEMO) during bacterial infection.^129^ Recently, proximity-dependent redox proteomics identified extra- mitochondrial cysteine modification following inhibition of CIII electron transport with antimycin A, but could not attribute these changes to any specific source.^130^ By combining CIII-ROS-selective pharmacology, complementary genetic manipulations, stochiometric redox proteomics, metabolomics, and transcriptomic profiling, we provide evidence for a signaling function of astrocytic CIII-ROS in disease (Fig. 7).

**Figure 7.**
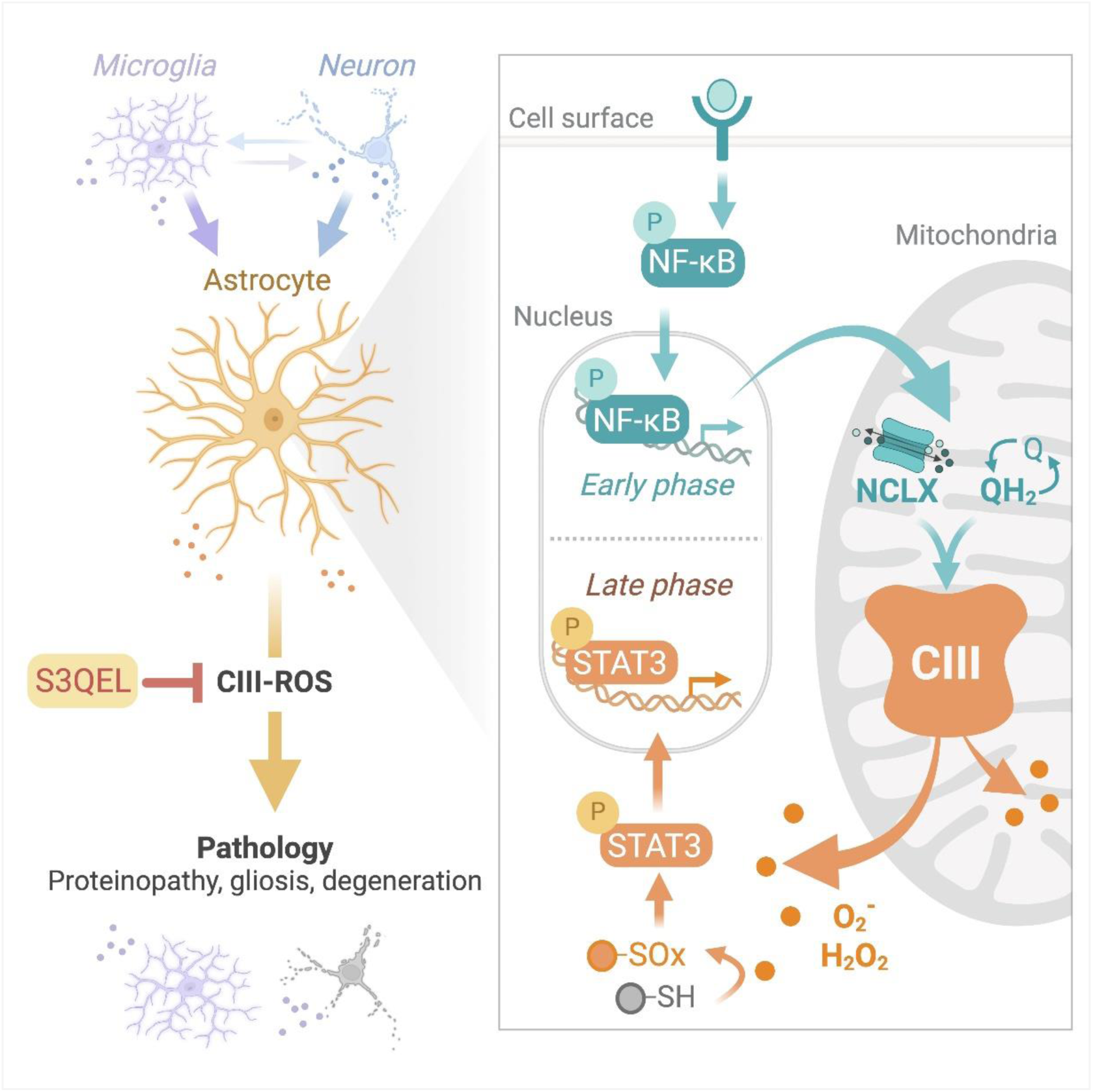
Summary of the molecular mechanisms and effects of astrocytic CIII-ROS. *Right:* Astrocytic mitochondria sense specific pathogenic cues through early-phase NF-κB activity that promotes NCLX- dependent CIII-ROS production. Increases in CIII-ROS lead to oxidation of select immunometabolic proteins and amplify late-phase transcriptional changes mediated by STAT3. *Left:* S3QELs selectively suppress CIII-ROS production and mitigate dementia-associated neuropathology.

In response to IL-1α, we did not detect generalized hyperoxidation of the cysteine proteome despite substantial increases in mtROS levels, with only 5% of cysteines changing their baseline oxidation by 10% or more. This high degree of selectivity in the redox proteome is consistent with observations in aging tissues.^86^ Targeted modification of specific cysteines may be caused by their close proximity to mitochondria and direct oxidation by CIII-ROS or by distinct redox relays acting over longer distances between subcellular compartments.^126,127,131–133^

S3QELs, which selectively suppress CIII-ROS, offer distinct advantages over broad antioxidants, genetic mutations, and potent respiratory or ribosomal inhibitors^11,130,134,135^ which disrupt cellular redox status and metabolism, among other cellular functions, and provide less insight into the dynamics and roles of individual sources of ROS. Our findings provide the first proteome-wide list of cysteines susceptible to redox modification by ROS emanating from a single, defined mitochondrial site. One target of IL-1α-induced CIII-ROS was electron transfer flavoprotein dehydrogenase (ETFD), an inner-mitochondrial protein that links fatty acid oxidation with the respiratory chain. ETFD was recently shown to interact with CIII^71^ and be oxidized by antimycin A-induced ROS in isolated mitochondria.^85^ S3QEL-mediated suppression of ETFD oxidation suggests that it is a direct target of CIII-ROS. Importantly, we identify the susceptible target as cys-386, which directly interacts with the FAD-cofactor within the FAD-binding domain.^136^ Mutations in this domain cause human metabolic disorders.^137–139^ Normal brain function relies on astrocytic fatty acid oxidation^140^ and disrupted lipid metabolism in astrocytes promotes neuroinflammation, in part through STAT3 activation.^141^ Therefore, CIII-ROS-mediated oxidation of the FAD-binding domain of ETFD might regulate astrocyte metabolism and coordinate responses to neuroinflammatory signals.

We also found that oxidation of mitochondrial malate dehydrogenase (MDHM) at cys-285 was increased approx. 10% by IL-1α and this increase was suppressed by S3QEL1.2. In support of a physiological role of this CIII-ROS-mediated reversible modulation of MDHM, our metabolomics data confirmed that the two tricarboxylic acid cycle intermediates immediately upstream of MDHM, malate and fumarate, were increased by IL-1α and blocked by S3QEL1.2, suggesting CIII-ROS is a dynamic modulator of metabolism. In agreement, our analysis identified numerous other metabolic and redox-related proteins susceptible to CIII-ROS, including FAS, HMGCL, CLYBL, MECR, ACSA, 6PGD and GSHR, further emphasizing the important roles of CIII-ROS in reinforcing changes to mitochondrial metabolic and redox states, in addition to regulating specific nuclear transcriptional programs. CIII-ROS-modulated cysteines also included several immune and disease-relevant proteins, including NFKB1, PP6R1, IL6RB, TRAF2 and GPC4 which, together with their sensitivity to S3QELs and recent advances in site-selective cysteine-reactive chemistry,^142–147^ may offer new targets for future therapeutic studies.

mtROS regulate immune signaling and inflammation^6,122,129,148,149^ and are closely linked to glial responses, cognitive deficits, and neuropathology.^8,11,26,49–52,140,150–152^ Our data point to STAT3 activation as a major functional effector of CIII-ROS induction in astrocytes. STAT3 is involved in astrocytic reactivity and diverse disease processes, such as chronic itch, CNS injury, tumor metastasis, drug withdrawal, and neurodegeneration.^56,88–91,93,105,106,153–159^ The induction of STAT3-regulated genes, such as *Lcn2*, *Nos2*, and *Cxcl5*, was blunted by S3QELs or AOX expression. Astrocytic LCN2 and CXCL5 are linked to inflammatory cascades and pathogenic processes.^110,160–164^ Similarly, NOS2 generates nitric oxide, a diffusible ROS that can also promote neurodegeneration.^165^ Of note, we also observed increased H_2_O_2_ extracellularly in stimulated astrocytes. The ability of S3QELs to lower releasable ROS may contribute to the non-cell autonomous protective effects of S3QELs in pathology. Indeed, astrocyte-derived signals have been reported to contribute to neurodegenerative pathology, including through a STAT3-dependent pathway involving astrocytic C3 release in tauP301S mice.^106^ A mixed STAT3/5 inhibitor, SH-4-54, was shown to reduce tau pathology and astrocytic and microglial changes,^106^ mirroring the effects of S3QEL2 in our study using the same mouse model. We additionally show that chronic S3QEL2 treatment extended lifespan in tauP301S mice. S3QELs were well-tolerated with chronic dosing and tended to delay mortality in healthy littermates as well. Taken together, our results suggest that S3QELs may alleviate neurodegenerative and inflammatory cascades that involve maladaptive, redox-dependent STAT3 activity.

The use of SELs has facilitated the identification of stimuli and contexts in which one site of mtROS production is selectively induced over others.^13,166–172^ However, evidence for site-specific mtROS production in neural cells and exact molecular mechanisms underlying site-selective induction have been lacking.^11^ Our study identifies several specific triggers and upstream regulators of CIII-ROS and the effects on pathology in co-cultures and mouse models of dementia-linked proteinopathy. We show that CIII-ROS production is dependent on NF-κB and NCLX activities and associated with increased ubiquinol (reported as AOX activation) and ΔΨm. Although the involvement of NF-κB in astrocytic and mitochondrial alterations, including mtROS, are known,^88,173–176^ here we establish the previously unknown requirement of NF-κB activation for induction of CIII-ROS. The activation of NF-κB was relatively rapid and independent of CIII- ROS, whereas activation of STAT3 was delayed and dependent on CIII-ROS. These findings suggest that CIII-ROS facilitate transcription factor cross-connectivity, linking early-phase NF-κB activation with late-phase STAT3 responses, thereby tuning gene expression and disease pathogenesis (Fig. 7).

NCLX, high ubiquinol, and hyperpolarization of ΔΨm have been linked to mtROS.^177–179^ In particular, NCLX was reported to influence mtROS in hypoxia, and increased ubiquinol and ΔΨm have been associated with mtROS from complex I.^1,180^ Here, we show that NCLX contextually regulates CIII-ROS even during stable O_2_ levels. Moreover, due to their enrichment in NCLX, astrocytic mitochondria appear functionally poised to produce high levels of CIII-ROS in pathological contexts. We also find that astrocytic CI-ROS is active under basal conditions in culture but does not increase further with stimulation. The lack of CI-ROS induction may be due to CI deactivation by some stimuli.^76,181^ Our findings suggest that there is a yet-to-be-identified molecular change that mediates the selective induction of CIII-ROS downstream of NCLX activation in normoxia.

In summary, our data show that CIII-ROS are important signal transducers, and that suppression of CIII-ROS can alleviate neuroimmune cascades and dementia-related pathology. We uncover that astrocytic CIII-ROS, but not other sources of ROS, have highly context-dependent responsivity to disease factors and affect specific signaling mechanisms and transcriptional changes linked to non-cell autonomous neurodegeneration and neural impairments. Our study identifies exact cysteine sites that are selectively oxidized in response to CIII-ROS and these modifications may represent functional mediators of redox signaling and new therapeutic targets for neurodegenerative conditions. Importantly, suppressors of CIII- ROS were well-tolerated and penetrant *in vivo* with no notable side-effects in animals treated for over one year with daily dosing. Thus, site-specific suppression of CIII-ROS may represent a promising therapeutic intervention in dementia and other neurological conditions that affect mitochondrial ROS production.

## Methods

### Reagents

Details for chemicals, antibodies, primers, and other reagents used are listed in Extended Tables 1 and 2. Amyloid-β oligomers were prepared from HFIP-dried or TFA-precipitated peptides (r-Peptide). TFA-peptides were solubilized into monomers with ice-cold HFIP for 2 h before evaporating overnight in a chemical safety cabinet. Oligomerization-state was evaluated by Western blotting, as previously described.^182^ S3QEL2 was sourced from Cayman Chemicals or custom synthesized by WuXi Apptec (Wuhan, China). Prior to use, the potency of new S3QEL2 stocks was evaluated in isolated skeletal muscle mitochondria, as originally described.^12^ The IC_50_ against CIII-ROS production for S3QEL2 preparations ranged from 1.4–1.7 µM, in agreement with the originally reported IC_50_ of 1.7 µM. S3QEL1.2 was custom synthesized by WuXi Apptec. During pre-HPLC (formic acid) purification, two stereoisomers were identified, separated as Peak 1 (*cis* arrangement of cyclohexane ring) and Peak 2 (*trans* arrangement), and distinguished by ^1^H-NMR (400 MHz, DMSO): S3QEL1.2 Peak 1(*cis*) = *δ* 8.16 (t, *J* = 5.6 Hz, 1H), 8.07 (dd, *J* = 7.8, 1.2 Hz, 1H), 7.78 - 7.84 (m, 1H), 7.63 (d, *J* = 8.0 Hz, 1H), 7.52 - 7.58 (m, 3H), 7.43 - 7.49 (m, 1H), 7.36 - 7.41 (m, 2H), 6.36 (dd, *J* = 3.0, 1.8 Hz, 1H), 6.15 - 6.18 (m, 1H), 4.55 (s, 2H), 4.20 (d, *J* = 5.8 Hz, 2H), 3.88 - 3.96 (m, 2H), 2.04 - 2.17 (m, 1H), 1.89 (s, 1H), 1.57 - 1.74 (m, 4H), 1.22 - 1.28 (m, 2H), 1.00 - 1.15 (m, 2H); S3QEL1.2 Peak 2 (*trans*) = *δ* 8.16 (t, *J* = 5.6 Hz, 1H), 8.07 (dd, *J* = 8.0, 1.2 Hz, 1H), 7.81 (td, *J* = 7.6, 1.6 Hz, 1H), 7.63 (d, *J* = 7.8 Hz, 1H), 7.52 - 7.58 (m, 3H), 7.43 - 7.49 (m, 1H), 7.35 - 7.40 (m, 2H), 6.36 - 6.39 (m, 1H), 6.17 - 6.20 (m, 1H), 4.55 (s, 2H), 4.25 (d, *J* = 5.6 Hz, 2H), 4.01 (br d, *J* = 6.4 Hz, 2H), 2.23 - 2.31 (m, 1H), 1.95 - 2.05 (m, 1H), 1.77 - 1.88 (m, 2H), 1.33 - 1.52 (m, 6H). During potency assessment against CIII-ROS, the IC_50_ of S3QEL1.2 Peak1 (*cis*) was determined to be 10-fold lower than S3QEL1.2 Peak 2 (*trans*) (52 nM versus 525 nM), so S3QEL1.2 Peak1 (*cis*) was used in all subsequent experiments and is referred to as S3QEL1.2 in the text. The IC_50_ of S3QEL1.2 Peak1 (*cis*) closely matches the expected IC_50_ based on estimates for unseparated S3QEL1.2 (approximately 50–100 nM).^12^

### Molecular biology and adeno-associated virus (AAV) preparation

The mtHyPer7 H_2_O_2_ sensor targeted to the mitochondrial matrix of astrocytes was PCR amplified from pCS2+MLS-HyPer7 (donated by Dr. Vsevolod Belousov, Addgene plasmid 136470; RRID: Addgene_136470) with primers containing SalI and HindIII restriction sites 5’ and 3’ to the coding sequence, respectively (SalI primer = 5’- ATA GTC GAC GAA TTC GCC ACC ATG; HindIII primer = 5’- GAG AAG CTT GCT AGC TCA ATC GCA GAT GAA GCT AA). Digested amplicon was ligated in place of the DIO-TDP43 sequence of pAAV-*GfaABC1D*-DIO-TDP43(WT)-WPRE-hGHpA^58^ to generate pAAV-*GfaABC1D*-mtHyPer7- WPRE-hGHpA.

Plasmids encoding codon-optimized Cliona intestinalis AOX and catalytically inactive AOX were cloned based on sequence in MAC_C_AOX (donated by Markku Varjosalo; Addgene plasmid 111661; RRID: Addgene_111661). An internal BamHI site was silently destroyed in both vectors and four missense mutations were taken from GenBank Accession MW222237 to generate catalytically inactive AOX. BamHI- AOX-NheI and BamHI-AOXmut-NheI DNA sequences without stop codons were synthesized and cloned into pUC-Amp by Genewiz (South Plainfield, NJ). Co-expression of AOX and mtHyPer7 in astrocytes was achieved by first replacing the NEUROD1 sequence in pAAV-*GfaABC1D*-NEUROD1-T2A-mCherry (donated by Chun-Li Zhang, Addgene plasmid 178582; RRID: Addgene_178582) with AOX or AOXmut using standard BamHI and NheI digestion and ligation to generate pAAV-*hGfaABC1D*-AOX-T2A-mCherry-WPRE-hGHpA and pAAV-*hGfaABC1D*-AOXmut-T2A-mCherry-WPRE-hGHpA. mtHyPer7 was cloned in place of the mCherry reporter in these vectors by removing the AgeI-mCherry-HindIII and replacing with PCR-amplified and digested Eco88I- mtHyPer7-HindIII using Eco88I-containing forward primer = 5’- CTA CCC GGG ATG TCC GTC CTG ACG CCG and HindIII-containing reverse primer = 5’- ACT AAG CTT TCA ATC GCA GAT GAA GCT.

Mutant human tauP301S was cloned for AAV-mediated, neuron-specific expression in two stages. First, the wild-type human tau sequence in a pcDNA3.1+ tau-WT-(1N4R) vector (provided by Dr. Wenjie Luo, Weill Cornell Medicine) was mutagenized with a Q5 Site-Directed Mutagenesis kit (New England Biolabs) to the P301S mutant using forward primer 5’-ATC AAA CAC GTC <u>T</u>CG GGA GGC GGC and reverse primer 5’-ATT ATC CTT TGA GCC ACA CTT GGA CTG GAC. Next, the validated tauP301S sequence was PCR amplified with primers containing SalI and EcoRI restriction sites 5’ and 3’ to the coding sequence, respectively (SalI primer = 5’- TAG ATA GTC GAC TCC GGA ATG GCT GAG CCC CGC C; EcoRI primer = 5’-GGC CGA GAA TTC CAA TTG TCA CAA ACC CTG CTT GGC CAG G). Digested amplicon was ligated in place of the DIO-rM3D(Gs)-mCherry sequence of Addgene plasmid 50458 (donated by Dr. Bryan Roth; RRID: Addgene_50458) to generate pAAV-*hSyn1*-tauP301S-(1N4R)-WPRE-hGHpA.

The AAV vector for astrocyte-specific Cre-dependent expression of 3x HA-tagged Rpl22^183^ was generated by PCR amplifying the Rpl22-3xHA sequence from pZac2.1-*GfaABC1D*-Rpl22-HA (donated by Dr. Baljit Khakh; RRID: Addgene_111811) using primers containing NheI and AscI restriction sites 5’ and 3’ to the coding sequence, respectively (NheI primer = 5’- CTA TAG GCT AGC GCC GCC ATG GCG CCT GTG AA; AscI primer = 5’- TCT AGA GGC GCG CCC TAC TGA GCA GCG TAA TC). Digested amplicon was ligated in place of the TDP43 coding region of pAAV-*GfaABC1D*-DIO-TDP43^58^ to generate pAAV- *GfaABC1D*-DIO-Rpl22-HA-WPRE-hGHpA.

NEB 5-alpha cells (New England Biolabs) were transformed with pAAV constructs and the integrity of inverted terminal repeats and expression-related elements in selected clones were confirmed by sequencing (Azenta/Genewiz) and restriction digests. AAV2/DJ*-hSyn1*-tauP301S-(1N4R) particles were produced by the Stanford Neuroscience Gene Vector and Virus Core Facility (RRID:SCR_023250). AAV2/PHP.eB-*hGfaABC1D*-DIO-Rpl22-HA-WPRE-hGHpA particles were produced by the UPENN Vector Core (RRID: SCR_022432).

### Mice

All animal experiments were conducted in accordance with guidelines set by the Institutional Animal Care and Use Committee of Weill Cornell Medicine. Mice were housed up to 5 mice per cage and maintained on a 12-h light/dark cycle with ad libitum access to food and water. Experiments were conducted during the light cycle and included littermate controls. Nontransgenic C57BL/6J mice (Jackson Laboratory stock #000664), *Aldh1l1*-Cre mice (B6;FVB-Tg(*Aldh1l1*-cre)JD1884Htz/J, Jackson Laboratory stock #023748), and floxed-stop AOX mice (fl-stop AOX)^69,184^ were used for primary cultures. *Aldh1l1*-Cre mice were backcrossed onto the C57Bl/6J background and express Cre recombinase downstream of the astrocytic *Aldh1l1* promoter. Fl-stop AOX mice were generously provided by Dr. Alexander Galkin (Weill Cornell Medicine). Transgenic mice expressing mutant human tau-P301S (B6;C3-Tg(*Prnp*- MAPT*P301S)PS19Vle/J, Jackson Laboratory stock #008169) and nontransgenic littermates were maintained on a B6/C3 hybrid background. Only male tauP301S mice were used in experiments because we and others have observed less pronounced and more variable pathology and early mortality in female littermates.^185^

### In vivo experiments

#### Pharmacokinetics

Levels of S3QEL2 and S3QEL1.1 in plasma from peripheral blood and saline-perfused brain tissue were quantified by LC-MS/MS using calibration standards. Oral gavage studies and associated MS-analysis were performed by XenoBiotic Laboratories/WuXi AppTec with timepoints between 0.25–24 h after administration of 5 mg S3QEL2/kg body weight using 5% DMSO, 5% Cremophor EL, 90% saline as vehicle. All other studies were conducted at Weill Cornell Medicine with perfused and flash-frozen brain tissue shipped on dry ice for LC-MS/MS analysis by Charles River Labs. For peripheral injections, S3QEL2 was suspended in DMSO and diluted to 5% DMSO, 5% Cremophor EL and 90% saline just prior to injection. Perfused tissue was collected 0–3 h after intraperitoneal injection. For oral administration in almond butter, S3QEL2 or S3QEL1.1 or DMSO vehicle were thoroughly mixed with a sterile spatula in almond butter (Justin’s Classic) pre-weighed into 30 mL plastic medicine cups. Dosing mixtures were then distributed to 15 mL (0.5 oz) plastic feeding cups according to the total mouse weight within each cage (divided to 1 or 2 cups for 1–3 or 4–5 mice per cage) to deliver an average of 5 mg S3QEL2 or 3 mg S3QEL1.1 per kg body weight. In time-monitored trials, all mice readily consumed 0.75 g almond butter per 30 g body weight within 2–3 h. For timed dosing trials with almond butter, tissue was harvested between 0–24 h. For chronic daily dosing with almond butter for up to 3 months, mice were weighed at least once weekly, and dosing adjusted accordingly. Daily dosing of 0.75 g almond butter supplement for this duration did not influence body weight. For chronic continual dosing with S3QEL2, chow was formulated with 0–480 mg S3QEL2 per kg Purina 5053 rodent chow by Research Diets (New Jersey). Irradiated chow was stored in Ziploc freezer bags at -80 °C for up to 6 months, and individual bags moved to 4°C storage for up to 1 month during feeding. The weights of mice, chow provided, and chow remaining in food hopper were measured every 1–7 days depending on the trial, stage, and duration.

#### Therapeutic dosing trials

TauP301S mice were dosed daily with 5 mg/kg S3QEL2 in almond butter from 8.5–10 months of age. One tauP301S mouse did not consume S3QEL-containing almond butter and was excluded from final analyses. Distinct cohorts of tauP301S mice were dosed with formulated chow (0, 120, or 240 mg S3QEL2 per kg chow) starting at 4 months of age. For pathological, biochemical, and molecular evaluations, tauP301S mice and nontransgenic littermates were euthanized at 9–10 months of age. For lifespan evaluations, cohorts were dosed continuously with chow until natural mortality or pre-established criteria for humane endpoints (e.g., loss of more than 20% peak body weight). Nontransgenic littermates of tauP301S mice were continuously dosed with 0–240 mg S3QEL2 per kg chow between 4–27 months of age and were assessed for VO2, VCO2, and distance traveled over 24 h using Promethion Metabolic Screening cages (Sable Systems). Cohorts were run in groups of 8 and analyses of metabolic data was performed by the WCM Metabolic Phenotyping Center.

### Cell culture experiments

All cultures were maintained at 37°C in a humidified 5% CO_2_-containing atmosphere. All primary cells were from cortices and hippocampi from mixed sex pups.

#### Primary astrocyte cultures

Pups at postnatal day 2–3 were dissected in cold PBS (Corning) to remove meninges and dissociated by manual trituration with a P1000 pipette in 1 mL fresh glial culture media consisting of high-glucose DMEM without glutamine or pyruvate (Corning) and supplemented with 20% heat-inactivated FBS (VWR), 1X GlutaMAX (ThermoFisher), and 1 mM sodium pyruvate (ThermoFisher). Cell suspensions were diluted to 3 mL with media, filtered through a 70-μm cell strainer (VWR), centrifuged at 500 g for 5 min at 22°C, resuspended with culture media, and plated into cell culture dishes (Corning Costar) or glass-bottom plates (Greiner Bio-One). All culture dishes and plates were pre-coated at ambient temperature for at least 30 min with sterile-filtered 100 µg/mL poly-D-lysine (Millipore-Sigma or ThermoFisher) dissolved in cell-culture grade water (Corning) and then rinsed thoroughly with water and dried prior to seeding cells. At DIV 4–5, the cells were washed in conditioned media to remove debris prior to replacement with fresh media. Cells were used for experiments after 7–14 days in culture. Primary astrocyte cultures were determined to contain greater than 96.5% astrocytes and less than 3.0% microglia, based on immunolabeling for glutamine synthetase, SOX9, Iba1 and 4′,6-diamidino-2-phenylindole (DAPI) (described under *Immunocytochemistry*). Unless specified otherwise, astrocytes were treated during a full media change to include one or more of the following reagents: 1–10 µM S3QEL1.2, 20–30 µM S3QEL2, 10 µM S1QEL2.2, 10–20 µM antimycin A, 3–10 µM myxothiazol, 3 ng/mL IL-1α, 300 nM oAβ, 30 ng/mL TNFα, 400 ng/mL C1q, 33 ng/mL IL-6, 10 ng/mL IFNγ, 5 µM APX-115, 1 µM TPCA-1, 10 µM CGP 37157, 1 µg/mL IL-1Ra, 3 µM IKK-16, 10 ng/mL oncostatin M. Stimuli and inhibitors were co-applied for entire treatment durations except for experiments that included TPCA-1, CGP 37157, IL-1Ra, and IKK-16 where astrocytes were pre-treated for 1 h prior to continued co-treatment with the specified cytokine or pathogenic factor.

#### Primary microglia cultures

Microglia were harvested from multiple 6-well plates of confluent primary astrocyte cultures at DIV 16–21 by pooling conditioned media with fresh media used to gently rinse loose, rounded microglia from atop the confluent astrocyte layer.^186^ Microglia-enriched suspensions were centrifuged at 1,000 g for 5 min and supernatants carefully aspirated. Pelleted cells were gently combined into a single tube and centrifuged again at 400 g for 3 min before carefully aspirating the supernatant and resuspending in glial culture media for microglia-only cultures or neuronal media if co-seeded atop primary neurons (described under *Primary microglia-neuron co-cultures*).

Three days after seeding in standard 24- or 96-well plates, microglia-only cultures were treated with 100 ng/mL lipopolysaccharides or 3 µM oAβ in prewarmed media together with DMSO vehicle or 10 µM S3QEL2 for 22–24 h prior to harvesting RNA (described under *qPCR*).

#### Primary neuron cultures

Forebrain neurons at postnatal day 0–1 were obtained as described previously^58^ with minor modifications. Briefly, papain-dissociated cells were filtered through a 40 µm cell strainer (Corning) to enrich for neurons and centrifuged at 500 g for 5 min to eliminate small debris. Neurons were suspended in complete neuronal medium consisting of Neurobasal (ThermoFisher) with B-27 supplement (ThermoFisher) and 1X GlutaMAX without antibiotics. Cells were seeded at 50–55K live cells per cm^2^ into plates coated with 100 µg/mL poly-D-lysine. Plating media was fully exchanged at DIV 1 using the same media except the substitution for B-27 supplement minus antioxidants (ThermoFisher). Half-media exchanges were performed every 3–4 days thereafter.

At 8–9 DIV, neurons in black-walled, clear-bottom 96-well plates (Corning) were transduced with 2.5 x 10^8^ particles of AAV2/DJ-*hSyn1*-tauP301S-(1N4R)-WPRE-hGHpA per well or treated with PBS vehicle and treated with 10 µM S3QEL2 (Cayman Chemical) or DMSO vehicle during a half-media exchange (with B-27 minus antioxidants). After 3 days, DMSO or 1x S3QEL2 were added with another half-media exchange. After 6 days post-transduction, neurons were fixed and labeled for high-content imaging, as described in *Immunocytochemistry*.

#### Primary astrocyte-neuron co-cultures

Postnatal day 0 neurons were isolated as described above but seeded onto near-confluent monolayers of DIV 4–7 astrocytes as described previously.^58^ Prior to seeding neurons, astrocytes were rinsed of debris and incubated with prewarmed neuronal media for 1 h to remove residual serum present in the glial media.

For endpoint imaging experiments, neurons were plated at 45,000 live cells per cm^2^ and cultured in black-walled 96-well plates as described in *“Primary neurons cultures*” including the switch to antioxidant-free medium 1 day after plating neurons and half-media exchanges every subsequent 3–4 days. Neurons were treated with 1 µM S3QEL2 or DMSO vehicle during a half-media exchange and transduced 15 min later with 1.2 x 10^9 particles of AAV2/DJ-*hSyn1*-tauP301S-(1N4R)-WPRE-hGHpA per well. Cultures were fixed and labeled for high-content imaging, as described in *Immunocytochemistry*.

For multielectrode array (MEA) experiments, neurons were seeded at 50,000 live cells per well in 48-well CytoView plates (Axion BioSystems). Neurons were seeded for MEA experiments in Neurobasal Plus culture system (ThermoFisher) with B-27 Plus supplement and 1X GlutaMAX as recommended by Axion BioSystems but switched to media containing B-27 minus antioxidants at the first half-media exchange at neuron DIV 5. Prior to this media exchange, neuronal activity was recorded for 30 min using the Maestro Pro MEA System at 37°C and 5% CO2 (Axion BioSystems). Recordings were analyzed using Neural Metric Software (Axion BioSystems). Following baseline recording, cells were transduced with 8 x 10^8 particles of AAV2/DJ-*hSyn1*-tauP301S-(1N4R)-WPRE-hGHpA per well or treated with PBS vehicle and treated with 1 µM S3QEL2 or DMSO vehicle during a half-media exchange (with B-27 minus antioxidants). Neuronal activity was recorded every 1–3 days for 10 days with fresh 1x S3QEL or DMSO added with half-media exchanges every 3 days.

#### Primary microglia-neuron co-cultures

Postnatal day 0 neurons were seeded at 55,000 per cm^2^, as described in *Primary neuron cultures*, in black-walled, clear-bottom 96-well plates and transduced at DIV 7– 8 by diluting 3.5 x 10^8 particles of AAV2/DJ-*hSyn1*-tauP301S-(1N4R)-WPRE-hGHpA per well in 25 µL reserved conditioned medium and adding to neurons without media exchange. The following day, microglia were harvested, as described in *Primary microglia cultures*, and the final suspension seeded atop neurons in 25 µL fresh neuronal media at 40–50K microglia per cm^2^ without media exchange. Microglia were allowed to settle onto neurons for 1–2 h in a cell culture incubator before DMSO vehicle or 3 µM S3QEL2 solutions were added in prewarmed, fresh neuronal media (B-27 minus antioxidants) during a half-media exchange. Cells were co-cultured for another 5 days prior to fixing and labeling for high-content imaging, as described in *Immunocytochemistry*.

### Immunocytochemistry

All immunolabeling steps were performed at ambient temperature unless specified otherwise. Briefly, cells were fixed with 4% paraformaldehyde and 4% sucrose in PBS (Corning) for 10 min, rinsed four times with PBS with 0.01% Triton X-100, and blocked and permeabilized in 5% normal goat serum and/or 5% normal donkey serum in 0.2–0.3% Triton X-100 in PBS for 1 h. Cells were incubated overnight at 4°C with primary antibodies diluted in 1% bovine serum albumin (BSA)(Sigma), 2% normal donkey serum, or 2% normal goat serum in 0.2–0.3% Triton X-100 in PBS: anti-human tau (clone HT7) (ThermoFisher, MN1000; 1:50), anti- MAP2 (Abcam, ab5392; 1:3200), anti-NeuN (Millipore-Sigma, ABN78; 1:1000), anti-Iba1 (Wako, 019-19741; 1:1000), anti-SOX9 (Millipore-Sigma, AB5535; 1:500), anti-glutamine synthetase (Abcam, Ab16802; 1:500), anti-STAT3 (Cell Signaling, 9139; 1:1500), anti-pSTAT3 (Y705) (Cell Signaling, 9145; 1:100), anti-GFP (Abcam, Ab6658; 1:750), anti-citrate synthase (Cell Signaling, 14309; 1:100). Cells were rinsed four times with PBS with 0.01% Triton X-100 and incubated for 1 h with 1:500 fluorescent-conjugated secondary antibodies (ThermoFisher) and 2.8 µM DAPI diluted in 1% BSA, 2% normal donkey serum, and/or 2% normal goat serum, and 0.2% Triton X-100 in PBS. Cells were rinsed twice with PBS with 0.01% Triton X- 100 and twice with PBS before imaging. For p-STAT3 and total STAT3 immunolabeling, astrocytes were fixed as described but permeabilized with pre-chilled 100% methanol at -20°C for 10 min according to manufacturer’s instructions. Methanol was aspirated and cells were rinsed with PBS and then blocked and immunolabeled as described above.

Co-localization of mtHyPer7 with the mitochondrial enzyme citrate synthase was estimated using linear regression of intensities across a 40 µm line ROI obtained using the Plot Profile function in Fiji software (v2.3).

### Immunohistochemistry

Mice were anesthetized with Avertin (2,2,2-tribromoethanol, 400–600 mg/kg body weight) and transcardially perfused for 2.5 min at 3 mL 0.9% saline per min before hemibrains were removed and stored in fixative (4% paraformaldehyde in PBS) overnight at 4°C on a rocking platform. Hemibrains were subsequently incubated in cryoprotectant (30% sucrose in PBS) for at least 48 h before cryosectioning.

Mouse brain tissue was sectioned (30 μm-thick sections) using a SM2010 R sliding microtome (Leica) equipped with a BFS-3MP freezing stage and cooling unit (Physitemp, Clifton, NJ). Free-floating sections were collected into cryopreservative (30% ethylene glycol, 30% glycerol in PBS) for long-term storage at -20°C.

Double or triple immunolabeling of free-floating mouse sections was performed with minor modifications depending on the antibodies used. All steps were performed at ambient temperature unless specified. Cryopreserved sections were thoroughly rinsed in PBS, permeabilized for 1 h in PBS containing 0.5% Triton X-100 (PBS-T), blocked with 10% donkey and/or goat serum in PBS-T for 1–2 h, incubated in primary antibodies in 3% serum in PBS-T for up to 48 h at 4°C: anti-phospho-tau (ser202, thr205) (clone AT8) (ThermoFisher, MN1020; 1:150), anti-GFAP (Millipore-Sigma, G9269; 1:1000), anti-ASC (Adipogen, AG-25B-0006; 1:400), anti-CD11b (Bio-Rad, MCA711G; 1:500). Sections were rinsed with PBS-T and protected from light in all subsequent steps. Tissue was incubated with 1:500 fluorescent-conjugated secondary antibodies (ThermoFisher) and 4.7 µM DAPI in 3% serum in PBS-T for 2 h rinsed with PBS-T, mounted, and dried on Superfrost glass slides (Electron Microscopy Sciences) before sealing beneath #1.5 coverglass (Corning) with Prolong Diamond Antifade Mounting Media (ThermoFisher). Slides were allowed to set overnight before acquiring images.

### Microscopy and image analyses

#### Fixed-cell or tissue imaging

Fixed and immunolabeled cultures were evaluated for phospho-tau or cell-type specific markers using procedures outlined previously.^58^ Briefly, plates were imaged using a 10X objective on an ImageXpress MICRO Confocal Automated High-Content Analysis System (Molecular Devices) at the Weill Cornell Medicine Automated Optical Microscopy Core Facility. For neuronal viability assays, 4 or 9 images were acquired around the center of each well and analyzed with the “Neurite Outgrowth” and “Multi Wavelength Cell Scoring” modules of MetaXpress software (Molecular Devices) to determine the number of NeuN-positive cells and total MAP2-positive neuronal process length in each image. NeuN immunolabeling was used as the nuclear-seed of MAP2-positive outgrowth.

For characterization of astrocyte cultures, fixed and labeled cells were manually counted as doubly positive for DAPI and astrocytic markers (glutamine synthase or SOX9) or microglial marker (Iba1) using a 20X objective on a BX-X710 microscope (Keyence).

To evaluate the intensity of protein expression in labeled mouse brain sections, slides were imaged on a BX-X710 microscope (Keyence) with a 10X or 20X objective (Nikon) using the tiling function. Images were stitched with BZ-X Analyzer Software (Keyence), and Fiji software (v2.3) was used for background subtraction and quantification of intensity and area of regions of interest, as previously described.^58^

#### Live-cell mitochondrial H_2_O_2_ imaging

Astrocytes were cultured on 96-well glass-bottom microplates (Greiner) or 10-chamber glass-bottom cell culture slides (Greiner) pre-coated with sterile-filtered 100 µg/mL poly-D-lysine, as described in *Primary astrocyte cultures*. Confluent astrocytes were transfected with 10 ng of pAAV-*GfaABC1D*-mtHyPer7 using Lipofectamine3000 (ThermoFisher) according to manufacturer instructions, rinsed with PBS after 5 h, and switched to imaging medium consisting of phenol red-free DMEM powder (Millipore-Sigma) supplemented with 20% heat-inactivated FBS, 1x GlutaMAX, 1 mM sodium pyruvate, 25 mM glucose, and 44 mM sodium bicarbonate, pH of 7.7 at ambient temperature and CO_2_ levels. The following day, astrocytes were stimulated for 0–8 h with cytokines, oligomeric Aβ, antimycin A, or vehicle controls without or with S3QEL or other pharmacological co-treatments. Cells expressing mtHyPer7 were imaged inside an environmental chamber at 37°C and 5% ambient CO_2_ with a 20X objective on a Zeiss LSM 880 confocal laser scanning microscope with spectral GaAsP detector or a Leica Stellaris 8 Confocal. Reduced and oxidized mtHyPer7 was detected by collecting 499–598 nm emission from sequential excitation with the 405 nm or 488 nm laser lines, respectively. Images from each channel were background-subtracted and fluorescence intensities within hand-drawn regions of interest encompassing entire individual astrocytes were determined for each wavelength using Fiji (v2.3). mtHyPer7 oxidized-to-reduced ratios were calculated for each cell, outliers were removed via ROUT analysis^187^ (GraphPad Prism), and ratios were normalized to vehicle within each experimental biological replicate.

### H_2_O_2_ efflux assay

Rates of H_2_O_2_ efflux were determined indirectly by HRP-coupled oxidation of Amplex UltraRed (Thermo Fisher), according to established protocols.^14,59^ Briefly, astrocytes were cultured on black-walled, clear-bottom 96-well plates (Corning Costar) pre-coated with poly-D-lysine, as described in *Primary astrocyte cultures*. Astrocytes were treated with cytokines, S3QELs, or other modulators in culture media for 6–24 h, rinsed once with Krebs-Ringer Buffer (135 mM NaCl, 5 mM KCl, 1 mM MgSO4, 0.4 mM K2HPO4, 20 mM HEPES pH 7.4) containing 25 mM glucose, 1x GlutaMAX, 1 mM sodium pyruvate, and 0.1% w/v bovine serum albumin (Sigma) prewarmed to 37°C, and then equilibrated in fresh prewarmed buffer for 30 min at 37°C and ambient atmosphere in a temperature-controlled BioTek Synergy H1 plate reader (Agilent). After equilibration, buffer was aspirated and replaced with H_2_O_2_ detection solution consisting of complete buffer with 25 μM AmplexRed, 25 U/mL superoxide dismutase, and 5 U/mL horseradish peroxidase. Fluorescence (530 nm excitation and 560 nm emission) was measured on a plate reader every 90 s for 45 min. Rates of H_2_O_2_ production were calibrated in parallel with each assay in cell-free control wells using H_2_O_2_ standard curves prepared in the detection solution. Following H_2_O_2_ measurements, media was aspirated, cells were washed once with DPBS, and then lysed for 10 min with 30 µL of RIPA buffer containing 10 mM Tris pH 7.5, 150 mM NaCl, 0.5% deoxycholate, 0.5% Triton X-100, 1X Complete Protease Inhibitor Cocktail (Millipore- Sigma) and 1% each of Phosphatase Inhibitor Cocktails 2 and 3 (Millipore-Sigma). 10 µL of lysate was transferred to a clear 96-well plate (Corning) and mixed with 350 µL of detergent-compatible Bradford reagent (Pierce) prior to measurement of absorbance at 595 nm on a BioTek Synergy H1 plate reader (Agilent). BSA standards were included in parallel blank wells.

### Cellular metabolism assays

Total ATP levels were measured using CellTiter-Glo 2.0 (Promega). Astrocytes were cultured in 96-well plates in standard medium, as described in *Primary astrocyte cultures*. The day prior to ATP measurements, growth medium was replaced with glucose-free DMEM (Millipore-Sigma) containing 20% heat-inactivated FBS, 1x GlutaMAX, 1 mM sodium pyruvate, and 10 mM galactose to enforce reliance on mitochondrial oxidative phosphorylation for cellular ATP production.^188^ After 24 h, medium was removed and astrocytes were treated for 10 or 30 min with indicated concentrations of S3QEL2, S3QEL1.2, S1QEL2.2, antimycin A, or myxothiazol in galactose-containing medium. At the indicated endpoints, 100 μL media was removed from each well and replaced with 100 μL pre-warmed CellTiter-Glo 2.0 reagent. Plates were mixed for 3 min on a benchtop shaker (VWR), then incubated without shaking for an additional 10 min at ambient temperature. Lysates were triturated to mix and 150 μL of each sample was transferred to a white 96-well plate (Costar) for luminescence measurement on a BioTek Synergy H1 plate reader (Agilent).

Oxygen consumption rates were measured on a Seahorse XFe96 analyzer (Agilent) according to manufacturer directions and as described previously.^35^ Briefly, primary astrocytes were plated on a 96-well Seahorse culture plate pre-coated with poly-D-lysine, as described in *Primary astrocyte cultures*. Prior to the assay, cells were incubated for 45 min with Seahorse XF Base Medium (Agilent), pH 7.4, supplemented with 20% FBS, 1 mM sodium pyruvate, 25 mM glucose and 2 mM glutamine (or 1 X GlutaMAX). Seahorse sensor cartridges (Agilent) were hydrated overnight in molecular biology grade water (Corning) and then calibrated in calibrant medium (Agilent) for at least 1 h prior to loading into the analyzer.

Compounds were added directly to wells just prior to the assay or, where indicated, cytokines, S3QEL or other modulators were loaded into the injection ports of pre-equilibrated sensor cartridges. To evaluate AOX function in transgenic astrocytes, myxothiazol and rotenone were injected sequentially following baseline respiration measurements. Oxygen consumption rates were determined using the Seahorse Wave software. Rates for individual wells were normalized to total protein using the detergent-compatible Bradford assay described in *H_2_O_2_ Efflux Assay*.

### Mitochondrial membrane potential

Tetramethylrhodamine methyl ester (TMRM) was used to assess the mitochondrial membrane potential. Astrocytes were cultured in 96-well glass-bottom microplates that were pre-coated with sterile-filtered 100 µg/mL poly-D-lysine, as described in *Primary astrocyte cultures*. Cells were switched to imaging medium consisting of phenol red-free DMEM powder (Millipore-Sigma) supplemented with 20% heat-inactivated FBS, 1x GlutaMAX, 1 mM sodium pyruvate, 25 mM glucose, and 44 mM sodium bicarbonate, pH of 7.7 at ambient temperature and CO_2_ levels. Prior to imaging, 5 nM TMRM was pre-equilibrated for 1 h prior to treatments except for time-courses lasting longer than 1 h. In these cases, TMRM was equilibrated during the final hour of treatment. Cells were imaged in an environmental chamber at 37°C and 5% ambient CO_2_ with a 20X objective on a Zeiss LSM 880 confocal laser scanning microscope with spectral GaAsP detector or a Leica Stellaris 8 Confocal. TMRM was excited at 548 nm and emission collected between 553–700 nm. Images were thresholded to a common value for each experiment to eliminate background (typically the average non-specific fluorescence within nuclear regions) and average fluorescence intensity of each image was determined using Fiji (v2.3).

### Polar Targeted Metabolomics

Astrocytes were cultured in 10cm plates that were pre-coated with sterile-filtered 100 µg/mL poly-D-lysine, as described in *Primary astrocyte cultures*. Astrocytes were then treated for 6 h with vehicle, IL-1α, IL-1α and TPCA-1, or IL-1α and S3QEL1.2 (n = 4 replicates per condition). Following treatment, polar metabolites were extracted on dry ice using pre-chilled 80% methanol (-80 °C). The extract was dried with a Speedvac, and redissolved in HPLC grade water before it was applied to the hydrophilic interaction chromatography LC-MS. Metabolites were measured on a Q Exactive Orbitrap mass spectrometer (Thermo Scientific), which was coupled to a Vanquish UPLC system (Thermo Scientific) via an Ion Max ion source with a HESI II probe (Thermo Scientific). A Sequant ZIC-pHILIC column (2.1 mm i.d. × 150 mm, particle size of 5 µm, Millipore Sigma) was used for separation of metabolites. A 2.1 × 20 mm guard column with the same packing material was used for protection of the analytical column. Flow rate was set at 150 μL/min. Buffers consisted of 100% acetonitrile for mobile phase A, and 0.1% NH_4_OH/20 mM CH_3_COONH_4_ in water for mobile phase B. The chromatographic gradient ran from 85% to 30% A in 20 min followed by a wash with 30% A and re-equilibration at 85% A. The Q Exactive was operated in full scan, polarity-switching mode with the following parameters: the spray voltage 3.0 kV, the heated capillary temperature 300 °C, the HESI probe temperature 350 °C, the sheath gas flow 40 units, the auxiliary gas flow 15 units. MS data acquisition was performed in the m/z range of 70–1,000, with 70,000 resolution (at 200 m/z). The AGC target was 1e6 and the maximum injection time was 250 ms. The MS data was processed using XCalibur (v4.1) (Thermo Scientific) to obtain the metabolite signal intensities. Identification required exact mass (within 5ppm) and standard retention times. Signal intensities were normalized to the total protein levels for each sample, uploaded to MetaboAnalyst (v6.0), then median normalized and log transformed.

### Western blotting

Cultured cells were rinsed with ice-cold PBS before lysis with ice-cold RIPA buffer containing 10 mM Tris pH 7.5, 150 mM NaCl, 5 mM EDTA, 0.5% deoxycholate, 0.5% Triton X-100, 1X Complete Protease Inhibitor Cocktail (Millipore-Sigma) and 1% each of Phosphatase Inhibitor Cocktails 2 and 3 (Millipore-Sigma). Collected lysates were sonicated on ice with a probe sonifier (Branson) for 5 s at 10% power, centrifuged at 10,000 g for 10–15 min at 4°C, and assayed for protein content using a detergent-compatible Bradford assay (ThermoFisher).

For brain tissue, cortices were homogenized in 150–200 µL RIPA lysis buffer containing HALT protease inhibitor (ThermoFisher) in a Fisherbrand Bead Mill 24 (Fisher Scientific) for 40 s at a speed setting of 5 in a pre-chilled adaptor tube rack. Samples were centrifuged at 1,000 g for 2 min at 4°C before sonication, lysate clarification, and protein content determination as described for cell culture lysates.

For all samples, 20 µg RIPA-soluble lysates were resolved on Bis-Tris SDS-PAGE gels (ThermoFisher) and transferred onto nitrocellulose membranes using a Mini Blot Module (ThermoFisher). Membranes were blocked with 5% BSA (VWR) in Tris-buffered saline (TBS) before probing overnight at 4°C with primary antibodies diluted in TBS containing 3% BSA and 0.2% Tween-20 (TBS-tw): anti-p-tau (ser202, thr205) (clone AT8, ThermoFisher, MN1020; 1:60), anti-GFAP (Millipore-Sigma, G9269; 1:1000), anti- STAT3 (Cell Signaling, 9139; 1:1000), anti-p-STAT3 (Y705) (Cell Signaling, 9145; 1:1000), anti-NF-κB (Cell Signaling, 6956; 1:1000), anti-p-NF-κB (S536) (Cell Signaling, 3033; 1:1000), anti-γ-tubulin (Millipore- Sigma, T5326; 1:1250), anti-β-actin (Millipore-Sigma, A2066; 1:2000), anti-total oxphos antibody cocktail (Abcam, Ab110413; 1:250), anti-citrate synthase (Cell Signaling, 14309; 1:1000). After overnight incubation in primary antibodies, all blots were rinsed with TBS-tw and probed with 1:15,000 IR Dye 680RD donkey anti-mouse and IR Dye 800CW donkey anti-rabbit fluorescent secondary antibodies (LI-COR) in TBS-tw with 3% BSA for 1 h. Blots were rinsed twice with TBS-tw, once with TBS, and dried for at least 1 h before scanning on the Odyssey CLx imaging system (LI-COR). Expression levels were quantified using LI-COR Image Studio software.

### Stoichiometric cys-redox proteomics

Quantification of the oxidation state of individual cysteine residues throughout the proteome was performed as described for cultured cells.^86^ Briefly, astrocytes treated with vehicle, IL-1α, or IL-1α and S3QEL1.2 (n = 3 replicates per condition) were harvested in ice-cold 20% trichloroacetic acid and lysed at 4°C with Lysing Matrix SS tubes (MP Biomedicals) on a Fisher Bead Mill 24 at speed 4 for 1.5 min. Lysates were divided into two 300-ug half-samples for reduction and 6C-CPT tagging (for “total” cys measurement) or iodoacetamide-blocking followed by reduction and CPT tagging (for “oxidized” cys measurement). Samples were further processed as originally described except the TMTpro 18-plex (ThermoFisher) was used for TMT-labeling. Pooled-sample TMT ratio checks, final LC-MS detection and analyses were performed by the Weill Cornell Medicine Proteomics and Metabolomics Core Facility. Raw files were processed using the MaxQuant computational proteomics platform (version 2.4.2.0) for protein and peptide identification. Fragment spectra were used to search the UniProt mouse protein database (downloaded on 09/21/2017).

TMT 18-plex on peptide N-term / lysine, oxidation of methionine, protein N-terminal acetylation and CPT on cysteine were used as variable modifications for database searches. Both peptide and protein identifications were filtered at 1% false discovery rate based on decoy search using a database with the protein sequences reversed. Cysteine-containing peptides were excluded if the probability of the identified cys- position was less than 95% (24 cysteine-peptides excluded). Total peptides in each channel were normalized based on the second ratio check of pooled TMT-labeled samples. For each cysteine-containing peptide, the TMT reporter ion intensity from the “Oxidized” channel was divided by the TMT reporter ion intensity from the “Total” channel to retrieve the % oxidation of each cysteine residue. Cysteine peptides with 0 values in the TMT reporter ion intensity of the “Total” channel were removed from analysis (130 cysteine-peptides excluded), as well as cysteine peptides that showed greater than 100% total oxidation (147 cysteine-peptides excluded). 8,263 cysteine-peptides were included in the final analysis after application of these criteria.

### Cys-redox proteomics gene ontology enrichment and subcellular localization analysis

Gene ontology enrichment analysis was performed using National Institutes of Health (NIH) Database for Annotation, Visualization and Integrated Discovery (DAVID).^189,190^ The input consisted of cysteines that increased oxidation 10% or more following IL-1α stimulation compared to vehicle (160 cysteines) and the Mus musculus genome was used as the background. P values were adjusted using the Benjamini-Hochberg correction. GO terms in the category GOTERM_BP_1 were plotted. The GO annotations are as follows: GO:0008152 (metabolic process), GO:0071840 (cellular component organization or biogenesis), GO:0051179 (localization), GO:0007610 (behavior), GO:0032502 (developmental process).

To determine the subcellular localization of proteins containing oxidized cysteines, experimentally validated protein localization data were downloaded from the Human Protein Atlas. Subcellular localizations from the atlas were then mapped onto proteins that contained cysteines that increased oxidation 10% or more following IL-1α stimulation (160 cysteines). Proteins without assigned locations were excluded from the analysis (36 cysteines). Proteins with one or more subcellular localizations in the following categories were grouped under single locations: nucleus = annotation for nucleus, nucleoplasm, nucleoli, nuclear bodies, nuclear fibrillar center, nuclear speckles, or nuclear membrane; cytoskeleton = annotation for cytoskeleton, actin filaments, cytokinetic bridge, microtubules, intermediate filament, centrosome, focal adhesion sites, midbody, or cell junctions; vesicles = vesicles or cytoplasmic bodies. All other proteins contained localizations that fit into single categories of cytosol, plasma membrane, cytoskeleton, Golgi apparatus, mitochondria, peroxisome, or endoplasmic reticulum. The number of proteins assigned to these compartments was summed and plotted.

### Retro-orbital injection and immunoprecipitation of ribosome-bound RNA

To confirm *Slc8b1* enrichment in astrocytes *in vivo*, *Aldh1l1*-Cre mice were briefly anesthetized with isoflurane (3%, Covetrus) and AAV2/PHP.eB-*hGfaABC1D*-DIO-Rpl22-HA (1 x 10^11^ Vg) diluted in saline to 100 µL was injected into one retro-orbital sinus. Petrolatum ophthalmic ointment (Puralube Vet Ointment) was applied to the injected eye and vectors were allowed to express for three weeks before isolating total RNA and astrocyte-specific ribosome-bound RNA. Mice were sacrificed by cervical dislocation and dissected cortices were thoroughly homogenized using glass Dounce homogenizers (20 times with each pestle and 0.013–0.064 mm final clearance; Kimble Chase) in polysome buffer consisting of 50 mM Tris pH 7.5, 100 mM KCl, 12 mM MgCl_2_, 1% Nonidet P-40, 1 mM DTT, 200 U/mL RNasin (Promega), 1 mg/mL heparin, 100 g/mL cycloheximide, cOmplete protease inhibitor tablets (Sigma) at 10% wet weight/volume. Samples were transferred to a fresh RNase-free tube (Eppendorf), centrifuged at 10,000 g for 10 min at 4°C, and 20% of cleared supernatant was frozen at -80°C until subsequent extraction for total RNA input. The remaining 80% cleared supernatant was transferred to a clean RNase-free tube and incubated with mouse anti-HA antibody (1:250; clone 16B12, Biolegend) on a rotating mixer for 4 h at 4°C while 300 uL protein G magnetic beads (Dynabeads; Invitrogen) were washed on a magnetic stand (ThermoFisher) with 500 μL citrate-phosphate buffer consisting of 24 mM citrate and 52 mM phosphate at pH 5.0 followed by three 500 μL washes with immunoprecipitation buffer consisting of 50 mM Tris pH 7.5, 100 mM KCl, 12 mM MgCl_2_ and 1% Nonidet P-40. After anti-HA incubation, homogenates were added directly to the washed magnetic beads and allowed to incubate on a rotating mixer overnight at 4°C after which the beads were placed on a magnetic stand, the supernatant with unbound RNA was aspirated, and Rpl22-HA-coupled beads were washed three times with high-salt buffer containing 50 mM Tris, pH 7.5, 300 mM KCl, 12 mM MgCl2, 1% Nonidet P-40, 1 mM DTT, and 100 ug/mL cycloheximide. After the final wash, 200 μL RNA lysis buffer consisting of 1% β-mercaptothanol in RLT Plus (Qiagen) was added to the beads, vortexed to lyse Rpl22-HA-bound RNA and transferred to a clean RNase-free tube. The extraction was repeated with 150 μL lysis buffer and the combined 350 μL bound RNA fractions processed for RNA extraction using the RNeasy Plus Micro kit (Qiagen 74034). The total RNA input from the 20% reserved cleared homogenate was thawed on ice and extracted using the RNeasy Mini kit (Qiagen 74106) with on-column DNase treatment (Qiagen 79256) according to manufacturer instructions.

### RT-qPCR

Bulk RNA was extracted using the RNeasy Mini Kit with on-column DNase treatment. Cultured primary cells were rinsed once with ice-cold PBS and collected in freshly prepared extraction buffer consisting of 1% β- mercaptothanol in RLT (Qiagen). Saline-perfused, microdissected mouse brain tissue was frozen on dry ice and stored at -80°C until RNA extraction. Tissue was homogenized for 20 s in fresh extraction buffer using a Fisher Scientific Bead Mill 24 at a speed setting of 5 in a pre-chilled adaptor tube rack.

Microfluidic RT-qPCR was performed as previously described.^58^ Briefly, cDNA was synthesized with Protoscript First Strand Synthesis Kit (NEB), treated with RNase H (NEB), and pre-amplified for 14 cycles against a pool of primers (Extended Data Table 1) using PreAmp Grandmaster mix (TATAA Biocenter) before exonuclease I treatment (NEB). Pre-amplified cDNA was diluted at least five-fold with nuclease-free water and mixed with SsoFast EvaGreen with Low ROX (BioRad) and chip-specific DNA Sample Reagents before loading into primed Flex Six or 96.96 Dynamic Array chips (Standard Biotools). Individual primers were mixed with DNA assay reagent (Standard Biotools) and loaded into chip inlets. Chips were primed and loaded using an IFC Controller HX (Standard Biotools) before measuring and analyzing amplification and melting curves on a BioMark HD System (Standard Biotools). Cycle of quantification (Cq) values were thresholded equally for all inlets across each chip run and normalized to the average of reference genes (Actb and Gapdh for tissue samples; Actb and/or Tbp and/or Gusb for cultured cells) before determining ddCq and fold-change relative to experimental control groups.

For targeted analysis of specific genes, RT-qPCR was performed on cDNA using PowerUp Sybr Green Master Mix (ThermoFisher) and a CFX96 Touch Real-Time PCR Detection System (BioRad). All qPCR primer sequences are detailed in Extended Data Table 1.

### RNA sequencing

RNA was extracted and cDNA synthesized as described for qPCR. cDNA was submitted to the Weill Cornell Genomics Resources Core Facility for quality control analysis and bulk RNA sequencing (n = 3 biological replicates per condition). The libraries were sequenced with paired-end 50 bps on a NovaSeq6000 sequencer. The raw sequencing reads in BCL format were processed through bcl2fastq (v2.20) (Illumina) for FASTQ conversion and demultiplexing. The adaptors were trimmed with cutadapt (v1.18) (https://cutadapt.readthedocs.io/en/v1.18/), RNA reads were aligned and mapped to the GRCm39 mouse reference genome by STAR (v2.5.2) (https://github.com/alexdobin/STAR),^191^ and transcriptome reconstruction was performed by Cufflinks (v2.1.1) (http://cole-trapnell-lab.github.io/cufflinks/). The abundance of transcripts was measured with Cufflinks in Fragments Per Kilobase of exon model per Million mapped reads (FPKM).^192,193^ Raw read counts per gene were extracted using HTSeq-count (v0.11.2).^194^ Gene expression profiles were constructed for differential expression, cluster, and principle component analyses with the DESeq2 package (https://bioconductor.org/packages/release/bioc/html/DESeq2.html).^195^ For differential expression analysis, pairwise comparisons between two or more groups were made using parametric tests where read-counts follow a negative binomial distribution with a gene-specific dispersion parameter. Corrected p-values were calculated based on the Benjamini-Hochberg method to adjusted for multiple testing. Corrected p-values of 0 were corrected to the minimum calculated non-zero value.

### Gene network analyses

Qiagen Ingenuity Pathway Analysis (IPA) was used to identify gene networks which regulate and are regulated by CIII-ROS. Differentially expressed genes (DEGs) were imported into IPA with log2 fold change values and -log10 p-values to identify involved pathways. The Fisher’s exact test was used to calculate the p-value of the overlap of our dataset with those genes of a reference set known to be regulated by particular upstream factors, with a statistical significance level p < 0.05. Z-scores were also calculated, with positive and negative scores indicating activation and inhibition of an upstream regulator, respectively. To identify S3QEL-mediated changes relevant to IL-1α, IPA analysis was performed on the gene list identified from the IL-1α+S3QEL vs. IL-1α comparison but excluding genes differentially changed in the S3QEL vs. Vehicle comparison.

### Statistical analyses

Statistical specifications are reported in the figures and corresponding figure legends. Data are presented as mean ± S.E.M. All statistical tests were performed using GraphPad Prism 10, except Fisher’s exact test and other statistical tests included in the Qiagen Ingenuity Pathway Analysis. The criterion for data point exclusion was established during the design of the study and was set to values above or below two standard deviations from the group mean or, in the case of cell culture studies, including mtHyPer7 ratios, removed by ROUT analysis.^187^ Two-sided Student’s t test was used to determine statistical significance between two groups. Differences among multiple groups were assessed by one-way or two-way ANOVA or followed by Tukey’s or Dunnett’s multiple comparisons post-hoc tests, as specified in the legends. mtHyPer7 measurements were not normally distributed according to the Kolmogorov-Smirnov test and were analyzed using non-parametric tests. The Mann-Whitney U test and the Kruskal-Wallis test were used to assess differences between two or more groups, respectively. Null hypotheses were rejected at p < 0.05.

## Resource Availability

Further information and requests for resources and reagents should be directed to the lead contact, Adam L. Orr.

## Materials availability

Plasmids generated in this study are available from the lead contact but may require a completed materials transfer agreement.

## Data and code availability

- All raw data reported in this paper are available upon request from the lead contact.
- This paper does not report original code.
- Any additional information required to reanalyze the data reported in this work is available from the lead contact upon request.

## Acknowledgements

We thank S. Tymchuk and M.A. Fatafta for technical assistance; G. Coronas-Samano, A. Carney and M. Garvey for administrative assistance; W. Luo for providing the human tau(1N4R) cloning vector, primary microglia cultures and lipopolysaccharide; A. Galkin for the fl-stop AOX mice; and L. Cantley for access to the Seahorse instrument. We thank G. Manfredi, R. Ratan, and J. Anrather for advice on project execution. We thank the Weill Cornell Medicine core facilities, including the Proteomics and Metabolomics Core, the Genomics Resources Core, the Metabolic Phenotyping Center, and the Microscopy and Imaging Core for assistance and resources. All schematics were generated with BioRender.com. This work was funded by the National Institutes of Health: R01AG068091 (A.G.O.), F31AG084165 (D.B.), K99AG073461 (H.X.), DK123095 and AG071966 (E.T.C.), Alzheimer’s Association: AARG-17-533273 (A.G.O.) and 23AARF- 1029892 (T.S.Z.), BrightFocus Foundation: A2019363S (A.G.O.) and BFA2023008F (T.S.Z.), Leon Levy Foundation (A.G.O.), Alzheimer’s Drug Discovery Foundation/Association for Frontotemporal Dementia: GA-202007-2020603 (A.G.O.), Sanofi Innovation iAward (A.L.O.), the Claudia Adams Barr Program (E.T.C), the Lavine Family Fund (E.T.C), the Pew Charitable Trust (E.T.C), the Smith Family Foundation (E.T.C), the American Federation for Aging Research (E.T.C) and the Weill Cornell Medicine Daedalus Fund for Innovation (A.L.O.).

## Author Contributions

Conceptualization: D.B., A.G.O., A.L.O.; Methodology: D.B., T.S.Z., H.X., E.T.C., A.G.O., A.L.O.; Investigation: D.B., T.S.Z., C.B., F.P., S.M.M., A.G.O., A.L.O.; Formal Analysis: D.B., T.S.Z., C.B., A.G.O., A.L.O.; Visualization: D.B., A.G.O., A.L.O.; Funding acquisition: D.B., T.S.Z., A.L.O., A.G.O., E.T.C.; Project administration: A.G.O., A.L.O.; Supervision: A.G.O., A.L.O.; Writing – original draft: D.B., A.G.O., A.L.O.; Writing – review and editing: all authors.

---

**Extended Data Fig. 1.**
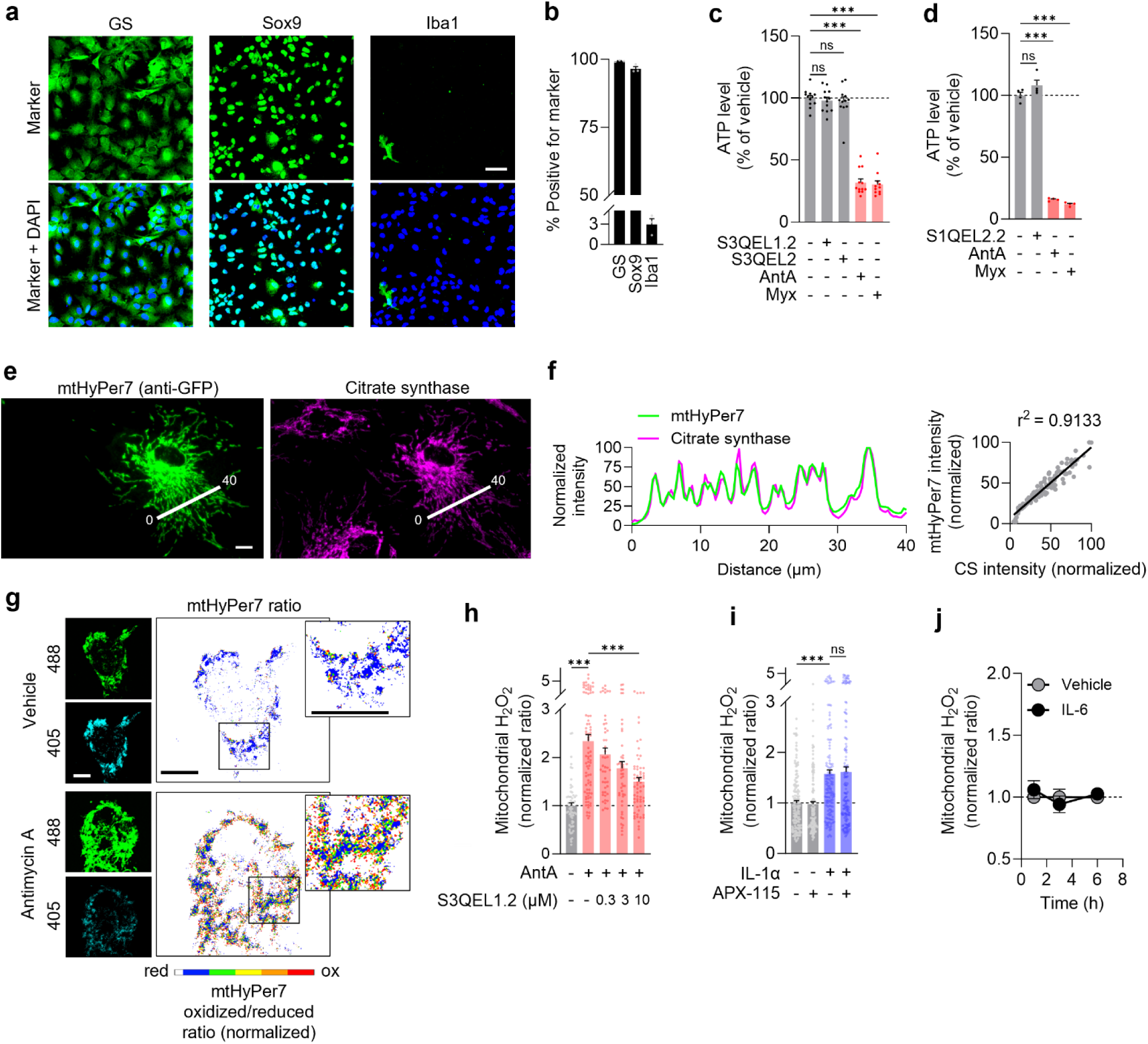
Further characterization of S3QELs, mtHyPer7, and ROS production. a,b, Representative immunostaining and quantification of glutamine synthetase (GS), SOX9, and Iba1 in primary astrocyte cultures. n = 3 wells. Scale bar: 50 µm. **c,d**, ATP levels after treatment with S3QEL1.2 (3 µM), S3QEL2 (20 µM), S1QEL2.2 (10 μM), antimycin A (AntA, 10 μM) or Myxothiazol (Myx, 3–10 μM). n=4–12 wells, ANOVA with Tukey’s test. **e,f**, Representative immunolabeling and correlational analysis of mtHyPer7 (anti-GFP) and mitochondrial citrate synthase. Scale bar: 10 µm. **g,h**, Representative images and quantification of mtHyPer7 after 1 h co-stimulation with vehicle, AntA (20 μM) or S3QEL1.2 (0.3–10 µM). Scale bars: 10 µm. n = 47–83 cells, Kruskal Wallace with Dunn’s test. **i**, Ratiometric mtHyPer7 measurement after 6 h co-stimulation with vehicle, APX-115 (5 μM), or IL-1α. n = 137–149 cells, Kruskal Wallace with Dunn’s test. **j**, Ratiometric mtHyPer7 measurement after 1–6 h treatment with IL-6. n = 73–177 cells, Mann-Whitney, unpaired two-tailed t-test. Data are shown as mean ± SEM. *p<0.05, **p<0.01, ***p<0.001.

**Extended Data Fig. 2.**
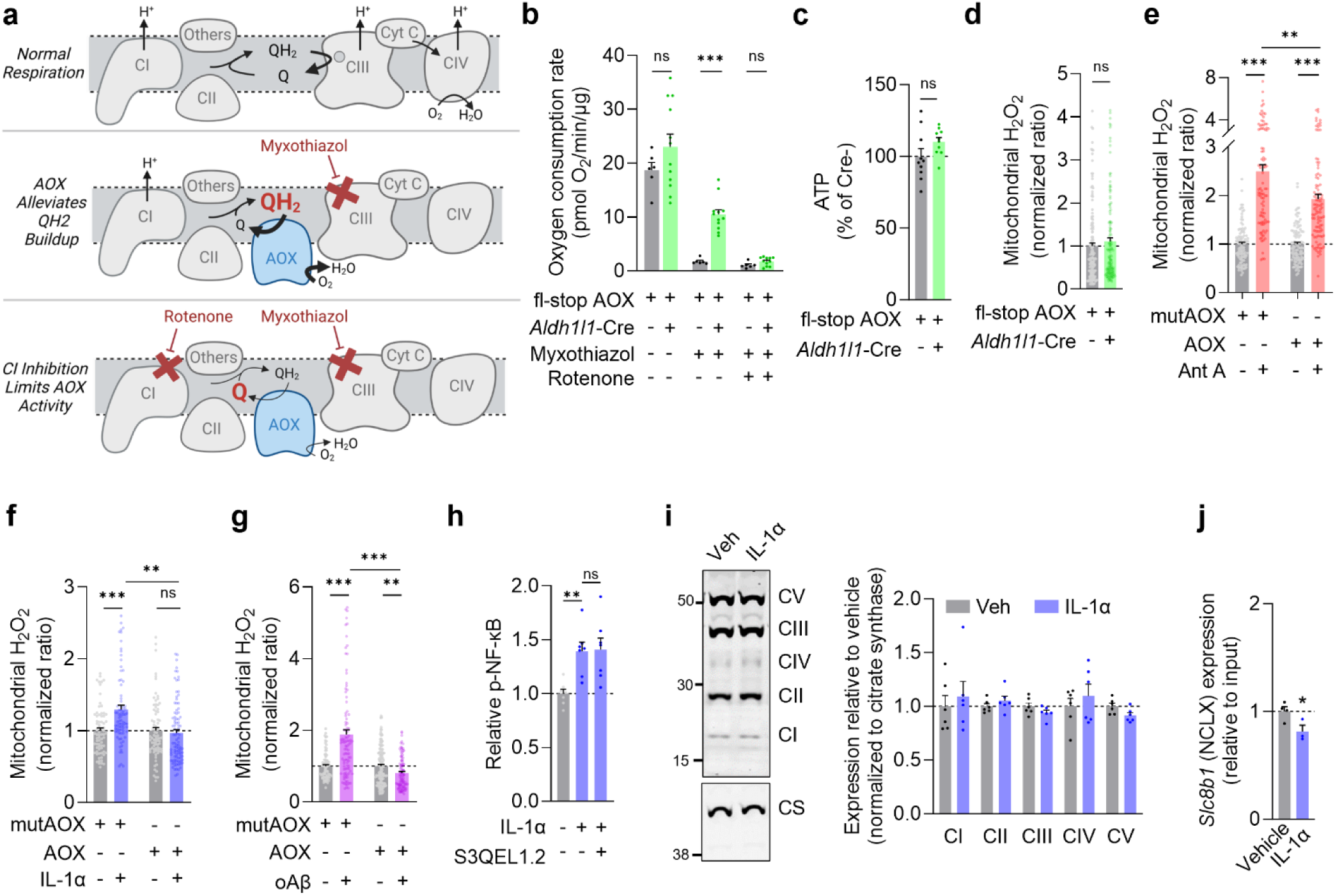
Further characterization of CIII-ROS regulation. a, Schematic depicting electron flow to ectopically expressed alternative oxidase (AOX) in the presence of inhibitors of CIII or CI. AOX is activated and reduces O_2_ to H_2_O when QH_2_ increases in response to complete inhibition of CIII by myxothiazol (top panel). In contrast, rotenone prevents QH_2_ formation at CI to limit AOX-dependent oxygen consumption (bottom panel). **b**, Oxygen consumption rates in astrocytes derived from *Aldh1l1*-Cre/fl-stop AOX or control fl-stop AOX mice at baseline and after sequential addition of myxothiazol (3 μM) and rotenone (3 μM). n = 6–12 wells, unpaired, two-tailed t-test. **c**, Mitochondria-enriched ATP levels in *Aldh1l1*- Cre/fl-stop AOX or control fl-stop AOX astrocytes. n = 10 wells, unpaired, two-tailed t-test. **d**, Quantification mtHyPer7 fluorescence in *Aldh1l1*-Cre/fl-stop AOX or control fl-stop AOX astrocytes. n = 135–143 cells, Mann-Whitney, unpaired, two-tailed t-test. **e**–**g**, Quantification of mtHyPer7 in astrocytes expressing wild- type AOX or catalytically inactive mutant AOX (mutAOX) and treated with vehicle, antimycin A (Ant A, 20 μM) (3 h), oAβ (3 h), or IL-1α (6 h). n = 40–101 cells, Mann-Whitney, unpaired, two-tailed t-test. **h**, Quantification of p-NF-κB after 30-min treatment with vehicle, IL-1α, or S3QEL1.2 (3 μM). Normalized to total NF-κB levels. n=7 wells, one-way ANOVA with Tukey’s test. **i**, Representative Western blots and quantification of electron transport chain complexes I–V (CI–CV) and the matrix enzyme citrate synthase (CS) after 6 h treatment with vehicle or IL-1α. n = 6 wells, unpaired, two-tailed t-test. **j**, *Slc8b1* (NCLX) mRNA expression after 6 h treatment with vehicle or IL-1α. n = 3–4 wells, unpaired, two-tailed t-test. Data are shown as mean ± SEM. *p<0.05, **p<0.01, ***p<0.001.

**Extended Data Fig. 3.**
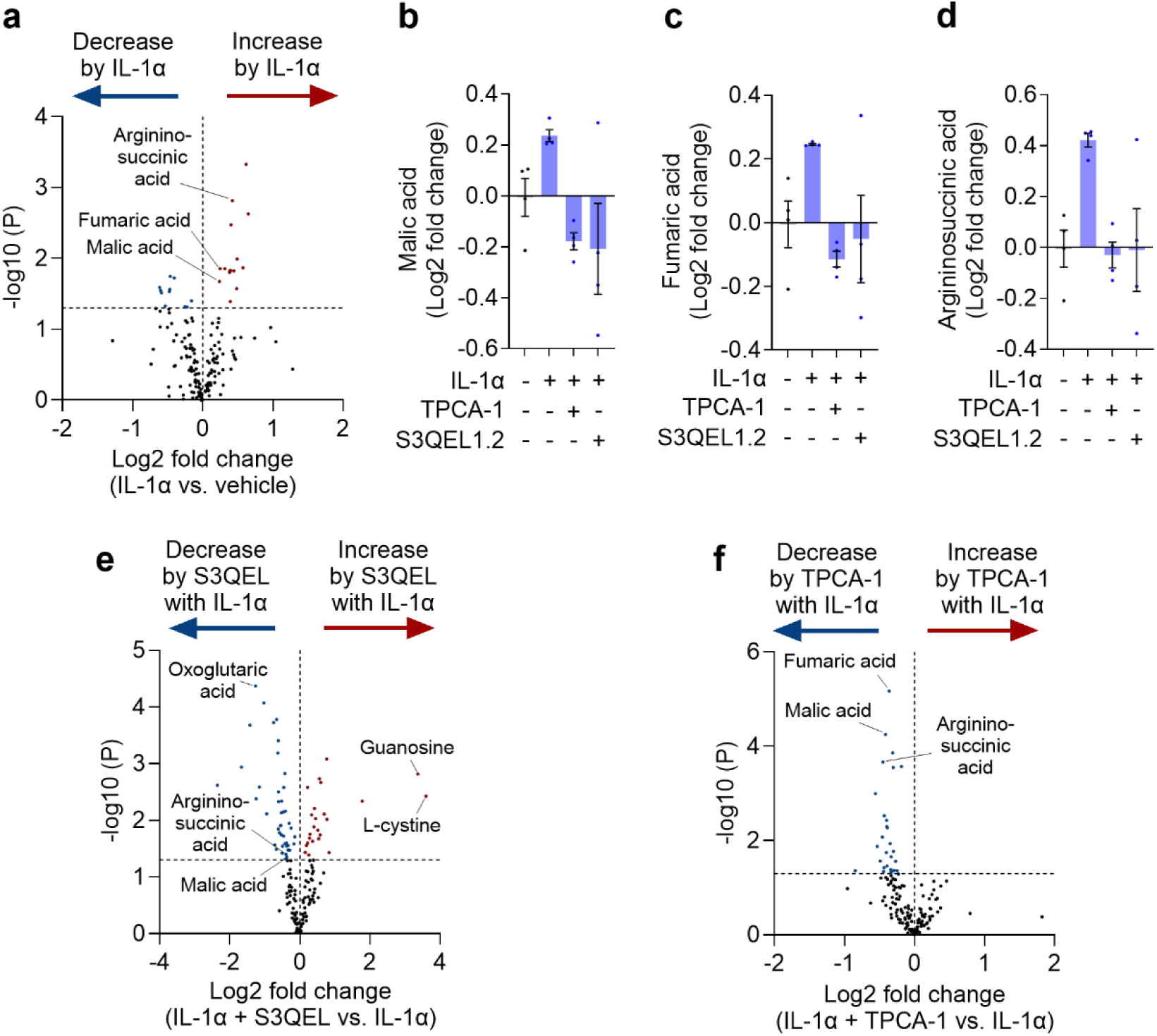
Metabolomic analyses of stimulated astrocytes. a, Metabolite changes after treatment with IL-1α. n = 4 replicates, unpaired t-test. **b**–**d**, Quantification of malic acid (**b**), fumaric acid (**c**), and argininosuccinic acid (**d**) after treatment with vehicle, IL-1α alone or with TPCA-1 or S3QEL1.2. n = 4 replicates. **e,f**, Metabolite changes after treatment with IL-1α alone or with TPCA-1 or S3QEL1.2. n = 4 replicates, unpaired t-test. Data are shown as mean ± SEM.

**Extended Data Fig. 4.**
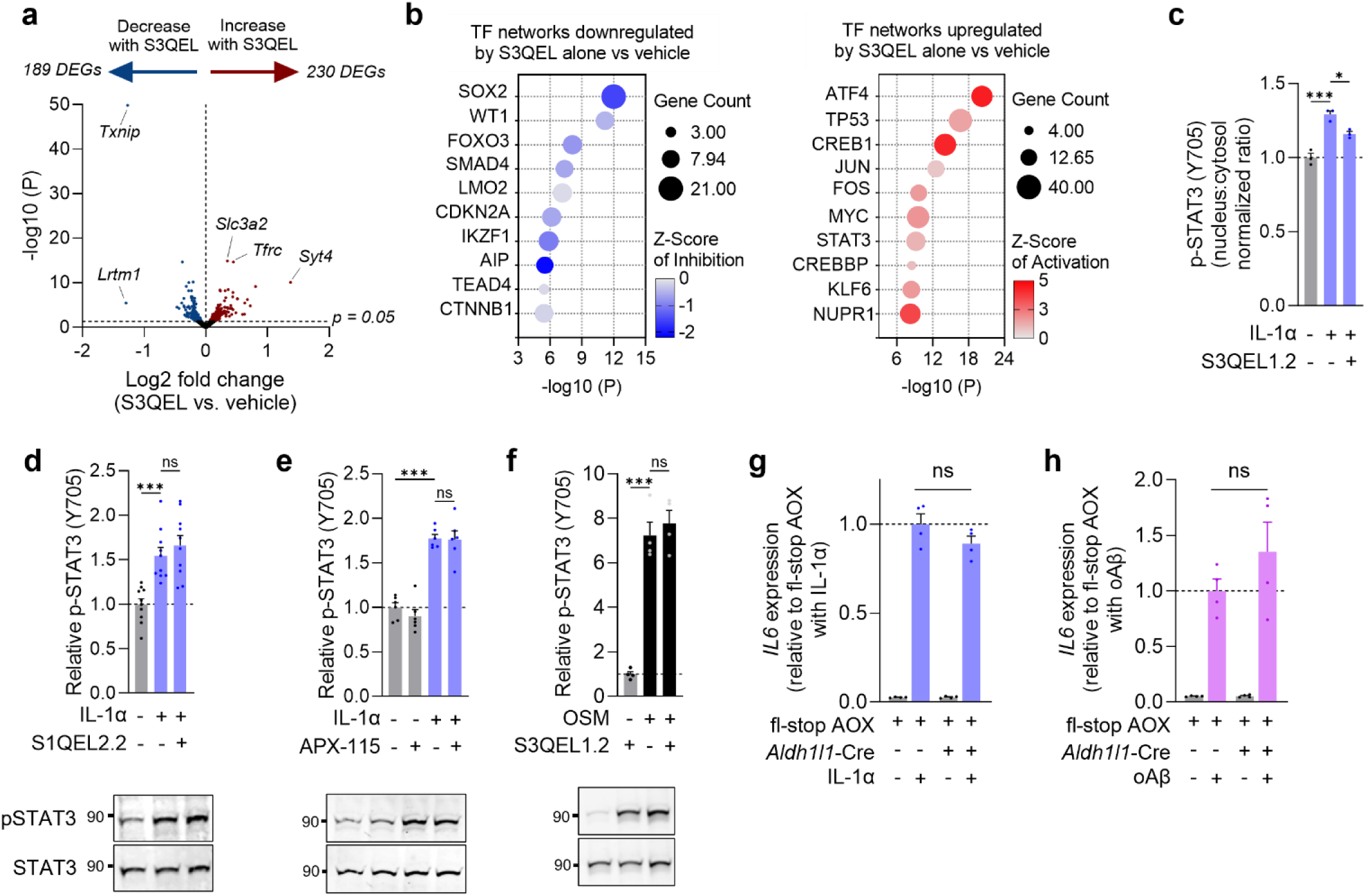
Further characterization of mtROS-dependent STAT3 signaling. a, Volcano plot of gene expression changes after 6 h co-treatment with vehicle or S3QEL1.2 (3 μM). n = 3 replicates. **b**, Transcription factor network analysis of gene expression changes in astrocytes treated with vehicle or S3QEL1.2 for 6 h. n = 3 replicates, listed factors exceed the p<0.05 threshold, Fischer’s exact test. **c**, Quantification of nuclear and cytosolic p-STAT3(Y705) immunofluorescence after 6 h treatment with vehicle, IL-1α, or IL-1α with S3QEL1.2. n = 3 wells, ANOVA with Tukey’s test. **d**–**f**, Representative Western blots and quantification of p-STAT3(Y705) levels after 6 h treatment with vehicle, IL-1α, S3QEL1.2, S1QEL2.2 (10 μM), or APX-115 (5 μM) or for 3 h with oncostatin M (OSM) (10 ng/mL). n = 3–10 wells, ANOVA with Tukey’s test. **g**,**h**, Relative RNA levels of *Il6* in control fl-stop AOX and *Aldh1l1*-Cre; fl-stop AOX astrocytes after 6 h treatment with vehicle, IL-1α, or oAβ n = 4 wells, two-way ANOVA with Bonferroni’s test. Data are shown as mean ± SEM. *p<0.05, **p<0.01, ***p<0.001.

**Extended Data Fig. 5.**
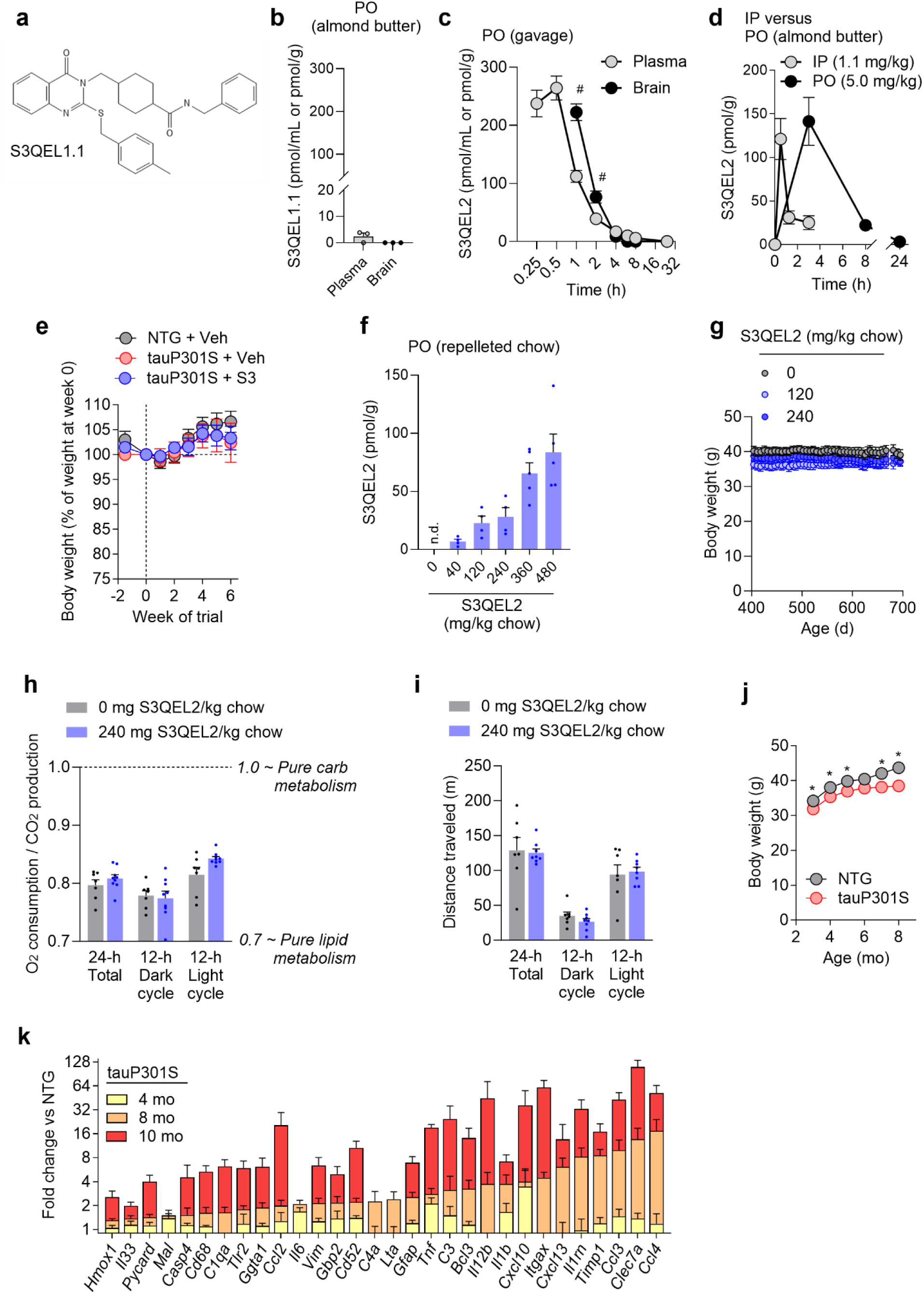
S3QEL2 crosses the blood-brain barrier and is well tolerated with chronic administration in mice. a,b, Plasma and brain levels of S3QEL1.1, a potent analog of S3QEL1.2, after a voluntary oral consumption in almond butter (per oral, PO). S3QEL1.1 was below the limit of quantitation in all brain samples and one plasma sample. n = 3 wild-type mice per group. **c**, Plasma and brain levels of S3QEL2 after a single oral gavage dose of 5 mg/kg. ^#^Brain:Plasma ratios were 1.98 and 1.96 at 1 and 2 h, respectively. n = 3–6 wild-type mice per group. **d**, S3QEL2 brain levels after a single intraperitoneal (IP) injection or voluntary oral consumption in almond butter (per oral, PO). n = 3–6 wild-type mice per group. **e**, Body weights of NTG and P301S male mice treated with S3QEL2 (S3) (5 mg/kg/day, PO in almond butter) or vehicle (Veh) daily for 6 weeks. **f**, S3QEL2 brain levels following chronic oral dosing with formulated chow for 2–3 weeks (n.d. = below limit of detection). n = 4–5 wild-type mice per group. **g**, Body weight of 18–23-month-old wild-type male mice treated with formulated chow starting at 4 mo of age (n = 15 mice per dose). **h,i**, Indirect calorimetry and behavioral tracking in Promethion metabolic cages in 26-month-old male mice after 22 months of chronic S3QEL2 chow administration. n = 7–9 mice per condition. **j**, Body weights of male nontransgenic (NTG) or hTauP301S (P301S) mice. n = 16–26 mice per genotype and time point. **k**, RT-qPCR quantification of neuroinflammation and glial gene expression in tauP301S hippocampus at 4, 8 and 10 mo of age. Genes are rank ordered based on fold changes at 8 mo relative to NTG littermate controls. n = 3–5 mice per genotype and age. Data are shown as mean ± SEM. *p<0.05.

**Extended Data Fig. 6.**
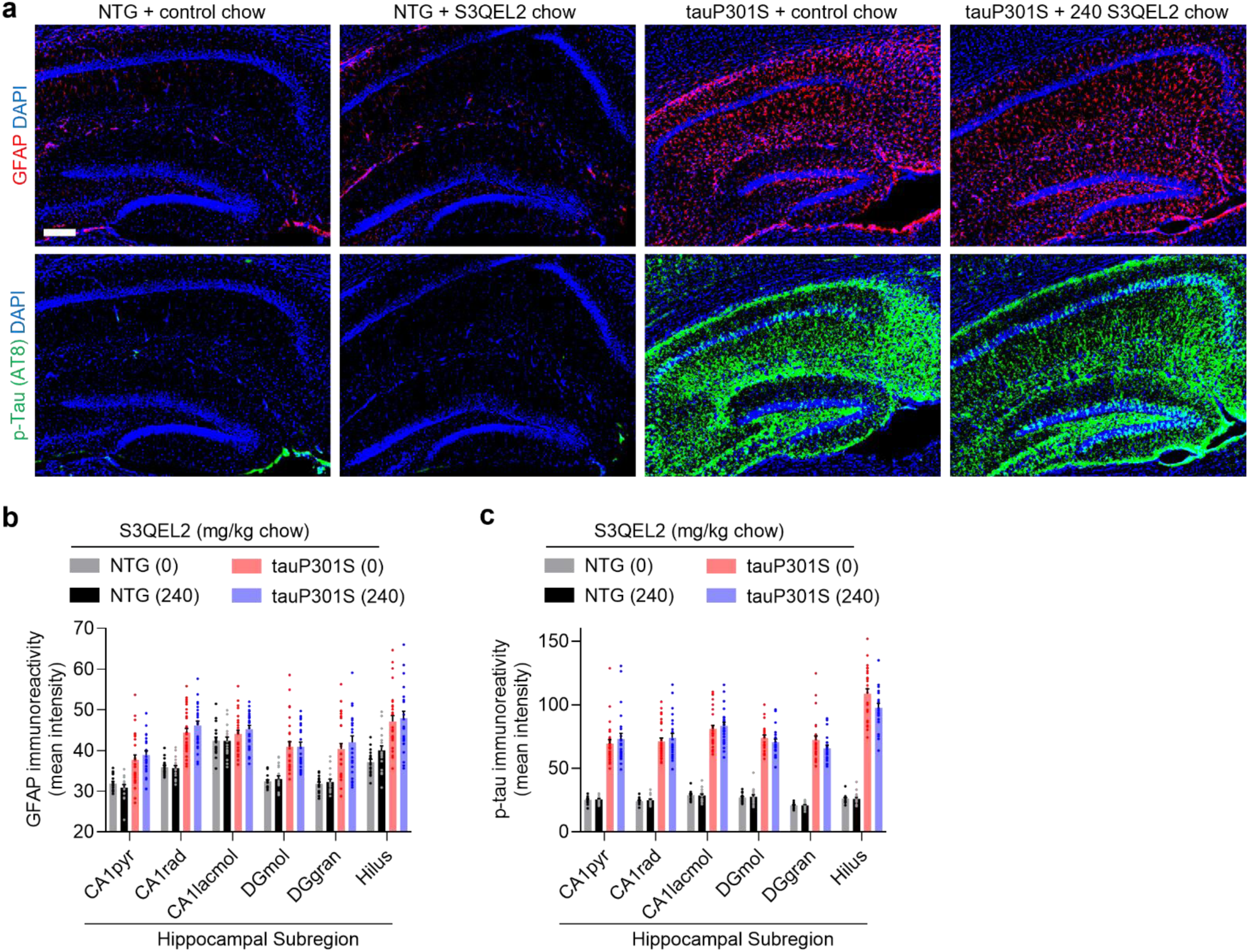
Further characterization of the effects of S3QEL2 in mouse model of tauopathy. a–c, Representative images (**a**) and quantification (**b,c**) of GFAP or p-tau (AT8) immunostaining in hippocampal subregions of tauP301S or NTG mice after 6 months of 0 or 240 mg S3QEL2 per kg chow. n = 9–14 mice per group. CA1pyr, CA1rad, CA1lacmol: pyramidal, radiatum and lacunosum moleculare layers of the Cornu Ammonis subfield 1; DGmol, DGgran: molecular and granular layers of the dentate gyrus. Scale bar: 500 µm. Data are shown as mean ± SEM.

**Extended Data Table 1.**
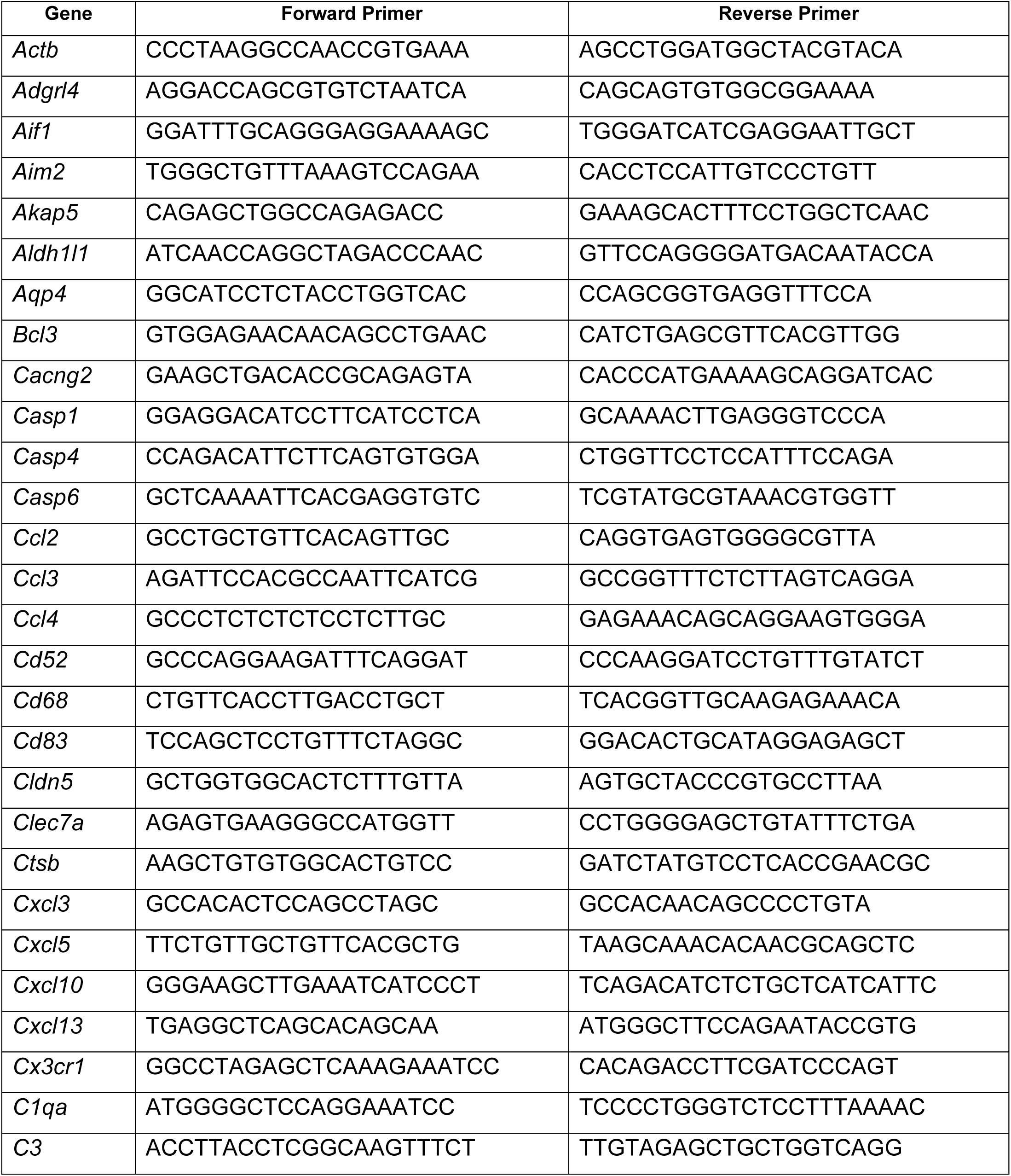
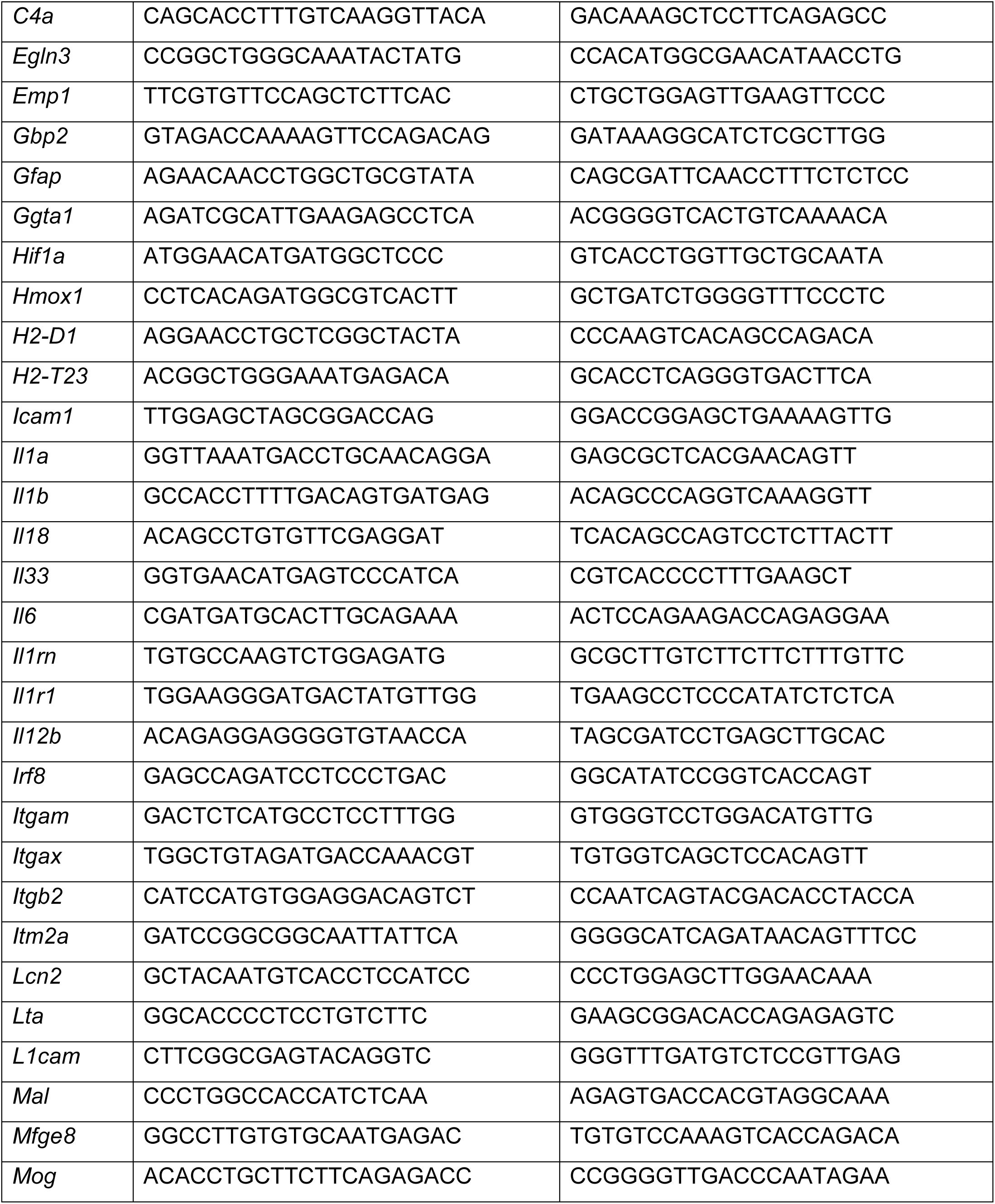
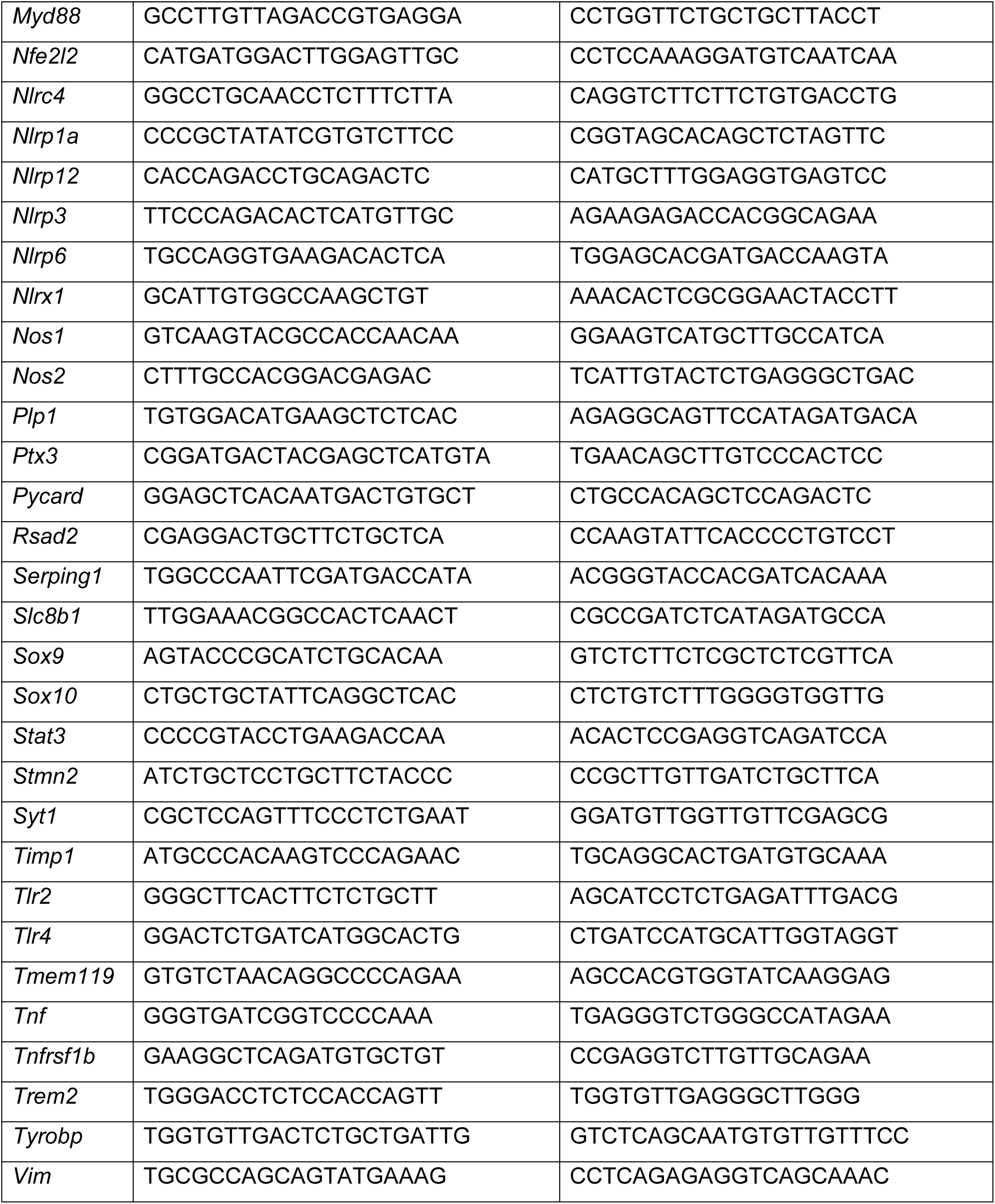
RT-qPCR primers.

**Extended Data Table 2:** Reagent and Resource Information.

| **Software for Data Collection and Analysis** |  |  |
| --- | --- | --- |
| Prism (v 10.1.1) |  |  |
| MaxQuant (v 2.4.2.0) |  |  |
| UniProt mouse protein database (downloaded on 09/21/2017) |  |  |
| Human Protein Atlas |  |  |
| ImageJ2 (v2.3) |  |  |
| Ingenuity Pathway Analysis (IPA) (v01-22-01; data uploaded 2024) |  |  |
| Database for Annotation, Visualization and Integrated Discovery (DAVID) Bioinformatics (v2024q1) |  |  |
| XCalibur (v4.1) |  |  |
| MetaboAnalyst (v6.0) |  |  |
| Image Lab (v5.2.5) |  |  |
| Seahorse Wave Software |  |  |
| DESeq2 package |  |  |
| STAR (v2.5.2) |  |  |
| HTSeq-count (v0.11.2) |  |  |
| Cufflinks (v2.1.1) |  |  |
| Cutadapt (v1.18) |  |  |
| bbcl2fastq (v2.20) |  |  |
| **Antibodies** | **Vendor** | **Identifier** |
| β-actin; rabbit | Millipore-Sigma | Cat#: A2066<br>RRID: AB_476693 |
| ASC; rabbit | Adipogen | Cat#: AG-25B-0006<br>RRID: AB_2490440 |
| CD11b; rat | Bio-Rad | Cat#: MCA711G<br>RRID: AB_323167 |
| Citrate synthase; rabbit | Cell Signaling | Cat#: 14309<br>RRID: AB_2665545 |
| GFAP; rabbit | Millipore-Sigma | Cat#: G9269<br>RRID: AB_477035 |
| GFP; goat | Abcam | Cat#: Ab6658<br>RRID: AB_305631 |
| Glutamine synthetase; rabbit | Abcam | Cat#: Ab16802<br>RRID: AB_302521 |
| HA.11 epitope tag; mouse | BioLegend | Cat#: 901514<br>RRID: AB_2565336 |
| Iba1; rabbit | WAKO | Cat#: 019-19741<br>RRID: AB_839504 |
| MAP2; chicken | Abcam | Cat#: ab5392<br>RRID: AB_2138153 |
| NeuN; rabbit | Millipore-Sigma | Cat#: ABN78<br>RRID: AB_10807945 |
| NF-κB; mouse | Cell Signaling | Cat#: 6956<br>RRID: AB_10828935 |
| Phospho-NF-κB (S536); rabbit | Cell Signaling | Cat#: 3033<br>RRID: AB_331284 |
| OxPhos antibody cocktail; mouse | Abcam | Cat#: Ab110413<br>RRID: AB_2629281 |
| SOX9; rabbit | Millipore-Sigma | Cat#: AB5535<br>RRID: AB_2239761 |
| STAT3; mouse | Cell Signaling | Cat#: 9139<br>RRID: AB_331757 |
| Phospho-STAT3 (Y705); rabbit | Cell Signaling | Cat#: 9145<br>RRID: AB_2491009 |
| Human-specific tau (clone HT7); mouse | ThermoFisher | Cat#: MN1000<br>RRID: AB_2314654 |
| tau (Tau-5); mouse | ThermoFisher | Cat#: AHB0042<br>RRID: AB_2536235 |
| tau (Tau 12); mouse | Millipore-Sigma | Cat#: MAB2241<br>RRID: AB_1977340 |
| Phospho-tau (ser202, thr205) (clone AT8); mouse | ThermoFisher | Cat#: MN1020<br>RRID: AB_223647 |
| γ-tubulin; mouse | Millipore-Sigma | Cat#: T5326<br>RRID: AB_532292 |
| AlexaFluor 488 Donkey anti-mouse | ThermoFisher | Cat#: A21202<br>RRID: AB_141607 |
| AlexaFluor 488 Donkey anti-rabbit | ThermoFisher | Cat#: A21206<br>RRID: AB_2535792 |
| AlexaFluor 555 Goat anti-chicken | ThermoFisher | Cat#: A21437<br>RRID: AB_2535858 |
| AlexaFluor 555 Goat anti-rat | ThermoFisher | Cat#: A21434<br>RRID: AB_141733 |
| AlexaFluor 555 Donkey anti-rabbit | ThermoFisher | Cat#: A31572<br>RRID: AB_162543 |
| AlexaFluor 555 Donkey anti-Goat | ThermoFisher | Cat#: A21432<br>RRID: AB_2535853 |
| AlexaFluor 647 Donkey anti-rabbit | ThermoFisher | Cat#: A31573<br>RRID: AB_2536183 |
| IR Dye 680RD Donkey anti-mouse | LI-COR | Cat#: 926-68072<br>RRID: AB_2814912 |
| IR Dye 800CW Donkey anti-rabbit | LI-COR | Cat#: 926-32213<br>RRID: AB_621848 |
| **Reagents, Biological Materials, and Experimental Models** | **Vendor** | **Identifier** |
| S3QEL2 | Cayman Chemical | 18556 |
| S3QEL2 | WuXi Apptec | Custom Synthesis |
| S3QEL1.2 Peak 1 (*cis* isomer) | WuXi Apptec | Custom Synthesis |
| S3QEL1.1 | Life Chemicals | K284-4711 |
| S1QEL2.2 | Life Chemicals | F2068-0013 |
| Interleukin-1α | Sigma | I3901 |
| IL1Ra | Novus Biologicals | NBP2-35105 |
| APX-115 | Millipore Sigma | ADVH97EBCDFC |
| TPCA-1 | Cayman Chemicals | 15115 |
| IKK-16 | Cayman Chemicals | 13313 |
| CGP 37157 | Cayman Chemicals | 15611 |
| TNFα | Cell Signaling | 8902SF |
| C1q | MyBioSource | MBS147305 |
| IL6 | Biolegend | 575702 |
| Interferon-γ | Cell Signaling | 80385S |
| Lipopolysaccharides from E. coli O111:B4 | Millipore Sigma | L4391 |
| Antimycin A | Millipore Sigma | A8674 |
| Myxothiazol | Millipore Sigma | T5580 |
| Rotenone | Millipore Sigma | R8875 |
| FCCP | Millipore Sigma | C2920 |
| Oligomycin | Millipore Sigma | 75351 |
| BAM15 | Tocris | 5737 |
| Amplex UltraRed | ThermoFisher | A36006 |
| Superoxide dismutase | Millipore Sigma | S7571 |
| Horseradish peroxidase, Type VI | Millipore Sigma | P8375 |
| 6C-Cysteine Phosphonic Acid Tag (6C-CPT) | ThermoFisher | A52285 |
| TMTpro 18-plex | ThermoFisher | A52045 |
| Lys-C, Lysyl Endopeptidase | Wako | 125-05061 |
| Trypsin, Sequencing Grade | Promega | V5113 |
| Lambda Protein Phosphatase | New England Biolabs | P0753 |
| Hydroxylamine | Millipore Sigma | 438227 |
| Bond-Breaker TCEP | ThermoFisher | 77720 |
| Iodoacetamide | Millipore Sigma | I1149 |
| Diethylenetriaminepentaacetic acid (DTPA) | Millipore Sigma | D6518 |
| Neocuproine hydrochloride monohydrate | Millipore Sigma | 72090 |
| Sep-Pak tC18 200 mg | Waters | 186004618 |
| Sep-Pak tC18 50 mg | Waters | WAT054960 |
| Tetramethylrhodamine, Methyl Ester, Perchlorate (TMRM) | ThermoFisher | T668 |
| Lipofectamine 3000 | ThermoFisher | L3000008 |
| 2,2,2-tribromoethanol (Avertin) | Fisher Scientific | AC421430100 |
| Justin's Classic Almond Butter | Amazon.com | B000V79VSY |
| 0.5 oz plastic feeding cups | Amazon | B089D7PKRP |
| Kolliphor EL (Cremophor EL) pH-range 6.0 - 8.0 | Millipore Sigma | C5135 |
| CellTiter-Glo 2.0 | Promega | G9242 |
| DAPI | Millipore Sigma | D9542 |
| Bovine serum albumin | VWR | 97062-904 |
| Bovine serum albumin | Millipore Sigma | A3803 |
| Heat-inactivated FBS | VWR | 89510-188 |
| Normal goat serum | Jackson ImmunoResearch | 005-000-121 |
| Normal donkey serum | Jackson ImmunoResearch | 017-000-121 |
| Paraformaldehyde | VWR | 100504-858 |
| B-27 supplement minus antioxidants | ThermoFisher | 10889038 |
| Poly-D-lysine (75–150 kDa) | Millipore Sigma | P6407 |
| Poly-D-lysine (50–150 kDa) | ThermoFisher | A3890401 |
| Complete Protease Inhibitor Cocktail | Millipore Sigma | 11836153001 |
| Phosphatase Inhibitor Cocktails 2 | Millipore Sigma | P5726 |
| Phosphatase Inhibitor Cocktails 3 | Millipore Sigma | P0044 |
| Seahorse XF Base Medium | Agilent | 103334-100 |
| DMEM without glucose, glutamine, pyruvate, bicarbonate or phenol red | Millipore Sigma | D5030 |
| DMEM with 4.5g/L glucose, without pyruvate and glutamine | Corning | 15-017-CV |
| GlutaMAX | ThermoFisher | 35050061 |
| Sodium Pyruvate | ThermoFisher | 11360070 |
| 70 µm Cell Strainers | VWR | 76327-100 |
| Flex Six Microfluidic Chips | Standard Biotools | 100-6308 |
| 96.96 Dynamic Array | Standard Biotools | BMK-M-96.96 |
| RNase H | New England Biolabs | M0297L |
| PowerUp Sybr Green Master mix for qPCR | ThermoFisher | A25742 |
| Exonuclease I treatment | New England Biolabs | M0293L |
| PreAmp Grandmaster mix | TATAA Biocenter | TA05 |
| SsoFast EvaGreen with Low ROX | BioRad | 1725211 |
| 96-well glass bottom microplates | Greiner | 655891 |
| 10-chamber glass-bottom cell culture slides | Greiner | 543078 |
| 96-well clear microplates | Corning | 9017 |
| 96-well clear bottom microplates | Corning | 3904 |
| Fisherbrand Bead Mill 24 Homogenizer | Fisher Scientific | 15-340-163 |
| Critical commercial assays |  |  |
| High-Select Fe-NTA Phosphopeptide Enrichment Kit | ThermoFisher | A32992 |
| High pH Reversed-Phase Peptide Fractionation Kit | Pierce | 84868 |
| Micro BCA Protein Assay Kit | ThermoFisher | 23235 |
| Detergent-Compatible Bradford Assay | Pierce | 23246 |
| Tris HCl, pH 7.5, for molecular biology | VWR | 10128-696 |
| Ethanol, pure, for molecular biology | Millipore Sigma | E7023 |
| KCl (2 M), RNase-free | ThermoFisher | AM9640G |
| MgCl<sub>2</sub> (1 M) | ThermoFisher | AM9530G |
| DL-Dithiothreitol (DTT) solution, BioUltra, for molecular biology, ~1 M in H<sub>2</sub>O | Millipore Sigma | 43816 |
| Heparin sodium salt from porcine intestinal mucosa | Millipore Sigma | H3393 |
| RNasin Ribonuclease Inhibitor, Recombinant | Promega | N2515 |
| Cycloheximide, crystalline, Ultra-Pure Grade | VWR | 97064-724 |
| Nonidet P-40 Surfact-Amps Detergent Solution; 10% NP-40 | ThermoFisher | 28324 |
| RNeasy Kit | Qiagen | 74106 |
| RNeasy Plus Micro Kit 74034 | Qiagen | 74034 |
| RNase-Free DNase Set | Qiagen | 79256 |
| Protoscript First Strand Synthesis Kit | New England Biolabs | E6300L |
| NEB 5-alpha cells | New England Biolabs | C2987H |
| AAV2/DJ-*hSyn1*-tauP301S-(1N4R)-WPRE-hGHpA | This study | N/A |
| AAV2/PHP.eB-*GfaABC1D*-DIO-Rpl22-3xHA-WPRE-hGHpA | This study | N/A |
| pCS2+MLS-HyPer7 | Pak et al., 2020 | Addgene plasmid: 136470; RRID: Addgene_136470 |
| MAC_C_AOX | Liu et al., 2018 | Addgene plasmid: 111661; RRID: Addgene_111661 |
| pAAV-*GfaABC1D*-NEUROD1-T2A-mCherry | Wang et al., 2021 | Addgene plasmid: 178582; RRID: Addgene_178582 |
| pcDNA3.1+ tau-WT-(1N4R) | Dr. Wenjie Luo | Dr. Wenjie Luo |
| DIO-rM3D(Gs)-mCherry | Dr. Bryan Roth | Addgene plasmid: 50458; RRID: Addgene_50458 |
| pAAV-*GfaABC1D*-mtHyPer7-WPRE-hGHpA | This study | N/A |
| pAAV-*hGfaABC1D*-AOX-T2A-mCherry-WPRE-hGHpA | This study | N/A |
| pAAV-*hGfaABC1D*-AOXmut-T2A-mCherry-WPRE-hGHpA | This study | N/A |
| pAAV-*hSyn1*-tauP301S-(1N4R)-WPRE-hGHpA | This study | N/A |
| pAAV-*GfaABC1D*-DIO-Rpl22-3xHA-WPRE-hGHpA | This study | N/A |
| Mouse: C57BL/6J | Jackson Laboratory | Stock#: 000664; RRID: IMSR_JAC:000664 |
| Mouse: B6; FVB-Tg(*Aldh1l1*-cre)JD1884Htz/J | Jackson Laboratory | Stock #: 023748; RRID: IMSR_JAX:023748 |
| Mouse: B6; C3-Tg(*Prnp*-MAPT*P301S)PS19Vle/J | Jackson Laboratory | Stock#: 008169; RRID: IMSR_JAX:008169 |
| Mouse: B6; SNAPf-AOX | Dr. Alexander Galkin; Dhandapani et al., 2019 | N/A |

## Source Data

Source Data: Original uncropped Western blots

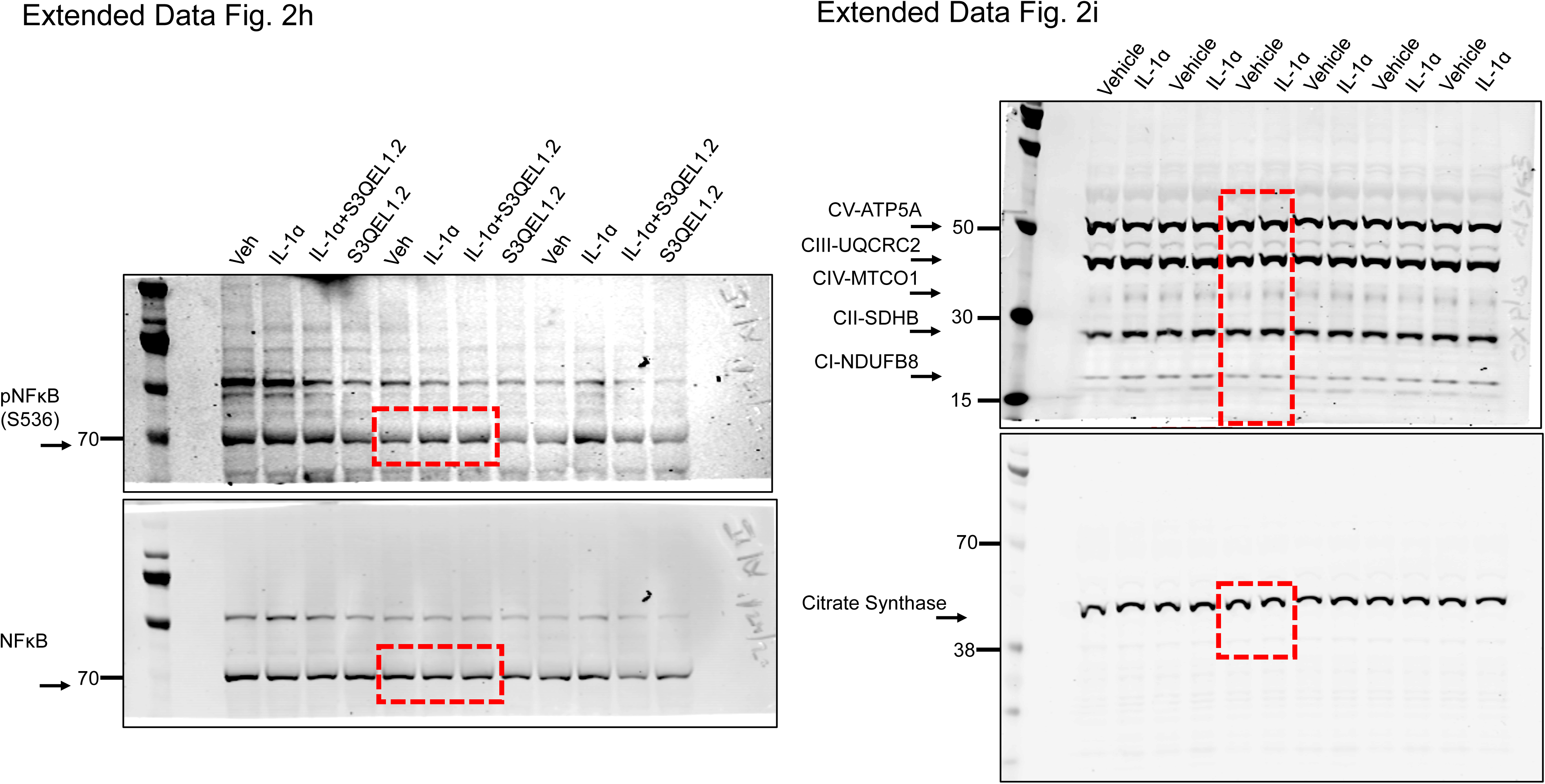
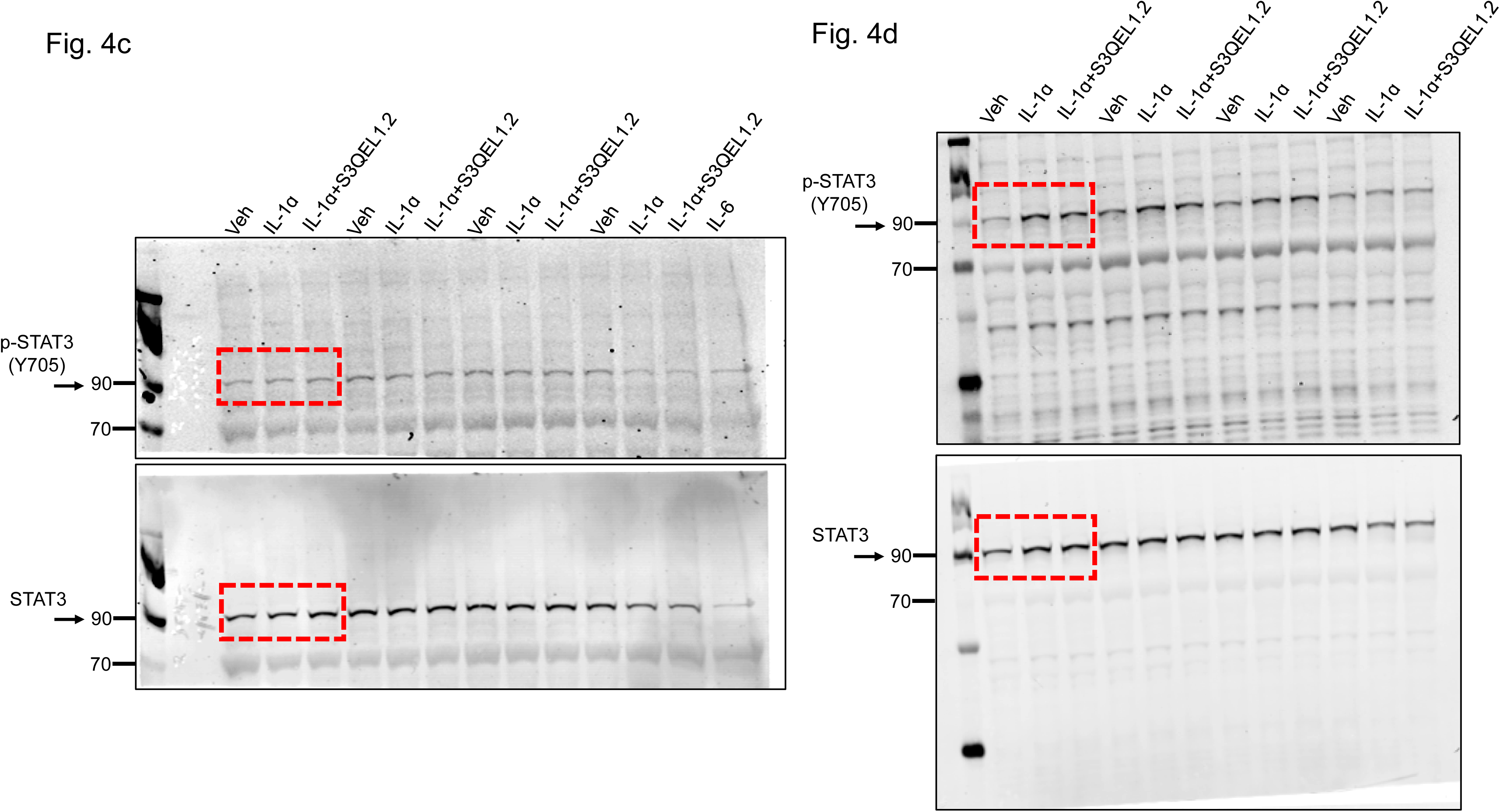
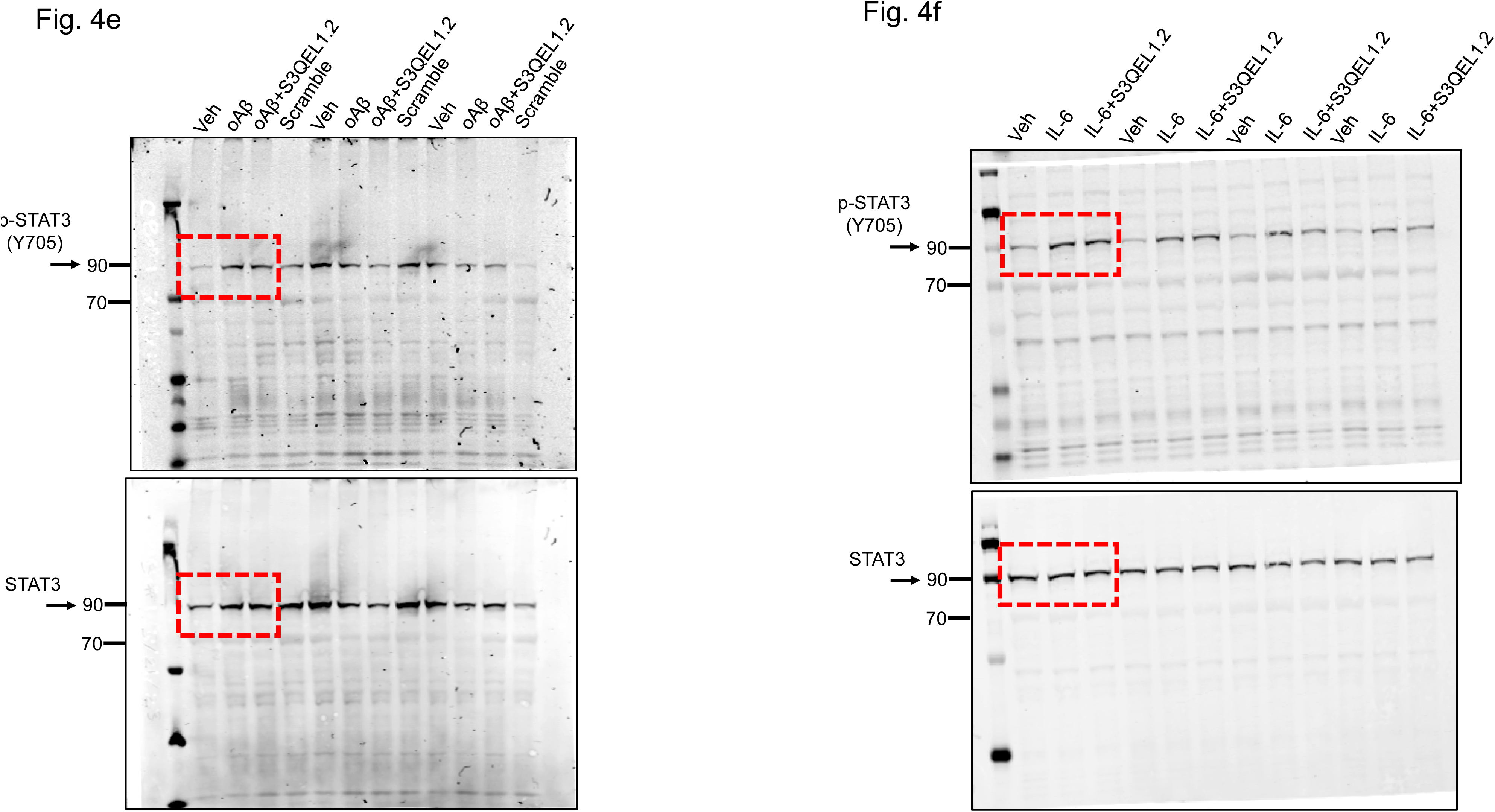
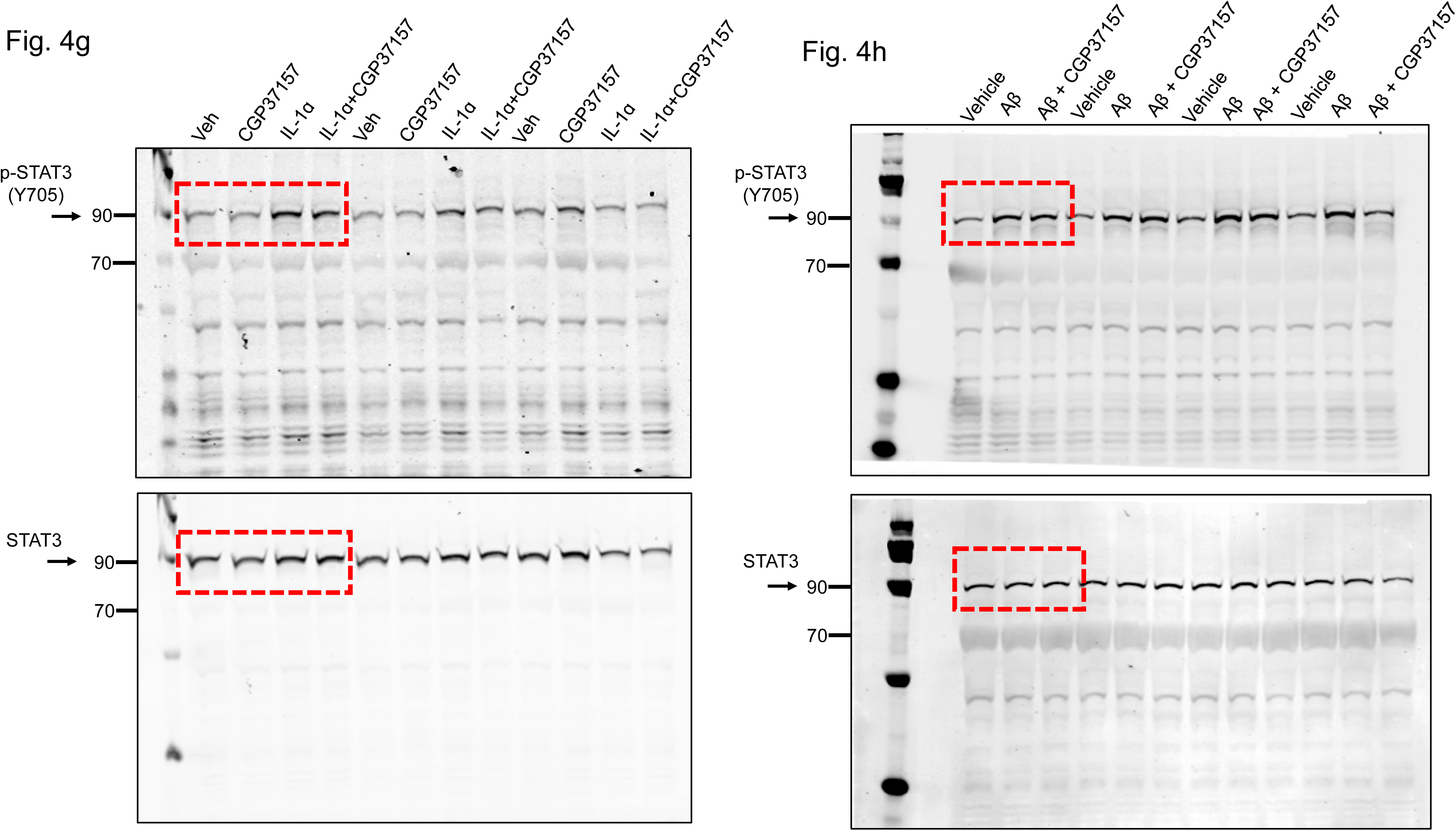
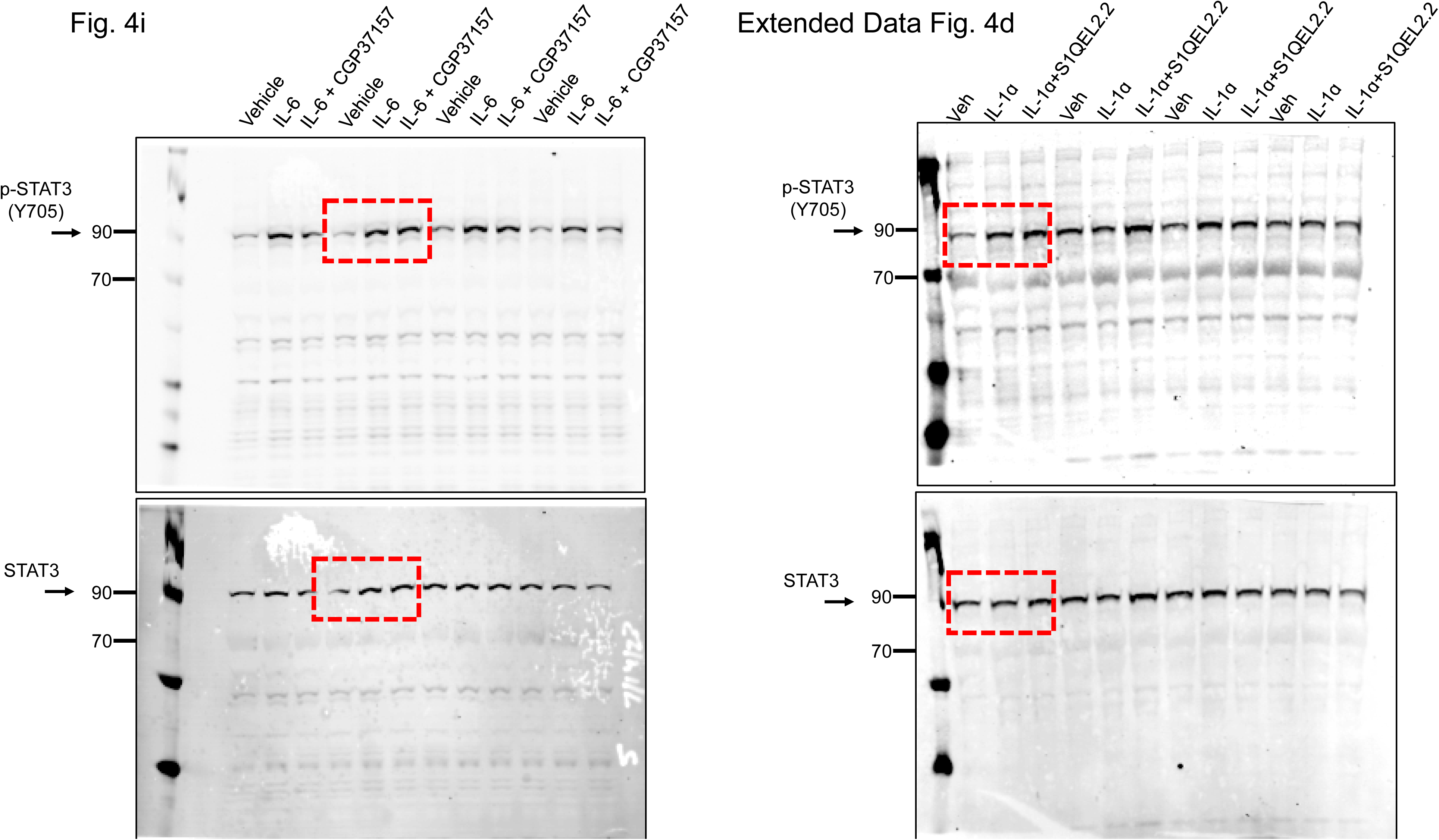
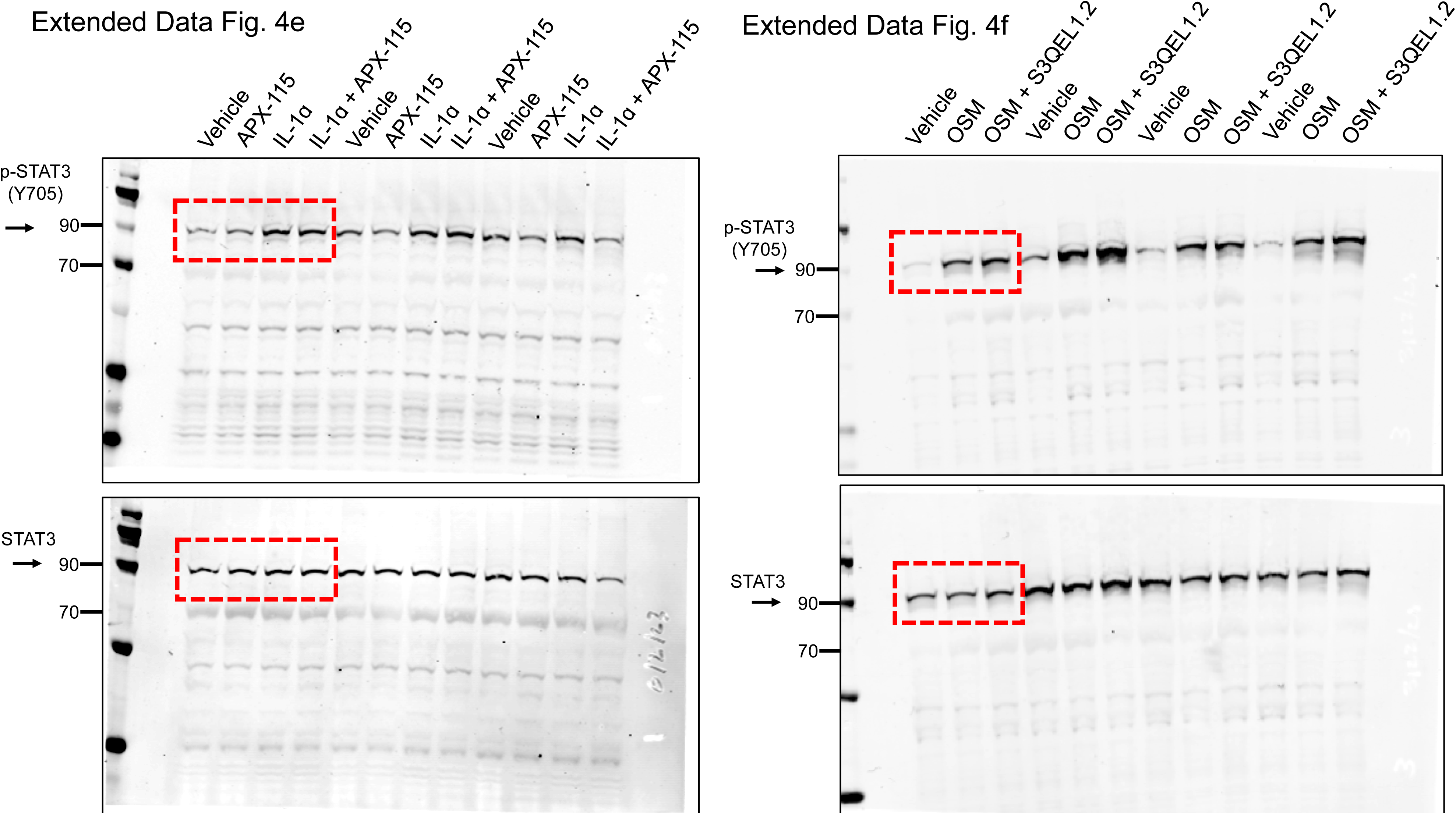

## References

1 Brand, M. D. Mitochondrial generation of superoxide and hydrogen peroxide as the source of mitochondrial redox signaling. Free Radic Biol Med 100, 14–31 (2016). 10.1016/j.freeradbiomed.2016.04.001

2 Guzy, R. D. et al. Mitochondrial complex III is required for hypoxia-induced ROS production and cellular oxygen sensing. Cell Metab 1, 401–408 (2005). 10.1016/j.cmet.2005.05.001

3 Bell, E. L., Klimova, T. A., Eisenbart, J., Schumacker, P. T. & Chandel, N. S. Mitochondrial reactive oxygen species trigger hypoxia-inducible factor-dependent extension of the replicative life span during hypoxia. Mol Cell Biol 27, 5737–5745 (2007). 10.1128/MCB.02265-06

4 Drose, S., Bleier, L. & Brandt, U. A common mechanism links differently acting complex II inhibitors to cardioprotection: modulation of mitochondrial reactive oxygen species production. Mol Pharmacol 79, 814–822 (2011). 10.1124/mol.110.070342

5 Chouchani, E. T. et al. Ischaemic accumulation of succinate controls reperfusion injury through mitochondrial ROS. Nature 515, 431–435 (2014). 10.1038/nature13909

6 Mills, E. L. et al. Succinate Dehydrogenase Supports Metabolic Repurposing of Mitochondria to Drive Inflammatory Macrophages. Cell 167, 457–470 e413 (2016). 10.1016/j.cell.2016.08.064

7 Goncalves, R. L. S., et al. Ubiquinone deficiency drives reverse electron transport to disrupt hepatic metabolic homeostasis in obesity. bioRxiv (2023). 10.1101/2023.02.21.528863

8 Peruzzotti-Jametti, L. et al. Mitochondrial complex I activity in microglia sustains neuroinflammation. Nature 628, 195–203 (2024). 10.1038/s41586-024-07167-9

9 Brand, M. D. The sites and topology of mitochondrial superoxide production. Exp Gerontol 45, 466–472 (2010). 10.1016/j.exger.2010.01.003

10 Zorov, D. B., Juhaszova, M. & Sollott, S. J. Mitochondrial reactive oxygen species (ROS) and ROS-induced ROS release. Physiol Rev 94, 909–950 (2014). 10.1152/physrev.00026.2013

11 Brand, M. D. Riding the tiger - physiological and pathological effects of superoxide and hydrogen peroxide generated in the mitochondrial matrix. Crit Rev Biochem Mol Biol 55, 592–661 (2020). 10.1080/10409238.2020.1828258

12 Orr, A. L. et al. Suppressors of superoxide production from mitochondrial complex III. Nat Chem Biol 11, 834–836 (2015). 10.1038/nchembio.1910

13 Brand, M. D. et al. Suppressors of Superoxide-H(2)O(2) Production at Site I(Q) of Mitochondrial Complex I Protect against Stem Cell Hyperplasia and Ischemia-Reperfusion Injury. Cell Metab 24, 582–592 (2016). 10.1016/j.cmet.2016.08.012

14 Wong, H. S., Benoit, B. & Brand, M. D. Mitochondrial and cytosolic sources of hydrogen peroxide in resting C2C12 myoblasts. Free Radic Biol Med 130, 140–150 (2019). 10.1016/j.freeradbiomed.2018.10.448

15 Plecita-Hlavata, L. et al. Potential of Mitochondria-Targeted Antioxidants to Prevent Oxidative Stress in Pancreatic beta-cells. Oxid Med Cell Longev 2019, 1826303 (2019). 10.1155/2019/1826303

16 Banba, A., Tsuji, A., Kimura, H., Murai, M. & Miyoshi, H. Defining the mechanism of action of S1QELs, specific suppressors of superoxide production in the quinone-reaction site in mitochondrial complex I. J Biol Chem 294, 6550–6561 (2019). 10.1074/jbc.RA119.007687

17 Watson, M. A. et al. S3QELs protect against diet-induced intestinal barrier dysfunction. Aging Cell 20, e13476 (2021). 10.1111/acel.13476

18 Wong, H. S. et al. Superoxide produced by mitochondrial site I(Q) inactivates cardiac succinate dehydrogenase and induces hepatic steatosis in Sod2 knockout mice. Free Radic Biol Med 164, 223–232 (2021). 10.1016/j.freeradbiomed.2020.12.447

19 Watson, M. A. et al. Suppression of superoxide/hydrogen peroxide production at mitochondrial site I(Q) decreases fat accumulation, improves glucose tolerance and normalizes fasting insulin concentration in mice fed a high-fat diet. Free Radic Biol Med 204, 276–286 (2023). 10.1016/j.freeradbiomed.2023.05.022

20 Cozzolino, M., Ferri, A., Valle, C. & Carri, M. T. Mitochondria and ALS: implications from novel genes and pathways. Mol Cell Neurosci 55, 44–49 (2013). 10.1016/j.mcn.2012.06.001

21 Wang, W., Zhao, F., Ma, X., Perry, G. & Zhu, X. Mitochondria dysfunction in the pathogenesis of Alzheimer’s disease: recent advances. Mol Neurodegener 15, 30 (2020). 10.1186/s13024-020-00376-6

22 Wilson, D. M., 3rd et al. Hallmarks of neurodegenerative diseases. Cell 186, 693–714 (2023). 10.1016/j.cell.2022.12.032

23 Rizzuto, R., De Stefani, D., Raffaello, A. & Mammucari, C. Mitochondria as sensors and regulators of calcium signalling. Nat Rev Mol Cell Biol 13, 566–578 (2012). 10.1038/nrm3412

24 Gorlach, A., Bertram, K., Hudecova, S. & Krizanova, O. Calcium and ROS: A mutual interplay. Redox Biol 6, 260–271 (2015). 10.1016/j.redox.2015.08.010

25 Cenini, G. & Voos, W. Role of Mitochondrial Protein Quality Control in Oxidative Stress-induced Neurodegenerative Diseases. Curr Alzheimer Res 13, 164–173 (2016). 10.2174/1567205012666150921103213

26 Joshi, A. U. & Mochly-Rosen, D. Mortal engines: Mitochondrial bioenergetics and dysfunction in neurodegenerative diseases. Pharmacol Res 138, 2–15 (2018). 10.1016/j.phrs.2018.08.010

27 Verma, M., Lizama, B. N. & Chu, C. T. Excitotoxicity, calcium and mitochondria: a triad in synaptic neurodegeneration. Transl Neurodegener 11, 3 (2022). 10.1186/s40035-021-00278-7

28 Trigo, D., Avelar, C., Fernandes, M., Sa, J. & da Cruz, E. S. O. Mitochondria, energy, and metabolism in neuronal health and disease. FEBS Lett 596, 1095–1110 (2022). 10.1002/1873-3468.14298

29 Bordt, E. A. & Polster, B. M. NADPH oxidase- and mitochondria-derived reactive oxygen species in proinflammatory microglial activation: a bipartisan affair? Free Radic Biol Med 76, 34–46 (2014). 10.1016/j.freeradbiomed.2014.07.033

30 Lin, C. H. et al. Mitochondrial UQCRC1 mutations cause autosomal dominant parkinsonism with polyneuropathy. Brain 143, 3352–3373 (2020). 10.1093/brain/awaa279

31 Diaz, F., Garcia, S., Padgett, K. R. & Moraes, C. T. A defect in the mitochondrial complex III, but not complex IV, triggers early ROS-dependent damage in defined brain regions. Hum Mol Genet 21, 5066–5077 (2012). 10.1093/hmg/dds350

32 Gibson, G. E., Starkov, A., Blass, J. P., Ratan, R. R. & Beal, M. F. Cause and consequence: mitochondrial dysfunction initiates and propagates neuronal dysfunction, neuronal death and behavioral abnormalities in age-associated neurodegenerative diseases. Biochim Biophys Acta 1802, 122–134 (2010). 10.1016/j.bbadis.2009.08.010

33 Choi, S. Y. et al. C9ORF72-ALS/FTD-associated poly(GR) binds Atp5a1 and compromises mitochondrial function in vivo. Nat Neurosci 22, 851–862 (2019). 10.1038/s41593-019-0397-0

34 Johri, A. & Beal, M. F. Mitochondrial dysfunction in neurodegenerative diseases. J Pharmacol Exp Ther 342, 619–630 (2012). 10.1124/jpet.112.192138

35 Orr, A. L. et al. Neuronal Apolipoprotein E4 Expression Results in Proteome-Wide Alterations and Compromises Bioenergetic Capacity by Disrupting Mitochondrial Function. J Alzheimers Dis 68, 991–1011 (2019). 10.3233/JAD-181184

36 Orr, A. L. et al. N-terminal mutant huntingtin associates with mitochondria and impairs mitochondrial trafficking. J Neurosci 28, 2783–2792 (2008). 10.1523/JNEUROSCI.0106-08.2008

37 Orr, A. L. et al. Long-term oral kinetin does not protect against alpha-synuclein-induced neurodegeneration in rodent models of Parkinson’s disease. Neurochem Int 109, 106–116 (2017). 10.1016/j.neuint.2017.04.006

38 Wang, P. et al. TDP-43 induces mitochondrial damage and activates the mitochondrial unfolded protein response. PLoS Genet 15, e1007947 (2019). 10.1371/journal.pgen.1007947

39 Walker, J. M. et al. Differential protein expression in the hippocampi of resilient individuals identified by digital spatial profiling. Acta Neuropathol Commun 10, 23 (2022). 10.1186/s40478-022-01324-9

40 Brandebura, A. N., Paumier, A., Onur, T. S. & Allen, N. J. Astrocyte contribution to dysfunction, risk and progression in neurodegenerative disorders. Nat Rev Neurosci 24, 23–39 (2023). 10.1038/s41583-022-00641-1

41 Lee, H. G., Lee, J. H., Flausino, L. E. & Quintana, F. J. Neuroinflammation: An astrocyte perspective. Sci Transl Med 15, eadi7828 (2023). 10.1126/scitranslmed.adi7828

42 Patani, R., Hardingham, G. E. & Liddelow, S. A. Functional roles of reactive astrocytes in neuroinflammation and neurodegeneration. Nat Rev Neurol 19, 395–409 (2023). 10.1038/s41582-023-00822-1

43 Zimmer, T. S., Orr, A. L. & Orr, A. G. Astrocytes in selective vulnerability to neurodegenerative disease. Trends Neurosci 47, 289–302 (2024). 10.1016/j.tins.2024.02.008

44 Bellaver, B. et al. Astrocyte reactivity influences amyloid-beta effects on tau pathology in preclinical Alzheimer’s disease. Nat Med 29, 1775–1781 (2023). 10.1038/s41591-023-02380-x

45 Lopez-Fabuel, I. et al. Complex I assembly into supercomplexes determines differential mitochondrial ROS production in neurons and astrocytes. Proc Natl Acad Sci U S A 113, 13063–13068 (2016). 10.1073/pnas.1613701113

46 Bonvento, G. & Bolanos, J. P. Astrocyte-neuron metabolic cooperation shapes brain activity. Cell Metab 33, 1546–1564 (2021). 10.1016/j.cmet.2021.07.006

47 Fernandez-Fernandez, S., Almeida, A. & Bolanos, J. P. Antioxidant and bioenergetic coupling between neurons and astrocytes. Biochem J 443, 3–11 (2012). 10.1042/BJ20111943

48 Jiwaji, Z. & Hardingham, G. E. The consequences of neurodegenerative disease on neuron-astrocyte metabolic and redox interactions. Neurobiol Dis 185, 106255 (2023). 10.1016/j.nbd.2023.106255

49 Vicente-Gutierrez, C. et al. Astrocytic mitochondrial ROS modulate brain metabolism and mouse behaviour. Nat Metab 1, 201–211 (2019). 10.1038/s42255-018-0031-6

50 Cassina, P. et al. Mitochondrial dysfunction in SOD1G93A-bearing astrocytes promotes motor neuron degeneration: prevention by mitochondrial-targeted antioxidants. J Neurosci 28, 4115–4122 (2008). 10.1523/JNEUROSCI.5308-07.2008

51 Chun, H. et al. Severe reactive astrocytes precipitate pathological hallmarks of Alzheimer’s disease via H(2)O(2)(-) production. Nat Neurosci 23, 1555–1566 (2020). 10.1038/s41593-020-00735-y

52 Polyzos, A. A. et al. Metabolic Reprogramming in Astrocytes Distinguishes Region-Specific Neuronal Susceptibility in Huntington Mice. Cell Metab 29, 1258–1273 e1211 (2019). 10.1016/j.cmet.2019.03.004

53 Liddelow, S. A. et al. Neurotoxic reactive astrocytes are induced by activated microglia. Nature 541, 481–487 (2017). 10.1038/nature21029

54 Bretheau, F. et al. The alarmin interleukin-1alpha triggers secondary degeneration through reactive astrocytes and endothelium after spinal cord injury. Nat Commun 13, 5786 (2022). 10.1038/s41467-022-33463-x

55 Italiani, P. et al. Circulating levels of IL-1 family cytokines and receptors in Alzheimer’s disease: new markers of disease progression? J Neuroinflammation 15, 342 (2018). 10.1186/s12974-018-1376-1

56 Lyra, E. S. N. M. et al. Pro-inflammatory interleukin-6 signaling links cognitive impairments and peripheral metabolic alterations in Alzheimer’s disease. Transl Psychiatry 11, 251 (2021). 10.1038/s41398-021-01349-z

57 Rothaug, M., Becker-Pauly, C. & Rose-John, S. The role of interleukin-6 signaling in nervous tissue. Biochim Biophys Acta 1863, 1218–1227 (2016). 10.1016/j.bbamcr.2016.03.018

58 Licht-Murava, A. et al. Astrocytic TDP-43 dysregulation impairs memory by modulating antiviral pathways and interferon-inducible chemokines. Sci Adv 9, eade1282 (2023). 10.1126/sciadv.ade1282

59 Fang, J., Wong, H. S. & Brand, M. D. Production of superoxide and hydrogen peroxide in the mitochondrial matrix is dominated by site I(Q) of complex I in diverse cell lines. Redox Biol 37, 101722 (2020). 10.1016/j.redox.2020.101722

60 Pak, V. V. et al. Ultrasensitive Genetically Encoded Indicator for Hydrogen Peroxide Identifies Roles for the Oxidant in Cell Migration and Mitochondrial Function. Cell Metab 31, 642–653 e646 (2020). 10.1016/j.cmet.2020.02.003

61 Loschen, G., Flohe, L. & Chance, B. Respiratory chain linked H(2)O(2) production in pigeon heart mitochondria. FEBS Lett 18, 261–264 (1971). 10.1016/0014-5793(71)80459-3

62 Boveris, A. & Chance, B. The mitochondrial generation of hydrogen peroxide. General properties and effect of hyperbaric oxygen. Biochem J 134, 707–716 (1973). 10.1042/bj1340707

63 De Strooper, B. & Karran, E. The Cellular Phase of Alzheimer’s Disease. Cell 164, 603–615 (2016). 10.1016/j.cell.2015.12.056

64 Narayan, P. et al. Rare individual amyloid-beta oligomers act on astrocytes to initiate neuronal damage. Biochemistry 53, 2442–2453 (2014). 10.1021/bi401606f

65 Ma, T. et al. Amyloid beta-induced impairments in hippocampal synaptic plasticity are rescued by decreasing mitochondrial superoxide. J Neurosci 31, 5589–5595 (2011). 10.1523/JNEUROSCI.6566-10.2011

66 Garwood, C. J., Pooler, A. M., Atherton, J., Hanger, D. P. & Noble, W. Astrocytes are important mediators of Abeta-induced neurotoxicity and tau phosphorylation in primary culture. Cell Death Dis 2, e167 (2011). 10.1038/cddis.2011.50

67 El-Khoury, R. et al. Engineering the alternative oxidase gene to better understand and counteract mitochondrial defects: state of the art and perspectives. Br J Pharmacol 171, 2243–2249 (2014). 10.1111/bph.12570

68 Sommer, N. et al. Bypassing mitochondrial complex III using alternative oxidase inhibits acute pulmonary oxygen sensing. Sci Adv 6, eaba0694 (2020). 10.1126/sciadv.aba0694

69 Billingham, L. K. et al. Mitochondrial electron transport chain is necessary for NLRP3 inflammasome activation. Nat Immunol 23, 692–704 (2022). 10.1038/s41590-022-01185-3

70 Roca, F. J., Whitworth, L. J., Prag, H. A., Murphy, M. P. & Ramakrishnan, L. Tumor necrosis factor induces pathogenic mitochondrial ROS in tuberculosis through reverse electron transport. Science 376, eabh2841 (2022). 10.1126/science.abh2841

71 Herrero Martin, J. C., et al. An ETFDH-driven metabolon supports OXPHOS efficiency in skeletal muscle by regulating coenzyme Q homeostasis. Nat Metab 6, 209–225 (2024). 10.1038/s42255-023-00956-y

72 Andjelkovic, A. et al. Diiron centre mutations in Ciona intestinalis alternative oxidase abolish enzymatic activity and prevent rescue of cytochrome oxidase deficiency in flies. Sci Rep 5, 18295 (2015). 10.1038/srep18295

73 Gajtko, A., Bakk, E., Hegedus, K., Ducza, L. & Hollo, K. IL-1beta Induced Cytokine Expression by Spinal Astrocytes Can Play a Role in the Maintenance of Chronic Inflammatory Pain. Front Physiol 11, 543331 (2020). 10.3389/fphys.2020.543331

74 Dinarello, C. A. Overview of the IL-1 family in innate inflammation and acquired immunity. Immunol Rev 281, 8–27 (2018). 10.1111/imr.12621

75 Dresselhaus, E. C. & Meffert, M. K. Cellular Specificity of NF-kappaB Function in the Nervous System. Front Immunol 10, 1043 (2019). 10.3389/fimmu.2019.01043

76 Hernansanz-Agustin, P. et al. Na(+) controls hypoxic signalling by the mitochondrial respiratory chain. Nature 586, 287–291 (2020). 10.1038/s41586-020-2551-y

77 Cox, D. A., Conforti, L., Sperelakis, N. & Matlib, M. A. Selectivity of inhibition of Na(+)-Ca2+ exchange of heart mitochondria by benzothiazepine CGP-37157. J Cardiovasc Pharmacol 21, 595–599 (1993). 10.1097/00005344-199304000-00013

78 Cabral-Costa, J. V. et al. Mitochondrial sodium/calcium exchanger NCLX regulates glycolysis in astrocytes, impacting on cognitive performance. J Neurochem 165, 521–535 (2023). 10.1111/jnc.15745

79 Parnis, J. et al. Mitochondrial exchanger NCLX plays a major role in the intracellular Ca2+ signaling, gliotransmission, and proliferation of astrocytes. J Neurosci 33, 7206–7219 (2013). 10.1523/JNEUROSCI.5721-12.2013

80 Kostic, M., Katoshevski, T. & Sekler, I. Allosteric Regulation of NCLX by Mitochondrial Membrane Potential Links the Metabolic State and Ca(2+) Signaling in Mitochondria. Cell Rep 25, 3465–3475 e3464 (2018). 10.1016/j.celrep.2018.11.084

81 Boyman, L., Williams, G. S., Khananshvili, D., Sekler, I. & Lederer, W. J. NCLX: the mitochondrial sodium calcium exchanger. J Mol Cell Cardiol 59, 205–213 (2013). 10.1016/j.yjmcc.2013.03.012

82 Kenwood, B. M. et al. Identification of a novel mitochondrial uncoupler that does not depolarize the plasma membrane. Mol Metab 3, 114–123 (2014). 10.1016/j.molmet.2013.11.005

83 Hamanaka, R. B. & Chandel, N. S. Mitochondrial reactive oxygen species regulate cellular signaling and dictate biological outcomes. Trends Biochem Sci 35, 505–513 (2010). 10.1016/j.tibs.2010.04.002

84 Bleier, L. & Drose, S. Superoxide generation by complex III: from mechanistic rationales to functional consequences. Biochim Biophys Acta 1827, 1320–1331 (2013). 10.1016/j.bbabio.2012.12.002

85 Bleier, L. et al. Generator-specific targets of mitochondrial reactive oxygen species. Free Radic Biol Med 78, 1–10 (2015). 10.1016/j.freeradbiomed.2014.10.511

86 Xiao, H. et al. A Quantitative Tissue-Specific Landscape of Protein Redox Regulation during Aging. Cell 180, 968–983 e924 (2020). 10.1016/j.cell.2020.02.012

87 Chouchani, E. T. et al. Mitochondrial ROS regulate thermogenic energy expenditure and sulfenylation of UCP1. Nature 532, 112–116 (2016). 10.1038/nature17399

88 Burda, J. E. et al. Divergent transcriptional regulation of astrocyte reactivity across disorders. Nature 606, 557–564 (2022). 10.1038/s41586-022-04739-5

89 Ben Haim, L., et al. The JAK/STAT3 pathway is a common inducer of astrocyte reactivity in Alzheimer’s and Huntington’s diseases. J Neurosci 35, 2817–2829 (2015). 10.1523/JNEUROSCI.3516-14.2015

90 Ceyzeriat, K. et al. Modulation of astrocyte reactivity improves functional deficits in mouse models of Alzheimer’s disease. Acta Neuropathol Commun 6, 104 (2018). 10.1186/s40478-018-0606-1

91 Choi, M., Kim, H., Yang, E. J. & Kim, H. S. Inhibition of STAT3 phosphorylation attenuates impairments in learning and memory in 5XFAD mice, an animal model of Alzheimer’s disease. J Pharmacol Sci 143, 290–299 (2020). 10.1016/j.jphs.2020.05.009

92 Mehla, J. et al. STAT3 inhibitor mitigates cerebral amyloid angiopathy and parenchymal amyloid plaques while improving cognitive functions and brain networks. Acta Neuropathol Commun 9, 193 (2021). 10.1186/s40478-021-01293-5

93 Reichenbach, N. et al. Inhibition of Stat3-mediated astrogliosis ameliorates pathology in an Alzheimer’s disease model. EMBO Mol Med 11 (2019). 10.15252/emmm.201809665

94 Roberts, J. A. et al. A brain proteomic signature of incipient Alzheimer’s disease in young APOE epsilon4 carriers identifies novel drug targets. Sci Adv 7, eabi8178 (2021). 10.1126/sciadv.abi8178

95 Santiago, J. A., Bottero, V. & Potashkin, J. A. Transcriptomic and Network Analysis Identifies Shared and Unique Pathways across Dementia Spectrum Disorders. Int J Mol Sci 21 (2020). 10.3390/ijms21062050

96 Leng, K. et al. CRISPRi screens in human iPSC-derived astrocytes elucidate regulators of distinct inflammatory reactive states. Nat Neurosci 25, 1528–1542 (2022). 10.1038/s41593-022-01180-9

97 Goncalves, R. L., Quinlan, C. L., Perevoshchikova, I. V., Hey-Mogensen, M. & Brand, M. D. Sites of superoxide and hydrogen peroxide production by muscle mitochondria assessed ex vivo under conditions mimicking rest and exercise. J Biol Chem 290, 209–227 (2015). 10.1074/jbc.M114.619072

98 Tabata Fukushima, C., et al. Reactive oxygen species generation by reverse electron transfer at mitochondrial complex I under simulated early reperfusion conditions. Redox Biol 70, 103047 (2024). 10.1016/j.redox.2024.103047

99 Akira, S. et al. Molecular cloning of APRF, a novel IFN-stimulated gene factor 3 p91-related transcription factor involved in the gp130-mediated signaling pathway. Cell 77, 63–71 (1994). 10.1016/0092-8674(94)90235-6

100 Hillmer, E. J., Zhang, H., Li, H. S. & Watowich, S. S. STAT3 signaling in immunity. Cytokine Growth Factor Rev 31, 1–15 (2016). 10.1016/j.cytogfr.2016.05.001

101 Liu, X. et al. Stat3 inhibition attenuates mechanical allodynia through transcriptional regulation of chemokine expression in spinal astrocytes. PLoS One 8, e75804 (2013). 10.1371/journal.pone.0075804

102 Traber, K. E. et al. Induction of STAT3-Dependent CXCL5 Expression and Neutrophil Recruitment by Oncostatin-M during Pneumonia. Am J Respir Cell Mol Biol 53, 479–488 (2015). 10.1165/rcmb.2014-0342OC

103 Fogli, L. K. et al. T cell-derived IL-17 mediates epithelial changes in the airway and drives pulmonary neutrophilia. J Immunol 191, 3100–3111 (2013). 10.4049/jimmunol.1301360

104 Puram, S. V. et al. STAT3-iNOS Signaling Mediates EGFRvIII-Induced Glial Proliferation and Transformation. J Neurosci 32, 7806–7818 (2012). 10.1523/JNEUROSCI.3243-11.2012

105 Anderson, M. A. et al. Astrocyte scar formation aids central nervous system axon regeneration. Nature 532, 195–200 (2016). 10.1038/nature17623

106 Litvinchuk, A. et al. Complement C3aR Inactivation Attenuates Tau Pathology and Reverses an Immune Network Deregulated in Tauopathy Models and Alzheimer’s Disease. Neuron 100, 1337–1353 e1335 (2018). 10.1016/j.neuron.2018.10.031

107 Liu, R. et al. NOX activation in reactive astrocytes regulates astrocytic LCN2 expression and neurodegeneration. Cell Death Dis 13, 371 (2022). 10.1038/s41419-022-04831-8

108 Wan, C. K. et al. Opposing roles of STAT1 and STAT3 in IL-21 function in CD4+ T cells. Proc Natl Acad Sci U S A 112, 9394–9399 (2015). 10.1073/pnas.1511711112

109 Yang, H. et al. STAT3 Inhibition Enhances the Therapeutic Efficacy of Immunogenic Chemotherapy by Stimulating Type 1 Interferon Production by Cancer Cells. Cancer Res 75, 3812–3822 (2015). 10.1158/0008-5472.CAN-15-1122

110 Bi, F. et al. Reactive astrocytes secrete lcn2 to promote neuron death. Proc Natl Acad Sci U S A 110, 4069–4074 (2013). 10.1073/pnas.1218497110

111 Avalle, L. et al. STAT3 induces breast cancer growth via ANGPTL4, MMP13 and STC1 secretion by cancer associated fibroblasts. Oncogene 41, 1456–1467 (2022). 10.1038/s41388-021-02172-y

112 Kim, J., Lim, J., Yoo, I. D., Park, S. & Moon, J. S. TXNIP contributes to induction of pro-inflammatory phenotype and caspase-3 activation in astrocytes during Alzheimer’s diseases. Redox Biol 63, 102735 (2023). 10.1016/j.redox.2023.102735

113 Nishiyama, A. et al. Identification of thioredoxin-binding protein-2/vitamin D(3) up-regulated protein 1 as a negative regulator of thioredoxin function and expression. J Biol Chem 274, 21645–21650 (1999). 10.1074/jbc.274.31.21645

114 Bugiani, O. et al. Frontotemporal dementia and corticobasal degeneration in a family with a P301S mutation in tau. J Neuropathol Exp Neurol 58, 667–677 (1999). 10.1097/00005072-199906000-00011

115 Sperfeld, A. D. et al. FTDP-17: an early-onset phenotype with parkinsonism and epileptic seizures caused by a novel mutation. Ann Neurol 46, 708–715 (1999). 10.1002/1531-8249(199911)46:5<708::aid-ana5>3.0.co;2-k

116 Chang, C. W., Evans, M. D., Yu, X., Yu, G. Q. & Mucke, L. Tau reduction affects excitatory and inhibitory neurons differently, reduces excitation/inhibition ratios, and counteracts network hypersynchrony. Cell Rep 37, 109855 (2021). 10.1016/j.celrep.2021.109855

117 Yoshiyama, Y. et al. Synapse loss and microglial activation precede tangles in a P301S tauopathy mouse model. Neuron 53, 337–351 (2007). 10.1016/j.neuron.2007.01.010

118 Jiwaji, Z. et al. Reactive astrocytes acquire neuroprotective as well as deleterious signatures in response to Tau and Abeta pathology. Nat Commun 13, 135 (2022). 10.1038/s41467-021-27702-w

119 Dejanovic, B. et al. Changes in the Synaptic Proteome in Tauopathy and Rescue of Tau-Induced Synapse Loss by C1q Antibodies. Neuron 100, 1322–1336 e1327 (2018). 10.1016/j.neuron.2018.10.014

120 Dejanovic, B. et al. Complement C1q-dependent excitatory and inhibitory synapse elimination by astrocytes and microglia in Alzheimer’s disease mouse models. Nat Aging 2, 837–850 (2022). 10.1038/s43587-022-00281-1

121 Picard, M. & Shirihai, O. S. Mitochondrial signal transduction. Cell Metab 34, 1620–1653 (2022). 10.1016/j.cmet.2022.10.008

122 Murphy, M. P. & O’Neill, L. A. J. A break in mitochondrial endosymbiosis as a basis for inflammatory diseases. Nature 626, 271–279 (2024). 10.1038/s41586-023-06866-z

123 Al-Mehdi, A. B. et al. Perinuclear mitochondrial clustering creates an oxidant-rich nuclear domain required for hypoxia-induced transcription. Sci Signal 5, ra47 (2012). 10.1126/scisignal.2002712

124 Desai, R. et al. Mitochondria form contact sites with the nucleus to couple prosurvival retrograde response. Sci Adv 6 (2020). 10.1126/sciadv.abc9955

125 Sutandy, F. X. R., Gossner, I., Tascher, G. & Munch, C. A cytosolic surveillance mechanism activates the mitochondrial UPR. Nature 618, 849–854 (2023). 10.1038/s41586-023-06142-0

126 Lennicke, C. & Cocheme, H. M. Redox metabolism: ROS as specific molecular regulators of cell signaling and function. Mol Cell 81, 3691–3707 (2021). 10.1016/j.molcel.2021.08.018

127 Sies, H. et al. Defining roles of specific reactive oxygen species (ROS) in cell biology and physiology. Nat Rev Mol Cell Biol 23, 499–515 (2022). 10.1038/s41580-022-00456-z

128 Kumar, A. et al. HIF1alpha stabilization in hypoxia is not oxidant-initiated. Elife 10 (2021). 10.7554/eLife.72873

129 Herb, M. et al. Mitochondrial reactive oxygen species enable proinflammatory signaling through disulfide linkage of NEMO. Sci Signal 12 (2019). 10.1126/scisignal.aar5926

130 Kisty, E. A., Falco, J. A. & Weerapana, E. Redox proteomics combined with proximity labeling enables monitoring of localized cysteine oxidation in cells. Cell Chem Biol 30, 321–336 e326 (2023). 10.1016/j.chembiol.2023.02.006

131 Sobotta, M. C. et al. Peroxiredoxin-2 and STAT3 form a redox relay for H2O2 signaling. Nat Chem Biol 11, 64–70 (2015). 10.1038/nchembio.1695

132 Guo, X. et al. Mitochondrial stress is relayed to the cytosol by an OMA1-DELE1-HRI pathway. Nature 579, 427–432 (2020). 10.1038/s41586-020-2078-2

133 Jezek, P., Holendova, B. & Plecita-Hlavata, L. Redox Signaling from Mitochondria: Signal Propagation and Its Targets. Biomolecules 10 (2020). 10.3390/biom10010093

134 Yan, T. et al. Proximity-labeling chemoproteomics defines the subcellular cysteinome and inflammation-responsive mitochondrial redoxome. Cell Chem Biol 30, 811–827 e817 (2023). 10.1016/j.chembiol.2023.06.008

135 Topf, U. et al. Quantitative proteomics identifies redox switches for global translation modulation by mitochondrially produced reactive oxygen species. Nat Commun 9, 324 (2018). 10.1038/s41467-017-02694-8

136 Zhang, J., Frerman, F. E. & Kim, J. J. Structure of electron transfer flavoprotein-ubiquinone oxidoreductase and electron transfer to the mitochondrial ubiquinone pool. Proc Natl Acad Sci U S A 103, 16212–16217 (2006). 10.1073/pnas.0604567103

137 Missaglia, S., Tavian, D., Moro, L. & Angelini, C. Characterization of two ETFDH mutations in a novel case of riboflavin-responsive multiple acyl-CoA dehydrogenase deficiency. Lipids Health Dis 17, 254 (2018). 10.1186/s12944-018-0903-5

138 Zhuo, Z. et al. A case of late-onset riboflavin responsive multiple acyl-CoA dehydrogenase deficiency (MADD) with a novel mutation in ETFDH gene. J Neurol Sci 353, 84–86 (2015). 10.1016/j.jns.2015.04.011

139 Fan, X. et al. Novel ETFDH mutations in four cases of riboflavin responsive multiple acyl-CoA dehydrogenase deficiency. Mol Genet Metab Rep 16, 15–19 (2018). 10.1016/j.ymgmr.2018.05.007

140 Morant-Ferrando, B. et al. Fatty acid oxidation organizes mitochondrial supercomplexes to sustain astrocytic ROS and cognition. Nat Metab 5, 1290–1302 (2023). 10.1038/s42255-023-00835-6

141 Mi, Y. et al. Loss of fatty acid degradation by astrocytic mitochondria triggers neuroinflammation and neurodegeneration. Nat Metab 5, 445–465 (2023). 10.1038/s42255-023-00756-4

142 Ruprecht, B. et al. Chemoproteomic profiling to identify activity changes and functional inhibitors of DNA-binding proteins. Cell Chem Biol 29, 1639–1648 e1634 (2022). 10.1016/j.chembiol.2022.10.008

143 Parenti, M. D. et al. Discovery of the 4-aminopiperidine-based compound EM127 for the site-specific covalent inhibition of SMYD3. Eur J Med Chem 243, 114683 (2022). 10.1016/j.ejmech.2022.114683

144 Yang, K. et al. Accelerating multiplexed profiling of protein-ligand interactions: High-throughput plate-based reactive cysteine profiling with minimal input. Cell Chem Biol (2023). 10.1016/j.chembiol.2023.11.015

145 McHenry, M. W. et al. Covalent inhibition of pro-apoptotic BAX. Nat Chem Biol (2024). 10.1038/s41589-023-01537-6

146 Stanton, C. et al. Covalent Targeting As a Common Mechanism for Inhibiting NLRP3 Inflammasome Assembly. ACS Chem Biol 19, 254–265 (2024). 10.1021/acschembio.3c00330

147 Li, H. et al. Assigning functionality to cysteines by base editing of cancer dependency genes. Nat Chem Biol 19, 1320–1330 (2023). 10.1038/s41589-023-01428-w

148 Gurung, P., Lukens, J. R. & Kanneganti, T. D. Mitochondria: diversity in the regulation of the NLRP3 inflammasome. Trends Mol Med 21, 193–201 (2015). 10.1016/j.molmed.2014.11.008

149 Marchi, S., Guilbaud, E., Tait, S. W. G., Yamazaki, T. & Galluzzi, L. Mitochondrial control of inflammation. Nat Rev Immunol 23, 159–173 (2023). 10.1038/s41577-022-00760-x

150 Ising, C. et al. NLRP3 inflammasome activation drives tau pathology. Nature 575, 669–673 (2019). 10.1038/s41586-019-1769-z

151 Heneka, M. T. et al. NLRP3 is activated in Alzheimer’s disease and contributes to pathology in APP/PS1 mice. Nature 493, 674–678 (2013). 10.1038/nature11729

152 Couturier, J. et al. Activation of phagocytic activity in astrocytes by reduced expression of the inflammasome component ASC and its implication in a mouse model of Alzheimer disease. J Neuroinflammation 13, 20 (2016). 10.1186/s12974-016-0477-y

153 Shiratori-Hayashi, M. et al. STAT3-dependent reactive astrogliosis in the spinal dorsal horn underlies chronic itch. Nat Med 21, 927–931 (2015). 10.1038/nm.3912

154 Zhao, J. B., Zhang, Y., Li, G. Z., Su, X. F. & Hang, C. H. Activation of JAK2/STAT pathway in cerebral cortex after experimental traumatic brain injury of rats. Neurosci Lett 498, 147–152 (2011). 10.1016/j.neulet.2011.05.001

155 Oliva, A. A., Jr., Kang, Y., Sanchez-Molano, J., Furones, C. & Atkins, C. M. STAT3 signaling after traumatic brain injury. J Neurochem 120, 710–720 (2012). 10.1111/j.1471-4159.2011.07610.x

156 Priego, N. et al. STAT3 labels a subpopulation of reactive astrocytes required for brain metastasis. Nat Med 24, 1024–1035 (2018). 10.1038/s41591-018-0044-4

157 Shi, P. et al. Astrocyte-selective STAT3 knockdown rescues methamphetamine withdrawal- disrupted spatial memory in mice via restoring the astrocytic capacity of glutamate clearance in dCA1. Glia 69, 2404–2418 (2021). 10.1002/glia.24046

158 Abjean, L. et al. Reactive astrocytes promote proteostasis in Huntington’s disease through the JAK2- STAT3 pathway. Brain 146, 149–166 (2023). 10.1093/brain/awac068

159 Sarmiento Soto, M., et al. STAT3-Mediated Astrocyte Reactivity Associated with Brain Metastasis Contributes to Neurovascular Dysfunction. Cancer Res 80, 5642–5655 (2020). 10.1158/0008-5472.CAN-20-2251

160 O’Sullivan, S. A., O’Sullivan, C., Healy, L. M., Dev, K. K. & Sheridan, G. K. Sphingosine 1-phosphate receptors regulate TLR4-induced CXCL5 release from astrocytes and microglia. J Neurochem 144, 736–747 (2018). 10.1111/jnc.14313

161 Cao, Q. et al. Astrocytic CXCL5 hinders microglial phagocytosis of myelin debris and aggravates white matter injury in chronic cerebral ischemia. J Neuroinflammation 20, 105 (2023). 10.1186/s12974-023-02780-3

162 Wang, L. Y., Tu, Y. F., Lin, Y. C. & Huang, C. C. CXCL5 signaling is a shared pathway of neuroinflammation and blood-brain barrier injury contributing to white matter injury in the immature brain. J Neuroinflammation 13, 6 (2016). 10.1186/s12974-015-0474-6

163 Staurenghi, E. et al. Oxysterols present in Alzheimer’s disease brain induce synaptotoxicity by activating astrocytes: A major role for lipocalin-2. Redox Biol 39, 101837 (2021). 10.1016/j.redox.2020.101837

164 Jin, M., Jang, E. & Suk, K. Lipocalin-2 Acts as a Neuroinflammatogen in Lipopolysaccharide-injected Mice. Exp Neurobiol 23, 155–162 (2014). 10.5607/en.2014.23.2.155

165 Wang, L. et al. Nitric oxide mediates glial-induced neurodegeneration in Alexander disease. Nat Commun 6, 8966 (2015). 10.1038/ncomms9966

166 Zhang, J. et al. Systematic identification of anticancer drug targets reveals a nucleus-to- mitochondria ROS-sensing pathway. Cell 186, 2361–2379 e2325 (2023). 10.1016/j.cell.2023.04.026

167 Soberanes, S. et al. Metformin Targets Mitochondrial Electron Transport to Reduce Air-Pollution- Induced Thrombosis. Cell Metab 29, 335–347 e335 (2019). 10.1016/j.cmet.2018.09.019

168 Plecita-Hlavata, L. et al. Mitochondrial Superoxide Production Decreases on Glucose-Stimulated Insulin Secretion in Pancreatic beta Cells Due to Decreasing Mitochondrial Matrix NADH/NAD(+) Ratio. Antioxid Redox Signal 33, 789–815 (2020). 10.1089/ars.2019.7800

169 Morris, O., Deng, H., Tam, C. & Jasper, H. Warburg-like Metabolic Reprogramming in Aging Intestinal Stem Cells Contributes to Tissue Hyperplasia. Cell Rep 33, 108423 (2020). 10.1016/j.celrep.2020.108423

170 Kong, H. et al. Metabolic determinants of cellular fitness dependent on mitochondrial reactive oxygen species. Sci Adv 6 (2020). 10.1126/sciadv.abb7272

171 Homma, T., Kobayashi, S., Sato, H. & Fujii, J. Superoxide produced by mitochondrial complex III plays a pivotal role in the execution of ferroptosis induced by cysteine starvation. Arch Biochem Biophys 700, 108775 (2021). 10.1016/j.abb.2021.108775

172 Hatinguais, R. et al. Mitochondrial Reactive Oxygen Species Regulate Immune Responses of Macrophages to Aspergillus fumigatus. Front Immunol 12, 641495 (2021). 10.3389/fimmu.2021.641495

173 Lian, H. et al. NFkappaB-activated astroglial release of complement C3 compromises neuronal morphology and function associated with Alzheimer’s disease. Neuron 85, 101–115 (2015). 10.1016/j.neuron.2014.11.018

174 Mauro, C. et al. NF-kappaB controls energy homeostasis and metabolic adaptation by upregulating mitochondrial respiration. Nat Cell Biol 13, 1272–1279 (2011). 10.1038/ncb2324

175 Moretti, M., Bennett, J., Tornatore, L., Thotakura, A. K. & Franzoso, G. Cancer: NF-kappaB regulates energy metabolism. Int J Biochem Cell Biol 44, 2238–2243 (2012). 10.1016/j.biocel.2012.08.002

176 Nisr, R. B., Shah, D. S., Ganley, I. G. & Hundal, H. S. Proinflammatory NFkB signalling promotes mitochondrial dysfunction in skeletal muscle in response to cellular fuel overloading. Cell Mol Life Sci 76, 4887–4904 (2019). 10.1007/s00018-019-03148-8

177 Luongo, T. S. et al. The mitochondrial Na(+)/Ca(2+) exchanger is essential for Ca(2+) homeostasis and viability. Nature 545, 93–97 (2017). 10.1038/nature22082

178 Rottenberg, H., Covian, R. & Trumpower, B. L. Membrane potential greatly enhances superoxide generation by the cytochrome bc1 complex reconstituted into phospholipid vesicles. J Biol Chem 284, 19203–19210 (2009). 10.1074/jbc.M109.017376

179 Quinlan, C. L., Gerencser, A. A., Treberg, J. R. & Brand, M. D. The mechanism of superoxide production by the antimycin-inhibited mitochondrial Q-cycle. J Biol Chem 286, 31361–31372 (2011). 10.1074/jbc.M111.267898

180 Murphy, M. P. & Chouchani, E. T. Why succinate? Physiological regulation by a mitochondrial coenzyme Q sentinel. Nat Chem Biol 18, 461–469 (2022). 10.1038/s41589-022-01004-8

181 Galkin, A. et al. Identification of the mitochondrial ND3 subunit as a structural component involved in the active/deactive enzyme transition of respiratory complex I. J Biol Chem 283, 20907–20913 (2008). 10.1074/jbc.M803190200

182 Stine, W. B., Jungbauer, L., Yu, C. & LaDu, M. J. Preparing synthetic Abeta in different aggregation states. Methods Mol Biol 670, 13–32 (2011). 10.1007/978-1-60761-744-0_2

183 Yu, X. et al. Reducing Astrocyte Calcium Signaling In Vivo Alters Striatal Microcircuits and Causes Repetitive Behavior. Neuron 99, 1170–1187 e1179 (2018). 10.1016/j.neuron.2018.08.015

184 Dhandapani, P. K. et al. Hyperoxia but not AOX expression mitigates pathological cardiac remodeling in a mouse model of inflammatory cardiomyopathy. Sci Rep 9, 12741 (2019). 10.1038/s41598-019-49231-9

185 Sun, Y. et al. The behavioural and neuropathologic sexual dimorphism and absence of MIP-3alpha in tau P301S mouse model of Alzheimer’s disease. J Neuroinflammation 17, 72 (2020). 10.1186/s12974-020-01749-w

186 Orr, A. G., Orr, A. L., Li, X. J., Gross, R. E. & Traynelis, S. F. Adenosine A(2A) receptor mediates microglial process retraction. Nat Neurosci 12, 872–878 (2009). 10.1038/nn.2341

187 Motulsky, H. J. & Brown, R. E. Detecting outliers when fitting data with nonlinear regression - a new method based on robust nonlinear regression and the false discovery rate. BMC Bioinformatics 7, 123 (2006). 10.1186/1471-2105-7-123

188 Aguer, C. et al. Galactose enhances oxidative metabolism and reveals mitochondrial dysfunction in human primary muscle cells. PLoS One 6, e28536 (2011). 10.1371/journal.pone.0028536

189 Huang da, W., Sherman, B. T. & Lempicki, R. A. Systematic and integrative analysis of large gene lists using DAVID bioinformatics resources. Nat Protoc 4, 44–57 (2009). 10.1038/nprot.2008.211

190 Sherman, B. T. et al. DAVID: a web server for functional enrichment analysis and functional annotation of gene lists (2021 update). Nucleic Acids Res 50, W216–W221 (2022). 10.1093/nar/gkac194

191 Dobin, A. et al. STAR: ultrafast universal RNA-seq aligner. Bioinformatics 29, 15–21 (2013). 10.1093/bioinformatics/bts635

192 Trapnell, C. et al. Differential analysis of gene regulation at transcript resolution with RNA-seq. Nat Biotechnol 31, 46–53 (2013). 10.1038/nbt.2450

193 Trapnell, C. et al. Transcript assembly and quantification by RNA-Seq reveals unannotated transcripts and isoform switching during cell differentiation. Nat Biotechnol 28, 511–515 (2010). 10.1038/nbt.1621

194 Anders, S., Pyl, P. T. & Huber, W. HTSeq--a Python framework to work with high-throughput sequencing data. Bioinformatics 31, 166–169 (2015). 10.1093/bioinformatics/btu638

195 Love, M. I., Huber, W. & Anders, S. Moderated estimation of fold change and dispersion for RNA- seq data with DESeq2. Genome Biol 15, 550 (2014). 10.1186/s13059-014-0550-8

